# Comprehensive analyses of RNA-seq and genome-wide data point to enrichment of neuronal cell type subsets in neuropsychiatric disorders

**DOI:** 10.1101/2021.05.06.442982

**Authors:** M Olislagers, K Rademaker, RAH Adan, BD Lin, JJ Luykx

## Abstract

Neurological and psychiatric disorders, including substance use disorders share a range of symptoms, which could be the result of shared genetic background. Many genetic loci have been identified for these disorders using genome-wide association studies, but conclusive evidence about cell types wherein these loci are active is lacking. We aimed to uncover implicated brain cell types in neuropsychiatric traits and to assess consistency in results across RNA datasets and methods. We therefore comprehensively employed cell-type enrichment methods by integrating single-cell transcriptomic data from mouse brain regions with an unprecedented dataset of 42 human genome-wide association study results of neuropsychiatric, substance use and behavioral/quantitative brain-related traits (n=12,544,007 individuals). Single-cell transcriptomic datasets from the Karolinska Institute and the 10x Genomics dataset were used. Cell type enrichment was determined using Linkage Disequilibrium Score Regression, Multi-marker Analysis of GenoMic Annotation, and Data-driven Expression Prioritized Integration for Complex Traits. We found the largest degree of consistency across methods for implication of pyramidal cells in schizophrenia and cognitive performance. For other phenotypes, such as bipolar disorder, two methods implicated the same cell types, i.e. medium spiny neurons and pyramidal cells. For autism spectrum disorders and anorexia nervosa, no consistency in implicated cell types was observed across methods. We found no evidence for astrocytes being consistently implicated in neuropsychiatric traits. In conclusion, we provide comprehensive evidence for a subset of neuronal cell types being consistently implicated in several, but not all psychiatric disorders, while non-neuronal cell types seem less implicated.

## Introduction

It is well documented that several neuropsychiatric disorders, including substance use disorders (SUDs), share symptoms, which could be the result of shared genetic underpinnings^1, 2^. Much of the heritability (h^2^) of genetically complex or polygenic brain disorders -e.g. schizophrenia (SCZ), Parkinson’s disease and alcohol use disorder-is due to common genetic variation^3^. In addition, genome-wide association studies (GWASs) have deepened the understanding of such disorders, unravelling thousands of associated loci^4, 5^. However, elucidating disease mechanisms has remained challenging. One reason is missing heritability, meaning the gap between twin-based and single-nucleotide polymorphism (SNP)-based h^2^ estimates, which may result from limited statistical power, phenotypic heterogeneity, clinical misclassification, GWASs not probing associations with rare variants, epigenetics, genomic interactions, and structural genomic alterations^6^. Due to missing heritability, underlying causal genetic contributors might remain undetected. This impedes the translation of GWAS associations into their functional effects. Another reason explaining why elucidating disease mechanisms in neuropsychiatric disorders has remained challenging is that over 90% of identified variants are located within non-coding regions of the genome, indicating that regulatory elements -e.g. promoters, enhancers and insulators-may explain part of the underlying genetic mechanisms in some polygenic disorders^4, 7^. Due to extensive linkage disequilibrium (LD), it is also challenging to identify a causal variant within a given associated locus^4^.

To overcome gaps between associated and causal genetic association, their functional effects and ultimately the biological pathways, extensive research has been performed to identify brain tissues having a role in neuropsychiatric disease. Functional genomic studies using macroscopic brain samples point to enrichment in phylogenetically conserved areas of the brain in psychiatric disorders and brain-related behavioral phenotypes, whereas typically fewer brain regions are found to be enriched in neurological disorders^3^. However, identification of specific cell types within brain tissues is considerably less well studied. Specific cell types that are associated with SCZ and anorexia nervosa (AN) have previously been identified by integrating GWAS findings with mouse single-cell RNA (scRNA) brain data: while medium spiny neurons (MSNs), cortical interneurons, hippocampal CA1 pyramidal cells (pyramidal CA1), and pyramidal cells from the somatosensory cortex (pyramidal SS) seem implicated in SCZ^8^, suggestive findings were reported for enrichment of MSNs and pyramidal cells (CA1) in AN^9^. Recently, more extensive cell type enrichment analysis was performed for 28 phenotypes using mouse gene expression from the entire central nervous system (CNS)^10^. In psychiatric disorders, enrichment was found for MSNs, cortical interneurons, striatal interneurons, neuroblasts, pyramidal cells (CA1 and SS)^10^. In neurological disorders, fewer cell types were identified and these were dissimilar across disorders^10^. These cell type enrichment analyses have mainly been performed using Linkage Disequilibrium Score Regression (LDSC) and/or Multi-marker Analysis of GenoMic Annotation (MAGMA). However, in this landmark and other studies, MAGMA versions <1.08 have been employed^8-10^, of which it was recently reported that its SNP-level P-value aggregation into gene-level P-values might result in type-I errors^11^. In addition, cell type enrichment in SUDs and several other disorders, such as anxiety disorders, has to the best of our knowledge not been studied.

Here, we systemically investigated cell type enrichment in an extensive set of brain-related phenotypes by integrating mouse scRNA brain data from the Karolinska Institute (KI) and 10x Genomics with summary statistics from 42 phenotypes related to neuropsychiatry, SUDs, and brain-related behavior. Our goals were to perform cell type enrichment for a more comprehensive set of brain-related traits than previously studied and to assess consistency in results across a wider array of methods. We went beyond previous studies by systematically performing cell type enrichment analyses using the most recent releases of different methods that rely on different assumptions and algorithms, i.e. LDSC, MAGMA v1.08, Data-driven Expression Prioritized Integration for Complex Traits (DEPICT) and Functional Mapping and Annotation (FUMA). We found evidence for a subset of neuronal cell types being consistently implicated in several, but not all psychiatric disorders, while non-neuronal cell types seem less implicated.

## Methods

### GWAS summary statistics

Our goals were to identify salient cell types that are implicated in more brain-related traits than previously studied and to assess consistency in results across methods. Brain-related GWAS summary statistics from predominantly European samples were obtained from publicly available sources. A total of 41 summary statistics from brain-related phenotypes (Table 1, Table S1) were obtained, among which 11 psychiatric disorders (486,142 cases and 1,002,695 controls), 11 neurological disorders (186,171 cases and 2,278,970 controls), and 8 substance use disorders (case/control: 11,569 cases and 34,999; cohorts with continuous substance use data: n=3,683,037). All psychiatric and neurological disorders for which summary statistics were available had also been included in the Brainstorm project^3^. Substance use disorders were added because of the high comorbidity^12, 13^ and genetic covariance^14^ with psychiatric traits. Eleven well-powered (N>250,000) brain-related behavioral/quantitative phenotypes (n=4,166,895) were additionally selected. Because of the association between BMI and brain structure, we considered BMI a brain-related trait^15^. Finally, to discriminate cell types that were specific to the brain, height (n=693,529) was included as a non-brain-related anthropomorphic trait.

**Table 1.**
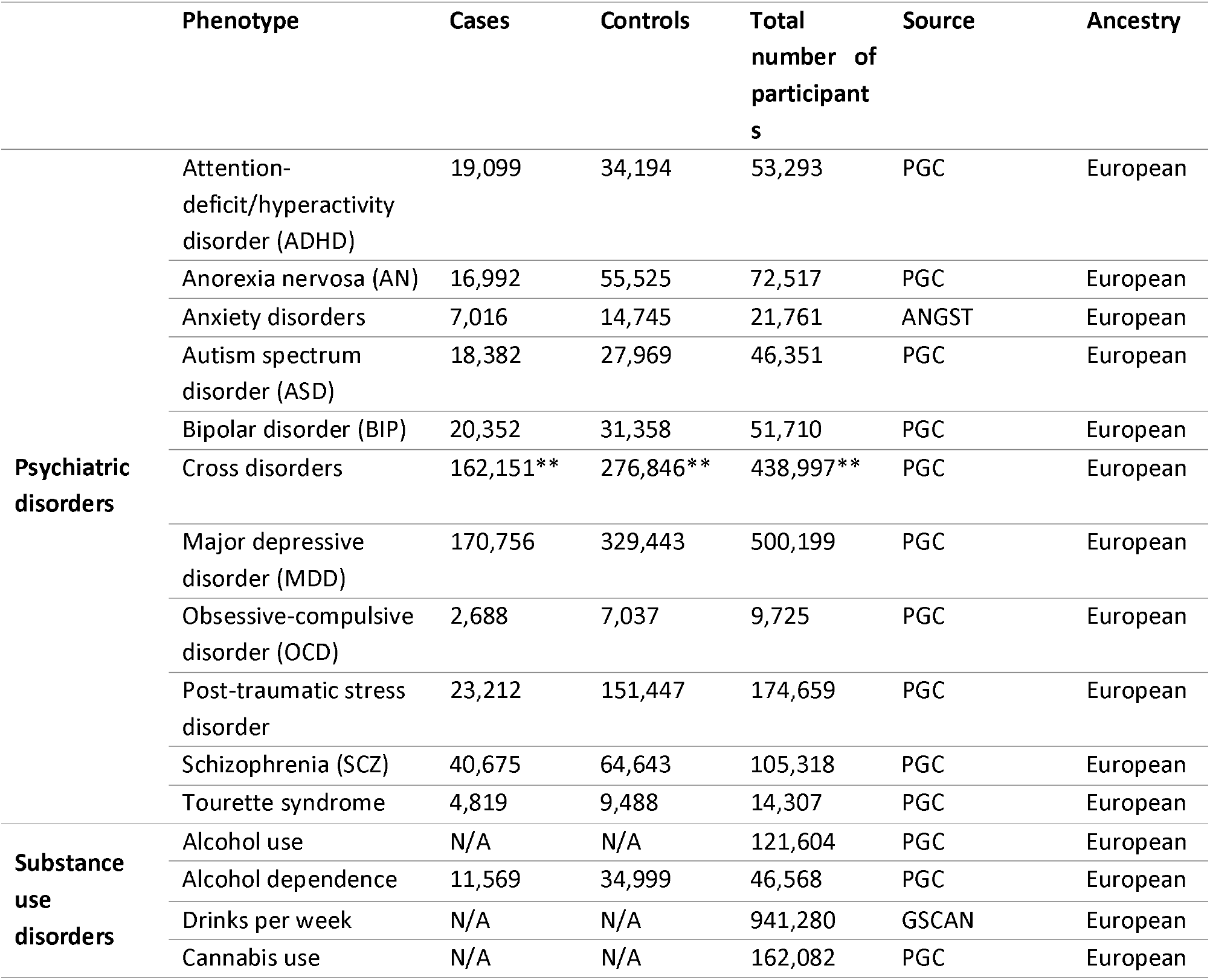
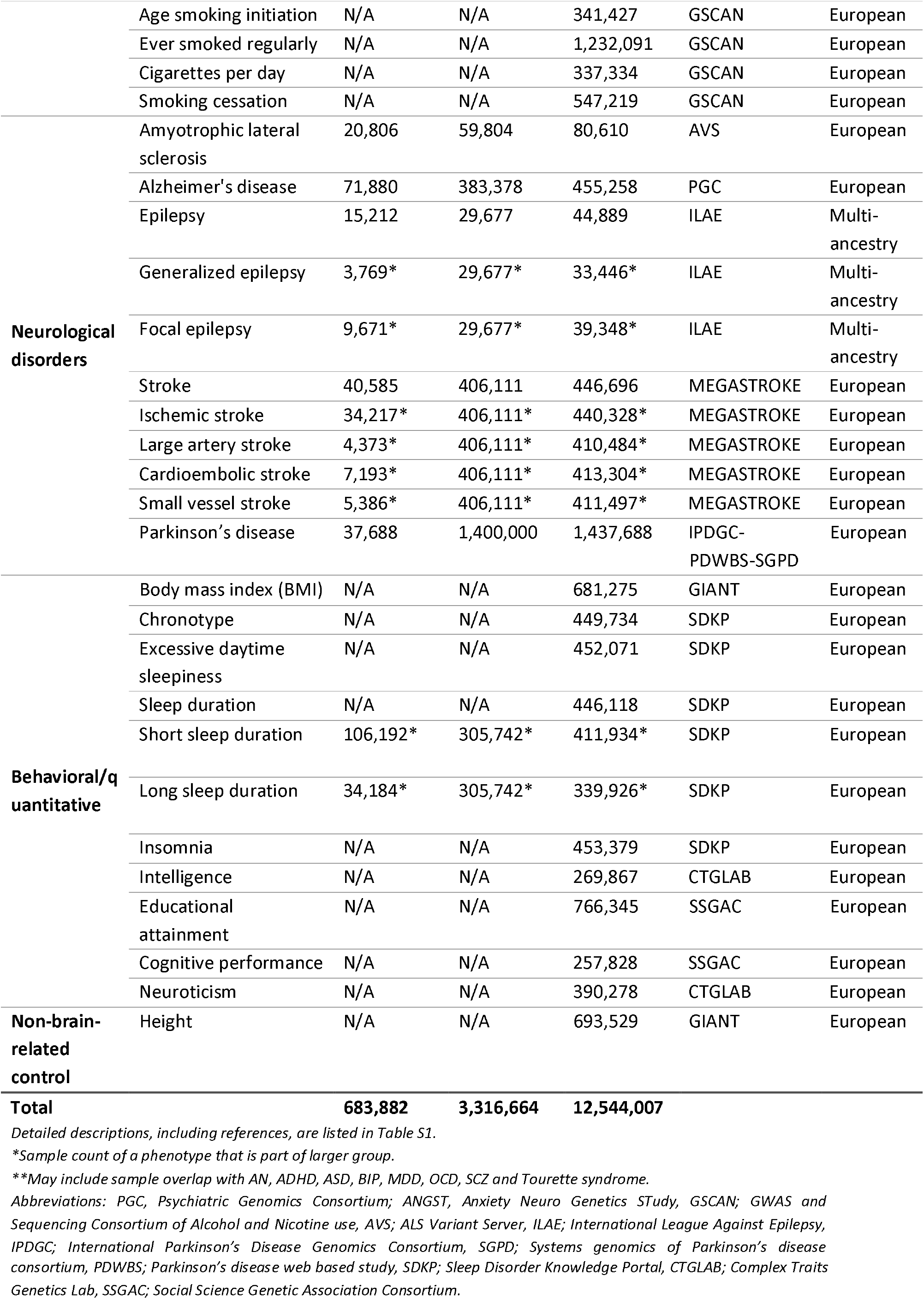
Phenotype descriptions.

### Single cell RNA sequencing datasets

All cell type enrichment analyses were conducted using the KI dataset^8, 16-18^ and the 10x Genomics dataset^19^. These datasets were selected because they cover brain regions that are generally accepted to be involved in the pathogenesis of brain-related disorders^20^. Additionally, their high coverage may enable the identification of different cell types. Detailed information about the 10x Genomics dataset, quality control, necessity of a randomized representative subset of cells, and cell type identification are reported in the Supplementary Methods. The quality control of the KI dataset is described elsewhere^16^.

### Overview of cell type enrichment analyses

To identify cell types underlying various phenotypes, we employed four methods (Figure 1). LDSC (version 1.0.1)^21, 22^ was first used to estimate SNP-h^2^ and bivariate genetic correlations across all traits. We then constructed the specificity metric Sg,c from the 10x Genomics dataset, denoting specificity of a gene g for cell type c, which was used as input for all three methods (Supplementary Methods). The specificity metric of the KI dataset was previously constructed^8^. Next, LDSC (version 1.0.1)^21, 23^, MAGMA (version 1.08)^8, 24^ and DEPICT (version 1, release 194)^25^ were employed to test cell type enrichment using the 10x Genomics and KI datasets. We utilized the diversity in statistical approaches of these methods to strengthen the confidence in consistently enriched cell types across methods. In brief, with LDSC we investigated the top 10% cell type-specific genes for enrichment of SNP-h^2^; with MAGMA we tested whether trait associations linearly increased with cell type-specificity or whether the top 10% cell type-specific genes were associated with gene-level association to traits; with DEPICT we evaluated the enrichment of cell type for genes from trait-associated loci.

**Figure 1.**
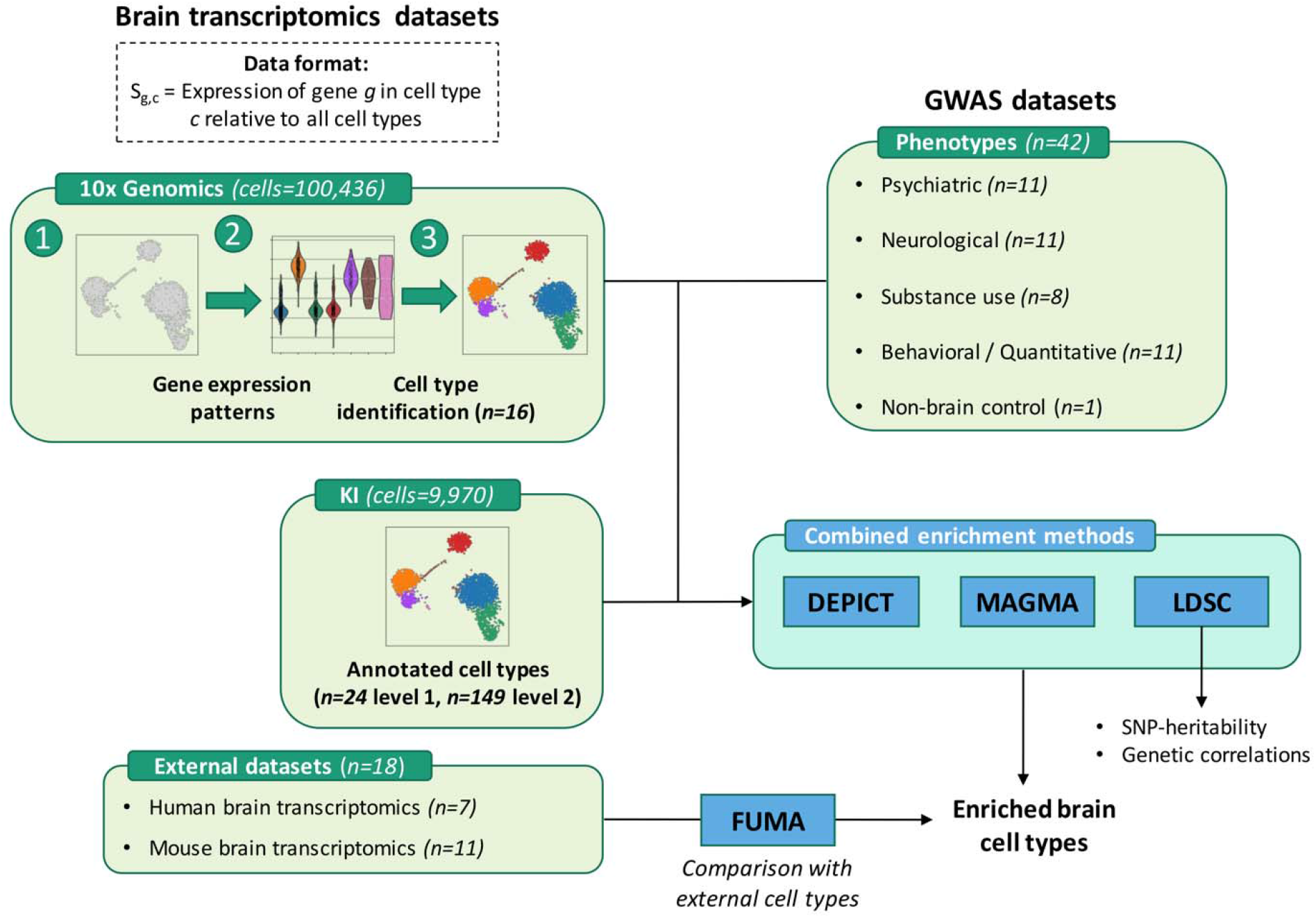
Overview of the approach of dataset integration as inputs for enrichment methods, in order to detect implicated brain cell types for various phenotypes. Two mouse brain transcriptomic datasets (10x Genomics, KI) have the data format Sg,c of cell-type specificity for genes, which was calculated by dividing expression of gene G in cell type C by expression of G in all cell types of a given dataset. Custom cell type identification was performed for 10x Genomics (16 detected cell types), while existing annotation was re-used for KI (first level of 24 cell types and second level of 149 cell (sub-)types). The datasets were integrated with genome-wide association study (GWAS) data, and these were the input for cell type enrichment methods DEPICT, MAGMA and LDSC. External human and mouse brain transcriptomics data were used in cell type enrichment method FUMA, so that enriched cell types from any of the other three methods could be compared to FUMA-enriched cell types. Finally, LDSC was also used to estimate SNP-based heritability for each GWAS phenotype and to calculate genetic correlations across all phenotypes.

LDSC computes an LD score by summarizing the correlations of a given SNP with all neighboring SNPs within 100 kb flanks. Then, the GWAS test statistic χ^2^ is regressed against the LD score, of which the slope is rescaled into an estimate of SNP-h^2^, explained by all SNPs included in the LD score. Based on Sg,c, we binned genes in specificity deciles and used LDSC to test for enrichment of SNP-h^2^ in the top 10% most associated genes. MAGMA aggregates χ^2^ association statistics within a 10 kb upstream and 1.5 kb downstream window into a gene-level P-value (Supplementary Methods). To compute gene-level P-values, Brown’s method^26^ was updated into Imhof’s approach^27^ in version 1.08^28^. DEPICT maps genes to loci by first selecting all significant SNPs and preserving lead SNPs as the most significant SNP out of possible SNP-pairs in LD (r^2^ > 0.1) and/or within < 1 Mb distance of each other (Supplementary Methods). Then, boundaries of the most distal SNP on either side around lead SNPs (r^2^ > 0.5) were used as criteria to list genes to SNPs, from which potential enrichment of these genes to specific cell types could later be assessed. By comparing cell type enrichment results of LDSC, DEPICT and MAGMA, we evaluated the relative stringency of each method. This was accomplished by comparing the P-values, denoting the strength of association of a given cell type with a given phenotype, of any two methods with one another.

Finally, additional human and mouse scRNA datasets were used to conduct additional cell type enrichment analysis, using FUMA (version 1.3.6a)^29^ (Supplementary Methods). FUMA builds on MAGMA (version 1.08). However, FUMA includes averaged expression per gene across cell type as covariate in their model instead of Sg,c and was therefore not included in our main analyses but as confirmational cell type enrichment analysis.

To allow for comparison between LDSC, MAGMA and DEPICT, we compared the P-values that refer to the strength of association of a given cell type with a given phenotype. KI level 1, KI level 2 and 10x Genomics cell types were identified as significant after passing a Bonferroni corrected significance level of P<0.05/(24*42), P<0.05/(149*42) and P<0.05/(16*42), respectively. We then counted the number of methods pointing to significant enrichment of specific cell types and report that number for both KI levels and 10x Genomics as our main outcome measure. Phenotypes implicating similar cell types were then identified by hierarchical clustering. We discuss these methods more elaborately below and in the Supplementary Methods.

### Cell type enrichment using LDSC

Human orthologs were obtained using the One2One R package that is incorporated in the MAGMA_Celltyping R package^8^. SNPs were annotated to the human genome (hg19, version 33) of the GENCODE project^30^. Binary annotations files were created for each cell type, containing 11 sub-annotations. The first sub-annotation contained SNPs that mapped to genes without a human ortholog (1 = SNP belongs to a sub-annotation). The other 10 sub-annotations represented the SNPs in specificity deciles for a particular cell type in increasing order. These specificity deciles were obtained by restructuring the specificity metric Sg,c, described in the Methods section ‘Overview of cell type enrichment analyses’ using the ‘prepare.quantile.groups’ function in the MAGMA_Celltyping package^8^. LD scores were then calculated for each annotation file using a 1 centimorgan (cM) window, 1000 Genomes Project Phase 3 files^31^ and restricted to 1,217,311 Hapmap3 SNPs. For each summary statistics dataset, we generated munged summary statistics by applying previously described quality control steps^22^ (Supplementary Methods), implemented in the LDSC ‘munge_sumstats.py’ script. Finally, SNP-h^2^ was partitioned, using the munged summary statistics, 1000 Genomes Project Phase 3 minor allele frequency files and both the 1000 Genomes Project phase 3 baseline model and all sub-annotations as independent variables. For the regression weights, we used the LD weights calculated for HapMap3 SNPs, excluding the major histocompatibility complex (MHC) region (chr6: 25-34 Mb) using the ‘overlap-annot’ to account for SNPs grouped into multiple deciles. In addition to the settings described above, we performed sensitivity analyses, including removing the HapMap3 SNPs restriction, using only SNPs that pass a genome-wide significance threshold, changing the software version and changing the reference genome version to determine differences in cell type enrichment results for SCZ using the KI dataset. To allow comparison of all enrichment methods, cell type enrichment figures show the P-value associated with the most specific decile for each cell type as not all methods provide an enrichment score. Methods for MAGMA, DEPICT and cell type enrichment analyses using additional mouse and human scRNA datasets are outlined in the Supplementary Methods.

## Results

### Cell type-specific gene expression in the 10x Genomics dataset

In the quality control of a randomized representative subset (n=108,844) of the 10x Genomics dataset, 8,408 cells and 6,419 non-expressed genes were removed from further analyses (Figure S1-S3). Altogether, the subset consisted of a matrix with 21,579 genes and 100,436 cells with 16 cell clusters (Table S3-S4). All cell clusters were subsequently mapped to brain cell types by specifically expressed marker genes (Figure S4).

### LDSC, MAGMA and DEPICT sensitivity analyses and quality control

For cell type enrichment analyses using LDSC, we initially adopted the same parameters that were previously described^8^. Additionally, to optimize the cell type enrichment pipeline we tested various settings (Figure S5, Table S5). The parameters described in the Methods section ‘Cell type enrichment using LDSC’ provided cell type enrichment results that were most consistent with DEPICT and MAGMA. For MAGMA, we found that restricting to HapMap3 SNPs and excluding the MHC region increased statistical power to identify associated cell types, whilst not inflating cell types that were not associated (Figure S6, Table S6). In addition to MAGMA version 1.08, we also performed cell type enrichment analyses using MAGMA version 1.07b (Table S7). We found that, although cell type associations follow similar patterns using both versions, the updated SNP-wise mean gene analysis model modestly exerts effects on cell type enrichment results, resulting in differently associated cell types (Figure S7). However, no consistent unidirectional differences in cell type enrichment results were observed. Finally, for DEPICT, we found that not restricting to HapMap3 SNPs increased statistical power to identify associated cell types, without an upwards bias for non-associated cell types (Figure S8, Table S8).

### Cell type enrichment analyses using the 10x Genomics dataset

Consistent with SNP-h^2^ estimate patterns (Table S9, Figure S9-10, Supplementary Methods, Supplementary Results) and genetic correlations (Table S10, Figure S11-12, Supplementary Methods, Supplementary Results), we found that cell type association patterns of neurological disorders were distinct from psychiatric, substance use and behavioral/quantitative association patterns by hierarchical clustering (Figure S13-14). For brain-related phenotypes, there were no cell types in the 10x Genomics dataset to which implicated genomic loci consistently mapped using all three methods (Figure S13-S15, Table S11). By comparing the P-values of each cell type computed by LDSC, MAGMA and DEPICT against one another, we found that MAGMA linear mode was considerably more lenient (Figure S16) and susceptible to bias due to sample size (Figure S17). Therefore, MAGMA linear mode was excluded from our main analyses.

Two methods provided evidence for certain neurons to be implicated in cross-disorders (8 psychiatric disorders jointly studied)^5^. A deeper cellular determination was not possible due to low sequencing depth. The same neuronal cells were also associated with educational attainment along with certain interneurons according to two methods. Additionally, these neurons were associated with intelligence. We also found evidence by two methods that implicated genomic loci of cognitive performance specifically mapped to certain neuroblasts. Suggestive findings are reported in the Supplementary Results.

### Cell type enrichment analyses using the KI dataset

After integrating GWAS findings with the 10x Genomics dataset, we leveraged cell type enrichment analyses by using the KI dataset, which is distinct from the 10x Genomics dataset because of its better coverage, higher resolution and higher specificity to identify cell type-specific gene expression markers. Therefore, considerably more cell types were identified on a deeper cellular level with the KI dataset. The largest degree of consistency across methods in brain-related traits was found for SCZ and cognitive performance (Figure 2 & 3, Figure S18-S19, Table S12). Genetic loci that are associated with SCZ consistently mapped to excitatory pyramidal cells (CA1 and SS), while those associated with cognitive performance only mapped to pyramidal cells (SS). For SCZ, we found evidence by two methods that MSNs were the main implicated inhibitory neurons. MSNs and both types of pyramidal cells were found to be associated with cross-disorders by two methods, while only pyramidal cells were associated with educational attainment and MSNs and pyramidal cells (CA1) were implicated in bipolar disorder. Suggestive findings are reported in the Supplementary Results.

**Figure 2.**
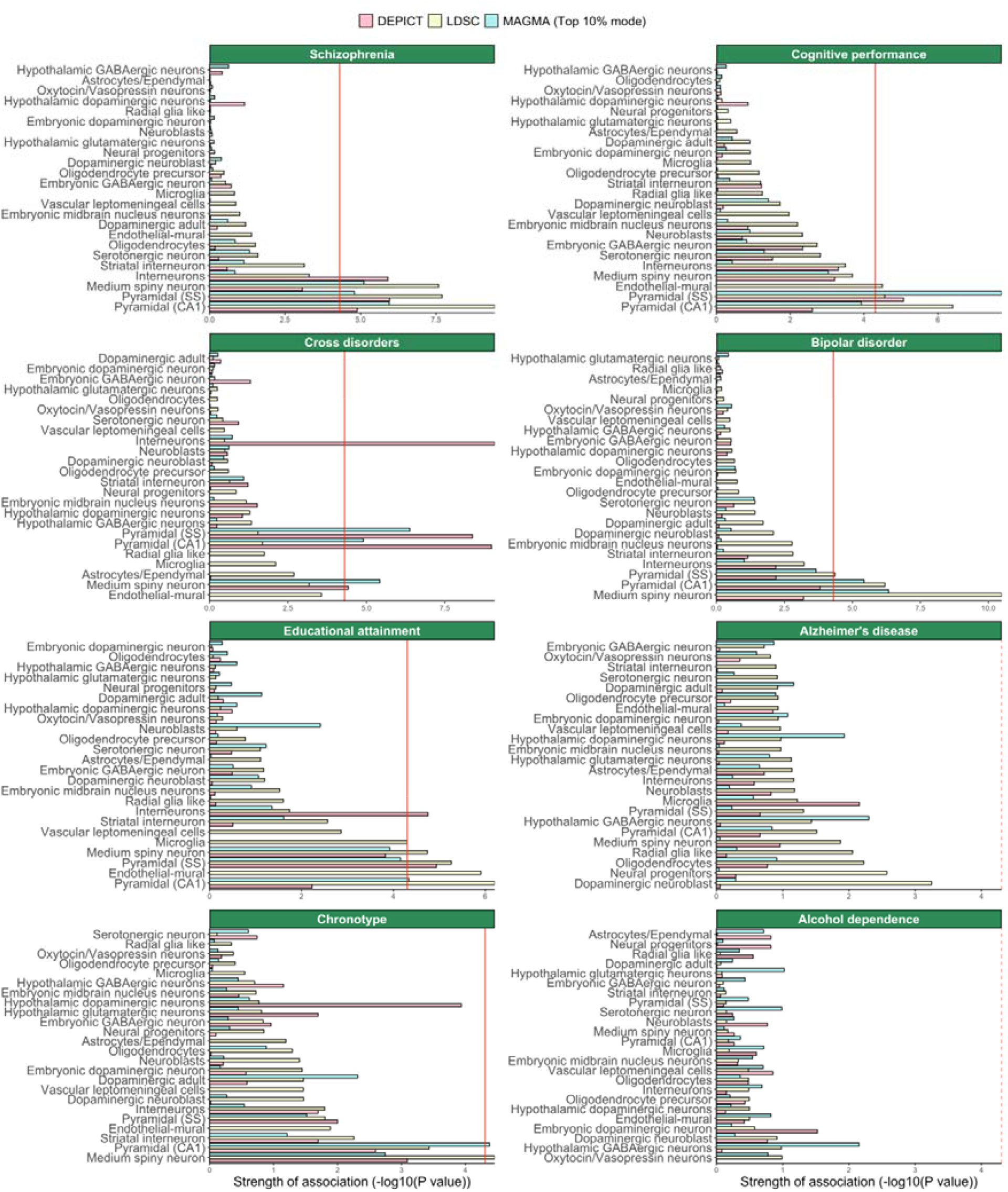
Cell type enrichment estimated by DEPICT, LDSC and MAGMA (top 10% mode) in selected brain-related phenotypes. Cell type enrichment results are generated using KI data. Bars represent the mean strength of association (-log10(P)) of LDSC, DEPICT and MAGMA (top 10%). The red line indicates the Bonferroni threshold P<0.05/(24*42). The red line is solid if any of the methods identified any cell type as significantly associated, and if none of the methods identified any of the cell types as significantly associated, the red line is dashed. A complete overview of cell type enrichment results using KI data, including MAGMA (linear) is available in the supplementary information (Figure S18).

**Figure 3.**
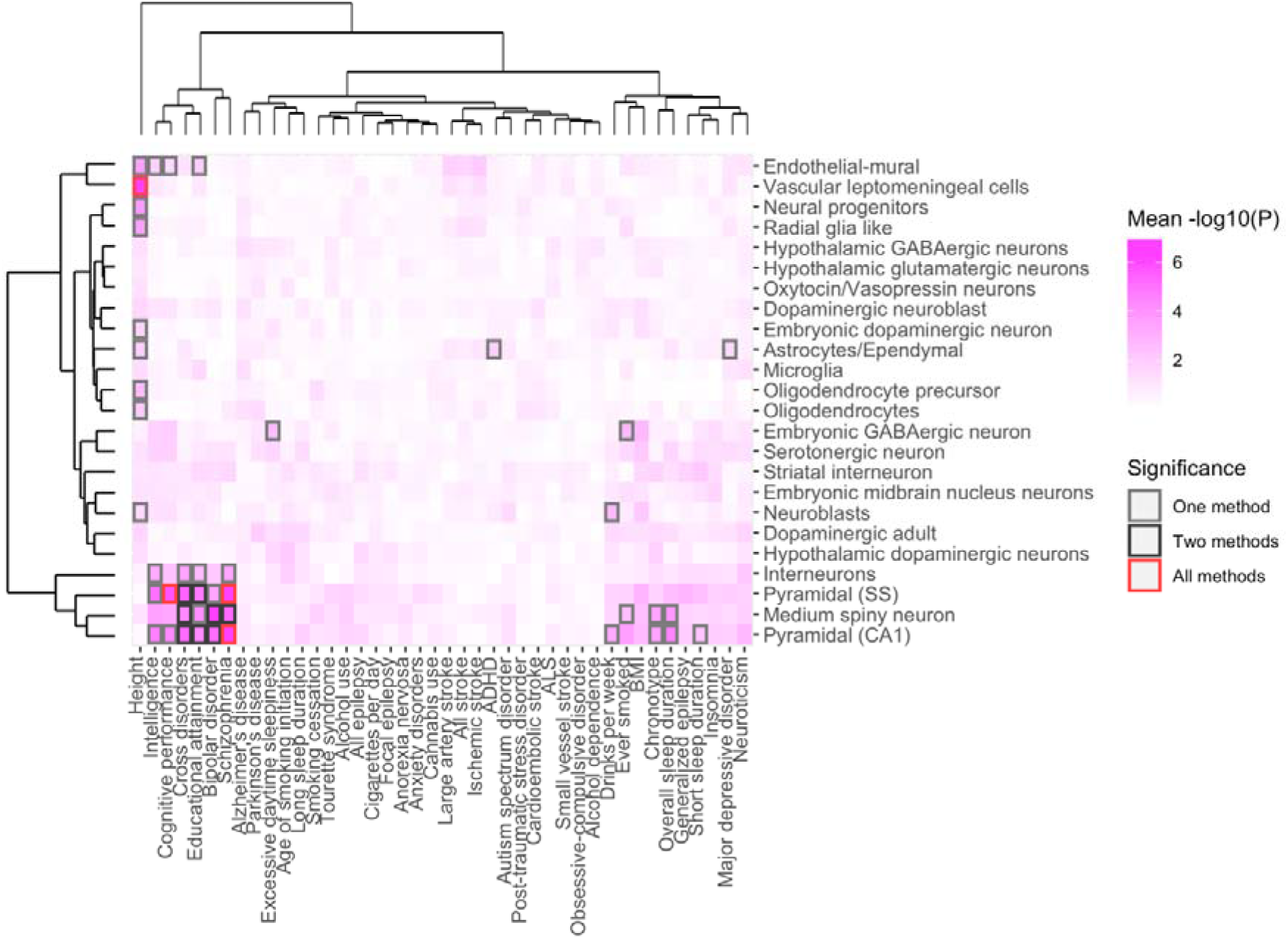
Overview of enriched cell types of 42 common-variant psychiatric, neurologic and behavioral/quantitative GWAS results in the KI dataset. Abbreviations: ADHD; attention deficit hyperactivity disorder, ALS; amyotrophic lateral sclerosis, BMI; body mass index. Analyses from LDSC, DEPICT and MAGMA (top 10% mode), referred to as ‘methods’ in the graph, show enrichment in MSNs and pyramidal cells (CA1) and pyramidal cells (SS) across brain-related phenotypes. The largest degree of consistency was found in SCZ and cognitive performance. Phenotypes and cell types are grouped by hierarchal clustering Shades of pink are proportional to the mean strength of association (-log10(P)) of all methods. The color of the frames refers to the number of methods that identified a given cell type as significant in a given phenotype, after Bonferroni correction (P<0.05/(24*42)). Grey frames: one method (intelligence, excessive daytime sleepiness, ADHD, drinks per week, ever smoked, chronotype, overall sleep duration, short sleep duration, MDD). Black frames: two methods (cross-disorders, educational attainment, BIP). Red frames: all three methods (human height, cognitive performance, SCZ).

To identify more differentiated cell types, the analysis was expanded by using the KI level 2 dataset (Figure S20-S22, Table S13), which includes 149 cell types that were subtypes of the cell types identified in the level 1 dataset. Using both the KI level 1 and level 2 datasets, we again found that MAGMA linear mode, compared to strength of association estimates computed by LDSC, MAGMA top 10% mode and DEPICT, provided disproportionally large estimates (Figure S23-S24) and was prone to inflated results due to sample size (Figure S25).

### Cell type enrichment analyses using additional scRNA datasets

Finally, we performed additional analyses with FUMA using additional mouse (n=11) and human (n=7) gene expression datasets to compare our findings and to assess consistency between rodent and human data. We were able to identify at least one implicated cell type in 22 phenotypes (Figure S26, Table S14). Using human gene expression data, 24 cell types were enriched in at least one phenotype, while using mouse scRNA data 70 cell types were revealed. Consistent with findings from 10x Genomics and KI datasets, pyramidal cells from various mouse brain regions, among which pyramidal cells (CA1 and SS), were implicated in SCZ and cognitive performance. Pyramidal cells were also enriched in numerous psychiatric disorders, SUDs and behavioral/quantitative phenotypes. Along with pyramidal cells, inhibitory GABAergic and MSNs were consistently enriched in psychiatric disorders, SUDs and behavioral/quantitative phenotypes. Using human datasets, enrichment of pyramidal cells (CA1) was confirmed in SCZ, cognitive performance, intelligence, cross-disorders, and overall sleep duration. Additionally, pyramidal cells (CA1) were enriched in cigarettes smoked per day. However, the strongest consistent evidence was found for enrichment of GABAergic neurons from the prefrontal cortex and midbrain in various psychiatric disorders, SUDs and behavioral/quantitative phenotypes. Consistent with findings using 10x Genomics and KI data, fewer enriched cell types were identified in neurological disorders. Human microglia were enriched in Alzheimer’s disease and human inhibitory GABAergic neurons from the prefrontal cortex and midbrain in generalized epilepsy. Generalized epilepsy was the only neurological phenotype in which mouse cell types were identified, namely certain pyramidal neurons and certain inhibitory neurons. We thus largely confirmed the main cell type enrichment findings from the 10x Genomics and KI dataset in mouse datasets and in human datasets using FUMA.

## Discussion

Here, we provide a comprehensive overview of specific brain cell types implicated in a range of brain-related phenotypes using both mouse and human brain scRNA data. We show that results from brain-related GWAS data consistently map to excitatory pyramidal neurons (CA1 and SS) and inhibitory MSNs and less so to glial and embryonic cells. The largest degree of consistency across methods and tissue origins (rodent and human) was found for implication of pyramidal cells in schizophrenia and cognitive performance. In contrast, GWAS data from neurological disorders mapped to fewer cell types that were distinct from psychiatric and substance use disorders.

Our SNP-h^2^ and genetic correlation findings confirm that neurological disorders are genetically distinct from one another and from psychiatric disorders and SUDs, as well as from behavioral/quantitative phenotypes, which is in line with previous evidence^3, 10^. Consistent with these findings, we found that GWAS findings from psychiatric disorders, SUDs and brain-related behavioral/quantitative phenotypes, but not neurological disorders, consistently map to excitatory hippocampal pyramidal neurons (CA1), excitatory pyramidal neurons (SS) and inhibitory MSNs and much less to glial and embryonic cells. This concurs with previous lines of evidence pointing to neurological disorders being genetically and functionally distinct from psychiatric disorders, SUDs and brain-related behavioral/quantitative phenotypes^3, 10^. Alzheimer’s disease was the only malady targeted here that showed evidence of exclusively human glial cells being implicated, underscoring the importance of key transcriptomic differences between human and mouse microglial signatures^33^. We confirmed our main findings with multiple external scRNA datasets using FUMA. This provides further evidence that genetic underpinnings of neurological disorders are distinct from those of psychiatric, SUDs and behavioral/quantitative phenotypes^8, 10^. Therefore, our main findings were based on the identification of cell types by LDSC, DEPICT and MAGMA top 10% mode. MAGMA linear mode was omitted because its strength of association estimates were consistently deviating substantially from LDSC, DEPICT and MAGMA top 10% mode. Therefore, it was deemed too lenient and thus prone to type I error inflation. This concurs with previous studies reporting that binned MAGMA analyses in linear mode inflated results since the binned scores can have strong correlations with the average gene expression across cell types^29^. Also in agreement with previous lines of evidence, we confirm that the statistical foundation of the SNP-wise mean gene analysis model MAGMA <1.07 may result in biased associations of cell types^11^.

We envision three broad implications for psychiatry as a medical specialty. First, our results consistently point to specific neuronal cell types being implicated in several psychiatric disorders. Therapeutic targeting of those cells could one day result in innovative treatments. Second, gene sets that are specifically enriched in those cells (for instance hippocampal pyramidal cells) could be used for risk scoring to differentiate patient subgroups and tailor therapy. Third, for the disorders with less consistent results, clinicians and researchers should aim to collect more samples and thus ensure future studies may shed light on their implicated cell types.

The discrepancy between the KI- and 10x Genomics-derived cell types could be a consequence of a lower sequencing depth in the 10x Genomics dataset (approximately 18,500 mapped reads per cell) than in the KI dataset (approximately 1.2 million mapped reads per cell). Notably, the minimum sequencing depth is generally considered to be between 25,000 and 50,000 mapped reads per cell^34^. This suggests that the relatively low sequencing depth of the 10x Genomics dataset led to overlapping cell clusters. Additionally, although k-means clustering is commonly used for single cell data, selecting the right value of k is challenging^34^. PCA-based clustering methods would be particularly well-suited for low sequencing depth^35^, and for instance could be expanded to initially select significant principal components with PCA and use these for subsequent clustering^36^.

Although we provide new insight with the largest and most comprehensive study of cell type enrichment in brain-related disorders, our results should be interpreted in light of inevitable limitations. First, using MAGMA, it is possible to test whether the genes specific to a phenotype are enriched in genetic associations of that phenotype while controlling for genetic associations of another phenotype^10^. However, as our main goal was to identify enriched cell types, such conditional analyses are beyond the scope of this study. Second, we found that microglia associated with age-induced neuroinflammation were exclusively found to be enriched in Alzheimer’s disease using human scRNA datasets, whereas no enriched glial cells were identified using mouse scRNA datasets. Therefore, mouse gene expression data from not only a spatial, but also a temporal resolution is warranted for future research to identify cell types implicated in disease during development. Additionally, improved coverage of brain-related regions, such as the entire CNS^10^, is warranted for future research. Third, a potential limitation of the FUMA model is that averaging gene expression disregards that low expression levels of certain genes can still be relevant to disease. This caveat illustrates the challenges of accounting for factors potentially inflating statistical results as well as capturing etiological mechanisms by examining cell type-specific gene expression^10^. Finally, to identify enriched cell types, we integrated human genomic findings with mouse scRNA brain expression data. Although considerable differences exist between mice and humans, we believe our choice is justified because mouse scRNA datasets cover transcripts that are missed in human single-nuclei sequencing^8^; cover more brain regions that are believed to be important in neuropsychiatric disorders than in human, such as the striatum for which no human scRNA expression data is available; and reveal key findings consistent with human data^8, 9^. In addition, gene expression data from rodents are often of higher quality, as fresh tissue can be more readily obtained and gene expression data cluster by cell type in different species, rather than by different cell types in the same species^8^. Although we believe that mouse scRNA data are suitable to apply to human GWAS data, there are limitations, e.g. less conserved brain regions might contain cell types that express genes differently; cell types could be specific to certain species (on the other hand, it has been shown that gene expression in the brain, including key gene expression patterns, is well conserved across species^32^); and cell types could have different functions or could be connected to and active in different brain circuits.

The identification of a specific subset of brain cell types being implicated in various brain disorders only marks the beginning of elucidating causal biological pathways. One question future research should address is what the effects of genetic variants in the non-coding genome are. One way to address this question is using an activity-by-contact model^37^. This model allows for the identification of cell type-specific enhancers and their target genes by leveraging single-cell chromatin accessibility and enhancer activity data. Additional insight could be obtained by performing cell prioritization analyses from human post-mortem brain samples and/or induced pluripotent stem cells from individuals with relevant genetic backgrounds using LDSC, MAGMA and DEPICT to identify genes that are predicted to be functionally similar to causal genes. Importantly, enriched cell types are not necessarily causal, but might be part of a neural network. To confirm our findings and to elucidate causality, selective chemo- and optogenetic manipulation of identified enriched cell types in rodents might provide additional insight in the role these cells play in the neural circuit underlying brain disorders^38^. Additionally, the recently developed computational toolkit CELL-type Expression-specific integration for Complex Traits (CELLECT) can provide additional insight in cell type enrichment^39^. CELLECT builds upon gene prioritization models, such as LDSC, DEPICT and MAGMA and subsequently performs cell type prioritization analyses using a continuous representation of cell type expression, rather than binary representation. Finally, statistical power is currently a major challenge in genetic studies. Future studies might benefit from multi-trait analysis of GWAS (MTAG)^40^, which is a method for analysis of multiple GWASs, thereby increasing the statistical power of each trait analyzed to identify genetic associations.

In sum, by incorporating different tools that rely on different assumptions and algorithms we provide robust evidence for a subgroup of neuronal cell types consistently implicated in several brain-related phenotypes. We thus provide a framework that furthers the understanding of cell types involved in brain-related phenotypes at a cellular level that can serve as a basis for future, more hypothesis-driven research.

## Data availability

All scRNA datasets used in this study are publicly available. All summary statistics are publicly available and the sources are listed in Table S1. We have made our code publicly available at https://github.com/mitchellolislagers/cell_type_enrichment_pipeline so that with the advent of new GWASs researchers may readily apply our pipeline to new data.

## Acknowledgements

Funding was provided through a personal Rudolf Magnus Talent Fellowship (H150) to Jurjen Luykx and by grants of the Netherlands Organisation for Scientific Research (NWO: 024.004.012, ALWOP.137 and OCENW.KLEIN.071) to Roger Adan.

The authors thank Kevin Kenna, Mark Bakker and Nathan Skene for fruitful input on statistical analyses. The authors thank Albert Batalla Cases for comments and suggestions on the manuscript.

## Disclosure

All authors declare they have no conflict of interest.

## Notes

### Competing Interest Statement

The authors have declared no competing interest.

### Summary of Updates

Added statistical backgrounds of assessed cell type enrichment methods; Supplemental files updated; Revised readability.

