## Supplementary Figures and Tables 1-4 for "Comprehensive analyses of RNA-seq and genome-wide data point to enrichment of neuronal cell type subsets in neuropsychiatric disorders"

^#^ Shared supervision.

**Keywords:** Cell type enrichment; GWAS; single-cell RNA sequencing; LDSC; MAGMA; DEPICT; FUMA

### **Supplementary Methods**

#### SNP-heritability estimates and bivariate genetic correlations

Linkage Disequilibrium Score Regression (LDSC) (version 1.0.1)^1^ was used to estimate single-nucleotide polymorphism-based heritability (SNP-h^2^), bivariate genetic correlations^2^ and cell type enrichment^3, 4^. To estimate SNP-h^2^ and genetic bivariate correlations, summary statistics were prepared using the LDSC munge_sumstats.py script. SNP-h^2^ was estimated by regressing the association *χ*^2^ statistic of a given SNP against the total amount of genetic variation that was tagged by that SNP. Bivariate genetic correlations across all phenotypes were estimated by regressing the product of z-scores against the total amount of genetic variation. The significance level was Bonferroni corrected, resulting in a threshold of p<0.05/861, thus correcting for 861 bivariate tests performed (42*(42-1)/2). All LDSC analyses were restricted to 1,217,311 Hapmap3 SNPs^5^. In case additional data was provided, further quality control was based on sample size (N>N.quantile(0.9)/1.5), INFO score (INFO>0.9) and minor allele frequency (minor allele frequency (MAF)>0.01). SNP-h^2^ and genetic bivariate correlations analyses were conducted using the --h2 and --rg flag, respectively, in the ldsc.py script, using the 1000 Genomes Project phase 3 baseline model and regression weights for Hapmap3 SNPs^6^. The major histocompatibility complex (MHC) region (chr6: 25-34 Mb) was excluded due to its extraordinary LD structure and gene density^7^.

#### Quality control and preparation of single cell RNA datasets

The Karolinska Institute (KI) dataset covers the neocortex^8^, hippocampus^8^, hypothalamus^9^, striatum and midbrain^10^, is composed of 9,970 cells, includes 24 identified brain cell types and was published including quality control^8^. The 10x Genomics dataset covers the cortex, hippocampus and subventricular zones, is composed of 1,306,127 cells, and was published without cell type annotation yet including cell clustering results (k-means clustering, ranging from k=2 to k=20)^11^.

Accessing the 10x Genomics dataset with Python instructions from the authors posed too large a computational burden, so that we generated subsets of ½, ¼, 1/8 and 1/12 the original size by randomly selecting cells. This approach was motivated by previous research which indicated that minimally 50,000 randomly chosen cells from this dataset could represent most biologically interpretable cell types^12^. The multiple cell clustering results were each projected on the full dataset in t-distributed stochastic neighbor embedding (t-SNE) plots in order to visually inspect which gave the most separation of cell clusters. Furthermore, cell cluster compositions were compared between the full datasets and subsets to check for subset-specific biases towards cell clusters. Ultimately, the choice for a 1/12th subset (n=108,844 cells) and k-means (k=20) clustering was made on the grounds of having comparatively well-distinguishable cell clusters, the absence of subset-specific biases in cell clusters, not suffering from computational burden and surpassing the previously suggested minimum of 50,000 cells.

SCANPY^13^, a Python toolkit for single-cell transcriptomics, was used for quality control and marker gene detection in the 10x Genomics subset. Cell quality control had the criteria: (1) UMIs per cell within μ±3σ, (2) genes per cell between 1000 and μ+3σ, (3) % mitochondrial DNA below μ+3σ. Distributions of these criteria were calculated; UMIs (4908 ± 3019), genes (2015 ± 777) and % mitochondrial DNA (3.39 ± 1.80) (Figure S1). Gene quality control consisted of removing non-expressed genes. Finally, cell clusters that would fail quality control or remain with low numbers of cells were removed. The identification of cell types in the 10x Genomics subset began with compiling a list of marker genes from literature^8, 11, 12, 14, 15^(Figure S2, Table S3) and the companion wiki from Zeisel et al.^8^ Subsequently, the SCANPY function ‘rank_genes_groups’ was implemented and used a Wilcoxon rank-sum test with Benjamini-Hochberg correction to identify top-ranked genes for cell cluster specificity. Finally, cell type identities were manually assigned by matching the detected marker genes to those in literature and those annotated to specific cell types in the companion wiki.

To identify cell type-specificity of genes, gene expression data was restructured into the specificity metric S_g,c_ (specificity of gene *g* for cell type *c*), by dividing the expression of *g* in *c* by the expression of *g* in all cell types. This metric was calculated for the 10x Genomics subset and was already publicly available for the KI dataset^4^.

#### Cell type enrichment using MAGMA

Multi-marker Analysis of GenoMic Annotation (MAGMA)^4, 16^ version 1.08 was used to identify either whether gene-level association of summary statistics linearly increased with cell type expression specificity or whether the top 10% specific gene-level association of the summary statistics were associated with cell type expression specificity. The analysis was restricted to Hapmap3 SNPs and the MHC region was excluded. The KI and 10x Genomics specificity metric Sg,c were transformed into 41 bins using the ‘prepare.quantile.groups’ from the MAGMA_Celltyping R package. Mouse genes were then mapped to human orthologs. A 10kb upstream and 1.5kb downstream window was used to include SNPs surrounding GWAS hits to compute single gene-level P-values. The association tests were performed with the ‘calculate_celltype_associations’ function in linear and top 10% mode. The top 10% mode is more specific as it only takes into account the top 10% most specific gene-level P-values, whereas the linear mode provides more power by taking into account gene-level P-values in all binned fractions in a linear regression model. As a sensitivity analysis, MAGMA was also run unrestricted to HapMap3 SNPs and including the MHC region to determine their effects on cell type enrichment results for schizophrenia (SCZ) using both the KI and 10x Genomics dataset. The analyses were additionally performed in MAGMA (version 1.07b) to detect the implications of the recently corrected statistical foundation in the ‘snp-wise mean model’ of version 1.07b that is used to aggregate SNP-level P-values into a gene-level test statistic^17^.

#### Cell type enrichment using DEPICT

Data-driven Expression Prioritized Integration for Complex Traits (DEPICT)^18^ (version 1, release 194) was used to identify cell types wherein genes from associated loci were significantly enriched using the specificity metric­ Sg,c. Only SNPs that passed a significance threshold (P<1x10^-5^) were included in the analysis. As DEPICT does not allow more than 1,000 associated SNPs to be included in the analysis, a stricter significance threshold was applied for body mass index (BMI) (P<5x10^-8^), educational attainment (P<5x10^-8^) and height (P<1x10^-15^) to reduce the number of associated SNPs that pass the significance threshold. Default parameters were used for the remaining settings. To evaluate the effect of restricting to HapMap3 SNPs on KI-derived cell type enrichment, DEPICT was run both restricted and unrestricted to HapMap3 SNPs for SCZ as a sensitivity analysis. Additionally, the MHC region was excluded.

#### Additional cell type enrichment analyses using additional mouse and human scRNA datasets

To confirm our cell type enrichment findings and to investigate whether these findings could be replicated using human data, additional cell type specificity analyses were performed using Functional Mapping and Annotation (FUMA)^19, 20^ (version 1.3.6a), which applies MAGMA (version 1.08) to test for positive relationships between cell type expression specificity of various external mouse (n=11) and human (n=7) scRNA datasets (Table S2) and associated SNPs that are aggregated to a gene-level P-value. The scRNA datasets originate from various regions of the brain, such as cortices (cerebral, frontal, somatosensory), basal ganglia (striatum, substantia nigra, globus pallidus), hippocampus, cerebellum and hindbrain. SNPs that passed the genome-wide significance threshold (P<5x10^-8^) were included in the analyses. Because no SNPs were independently associated with obsessive-compulsive disorder (OCD) and small vessel stroke, a lenient significance threshold (P<1x10^-5^) was applied for those phenotypes. Conditional pair-wise analyses were applied both per dataset and across datasets to identify independently associated cell types.

### **Supplementary Results**

#### SNP-heritability

Among all phenotypes, low to modest SNP-h^2^ were observed (Figure S9, Table S9). Psychiatric disorders were generally moderately heritable, with SCZ being the most heritable psychiatric disorder (SNP-h^2^(standard error (SE))=0.41(0.01) and post-traumatic stress disorder (PTSD) the least heritable (SNP-h^2^(SE)=0.02(0.00)). Contrastingly, substance-use disorders (SUDs) traits were generally less heritable, with the highest SNP-h^2^ estimate for alcohol use (SNP-h^2^(SE)=0.09(0.01)) and the lowest SNP-h^2^ estimate for smoking cessation (SNP-h^2^(SE)=0.03(0.00)). Of neurological disorders, generalized epilepsy was the most heritable (SNP-h^2^(SE)=0.27(0.03)) phenotype and large artery stroke was the least heritable (SNP-h^2^(SE)=0.00(0.00)). Finally, height was the most heritable behavioral/quantitative phenotype (SNP-h^2^(SE)=0.46(0.02)) that was assessed and long sleep duration was the least heritable behavioral/quantitative phenotype (SNP-h^2^(SE)=0.03(0.00)). No inflation of SNP-h^2^ estimation was observed with phenotypes containing a larger sample size (r=-0.17, P=0.27) (Figure S10). Generally, it can be considered that psychiatric and behavioral/quantitative phenotypes were relatively heritable whereas neurological and SUDs were typically less heritable.

#### Bivariate genetic correlations across psychiatric disorders

Widespread genetic correlations were observed across psychiatric disorders (Figure S11-S12, Table S10). Cross disorders, which is a meta-analysis of the eight psychiatric disorders anorexia nervosa (AN), attention-deficit/hyperactivity disorder (ADHD), autism spectrum disorder (ASD), bipolar disorder (BIP), major depressive disorder (MDD), OCD, SCZ, and Tourette syndrome (TS), was genetically correlated with most of the psychiatric disorders, but only nominally with ADHD (genetic correlation (r_g_) = 0.88, P=3.0x10^-3^), OCD (r_g_=0.32, P=3.8x10^-4^) and Tourette syndrome (r_g_=0.41, P=2.6x10^-4^). SCZ revealed genetic links with most of the other psychiatric disorders, but not ADHD, AN, anxiety disorders, or TS. In addition, MDD was highly correlated to most of the psychiatric disorders, except for ADHD, anxiety disorders and TS, whereas BIP was only genetically linked with SCZ and MDD. However, non-significant correlations between BIP, as well as OCD, and the other psychiatric traits became apparent. ADHD, AN and anxiety disorders were genetically less correlated with other psychiatric disorders.

#### Bivariate genetic correlations across neurological disorders

In contrast to the psychiatric disorders, genetic correlations were estimated to be small across neurological disorders (Figure S11-S12, Table S10). Significant genetic links were only observed across subtypes of disorders (i.e. epilepsy and stroke). Amyotrophic lateral sclerosis (ALS) was genetically positively correlated to some extend with Alzheimer’s disease (r_g_=0.40, P=6.5x10^-3^) and negatively correlated with cardioembolic stroke (r_g_=-0.33, P=0.04) and large artery stroke (r_g_=-0.43, P=0.08), albeit much less significant.

#### Bivariate genetic correlations across substance use disorders

Many genetic correlations were identified across SUDs (Figure S11-S12, Table S10), with the exception for the number of cigarettes smoked per day and smoking cessation. Negative correlations were identified between the age of smoking initiation, ever smoked and drinks per week. The age of smoking initiation is coded so that a young age of smoking initiation related to a low score. Therefore, a negative correlation between smoking initiation and, for example alcohol dependence, is inversely positively correlated with a young age of smoking initiation. The genetic correlations across smoking, cannabis use and alcohol phenotypes, reveal a common genetic liability to substance use disorders.

#### Bivariate genetic intercorrelations across all phenotypes

Other than genetic intracorrelations within psychiatric and substance use disorders, genetic intercorrelations were also observed between the phenotypes (Figure S11-S12, Table S10). Psychiatric disorders were typically genetically correlated with the age of smoking initiation and whether people had ever smoked, and to a lesser extent with cannabis use. ADHD, SCZ and PTSD were additionally genetically associated with the number of cigarettes smoked per day and smoking cessation. Apart from that, many genetic correlations were observed with other brain-related phenotypes. Most profound negative genetic links were observed with intelligence, educational attainment and cognitive performance. Interestingly, cannabis use and alcohol use were positively correlated with educational attainment and cognitive performance. Intelligence was additionally positively correlated with alcohol use, and to a lesser extent with cannabis use (r_g_=0.33, P=2.4x10^-3^), which was non-significant after correcting for multiple testing. Neuroticism was positively correlated with most psychiatric disorders, as well as alcohol dependency and smoking phenotypes. Neurological disorders were poorly genetically correlated with other brain-related phenotypes. ALS was negatively correlated with educational attainment and cognitive performance, and Alzheimer’s disease was negatively correlated with educational attainment. Many stroke subtypes were also negatively correlated with educational attainment and cognitive performance. Sleeping phenotypes were moderately positively correlated with psychiatric disorder and SUDs. These were most profoundly correlated with BIP, SCZ, age of smoking initiation, ever smoked, cigarettes smoked per day and smoking cessation. These findings reveal a large genetic overlap among psychiatric disorders, SUDs and other brain-related phenotypes, but not across neurological disorders.

#### Cell type-specific gene expression in the 10x Genomics dataset

The full 10x Genomics dataset (n=1,306,127) was randomly subsampled to a subset with one twelfth the original size (n=108,844) in an effort to address computational limitations that occurred in the full dataset. Additionally, k-means clustering (k=20) was chosen as the optimal clustering for cell type identification, because projection onto cells in a t-SNE plot could relatively distinguish clusters the most, although a degree of overlap was unavoidable (Figure S2). A further comparison of cell cluster compositions between the full dataset and subset didn’t reveal any subset-specific biases for cell clusters, which indicated that the chosen subset was representative of the full dataset (Figure S3).

Initially 8,306 cells were removed from the subset for failing quality control criteria, which included one complete cell cluster. Three more clusters which comprised 102 cells were removed since most of their cells failed quality control. A total of 6,419 non-expressed genes were also removed from the subset.

Next, cell types were identified in the 10x Genomics subset through an approach of linking specifically expressed genes to known marker genes from literature (Table S3). Through this approach various neuronal, glial and enteric nervous cell types could be recovered (Figure S4); ranging from neuroblasts and intermediate progenitors to matured neurons, interneurons, astrocytes and microglia, oligodendrocytes, Cajal-Retzius cells, (vascular) endothelial cells and finally enteric neuronal and glial cells.

Neuroblasts were recovered in two clusters and were marked by *Igfbpl1* and *Islr2*, the expression of *Eomes* identified intermediate progenitors (Table S4). Neurons (glutamatergic) also consisted of two clusters with *Gria2*, *Neurod2* and *Neurod6* marking their presence. Specific expression of *Dlx2* was clearly recovered in both interneuron clusters, while *Dlx6os1* stood out in only one cluster. Identification of two astrocyte clusters followed from *Aldoc* and *Dbi*. The expression of *Csf1r*, *Cx3cr1* was indicative of the presence of microglia. The specific expression of *Olig1* and *Olig2* strongly indicated the presence of oligodendrocytes and similarly *Reln* was a highly specific marker for Cajal-Retzius cells. Furthermore, *Igfbp7* was another distinct marker, namely of endothelial cells.

#### Suggestive cell type enrichment findings using 10x Genomics data

Suggestive evidence was provided by one method, pointing towards certain interneurons, certain neuroblasts, certain neurons and excitatory glutamatergic neurons being enriched in several brain-related traits, such as cross-disorders, SCZ, BIP, drinks per week, chronotype, overall sleep duration, excessive daytime sleepiness, educational attainment, intelligence, cognitive performance, BMI, and neuroticism. The associated cell types we found were not pan-neuronal, as only a specific subset of neuronal cell types was associated. The implicated cell types in brain-related traits were fairly specific to brain-related traits when comparing those to cells implicated in human height (Figure S14-15).

#### Suggestive cell type enrichment findings using KI data

Suggestive evidence was provided by one method for cortical interneurons to be implicated in intelligence, cross-disorders, educational attainment and SCZ. Suggestive evidence also pointed towards pyramidal cells (CA1) being implicated in intelligence, cognitive performance, drinks per week, chronotype, overall sleep duration and short sleep duration. MSNs were associated with educational attainment, whether people had ever smoked, chronotype and overall sleep duration. Moreover, suggestive evidence was found for neuroblasts to be associated with drinks per week, embryonic GABAergic neurons with excessive daytime sleepiness and whether people had ever smoked. Endothelial and mural cells were found to be associated with intelligence, cognitive performance and educational attainment, and astrocytes and ependymal cells were implicated in MDD and ADHD. No evidence was found for consistently implicated cell types in ASD and AN. Similarly to the analysis using the 10x Genomics dataset, cell types that were enriched in brain-related phenotypes were generally dissimilar from those enriched in the height phenotype (Figure 3, Figure S19).

#### MAGMA linear regression leniency

We found that 10x Genomics cell type enrichment using MAGMA in linear mode was more lenient than LDSC, DEPICT and MAGMA in top 10% mode and thus prone to type-I error inflation (Figure S16). A positive linear correlation was found between the strength of association of the top associated cell type per phenotype, out of LDSC, DEPICT, MAGMA in top 10% and MAGMA in linear mode, and sample size (r=0.34, P=0.026) (Figure S17A). However, this correlation lost significance (r=0.30, P=0.055) when MAGMA in linear mode was excluded (Figure S17B), suggesting that MAGMA in linear mode is less robust to sample size inflation.

Similarly, KI level 1 and KI level 2 cell type enrichment using MAGMA in linear mode was considerably more lenient than the other methods tested (Figure S23-24). A positive correlation (r=0.32, P=0.039) was found between the strength of association of the top associated cell type for each phenotype, out of LDSC, DEPICT and MAGMA in top 10% mode, and sample size (Figure S25A). This correlation also lost significance (r=0.17, P=0.30) when MAGMA in linear mode was excluded (Figure S25B), strengthening the finding that MAMGA in linear mode is to a larger extent than LDSC, DEPICT and MAGMA in top 10% mode vulnerable to sample size inflation.

### **Supplementary figures**

**Figure S1. Effect of quality control on a 108,844 cell subset of the 10x Genomics mouse brain scRNA-seq dataset.**

Cell quality metrics UMIs per cell (log10-transformed), genes per cell and % mtDNA are shown before- (blue) and after (red) quality control. Red dotted lines in the blue plots indicate the filter ranges within which cells were kept. These ranges were set to; μ±3σ UMIs (log10), 1000 up to μ+3σ genes and below μ+3σ % mtDNA. Quality control reduced the subset to 100,538 cells in total. UMI: Unique Molecular Identifier, mtDNA: mitochondrial DNA.

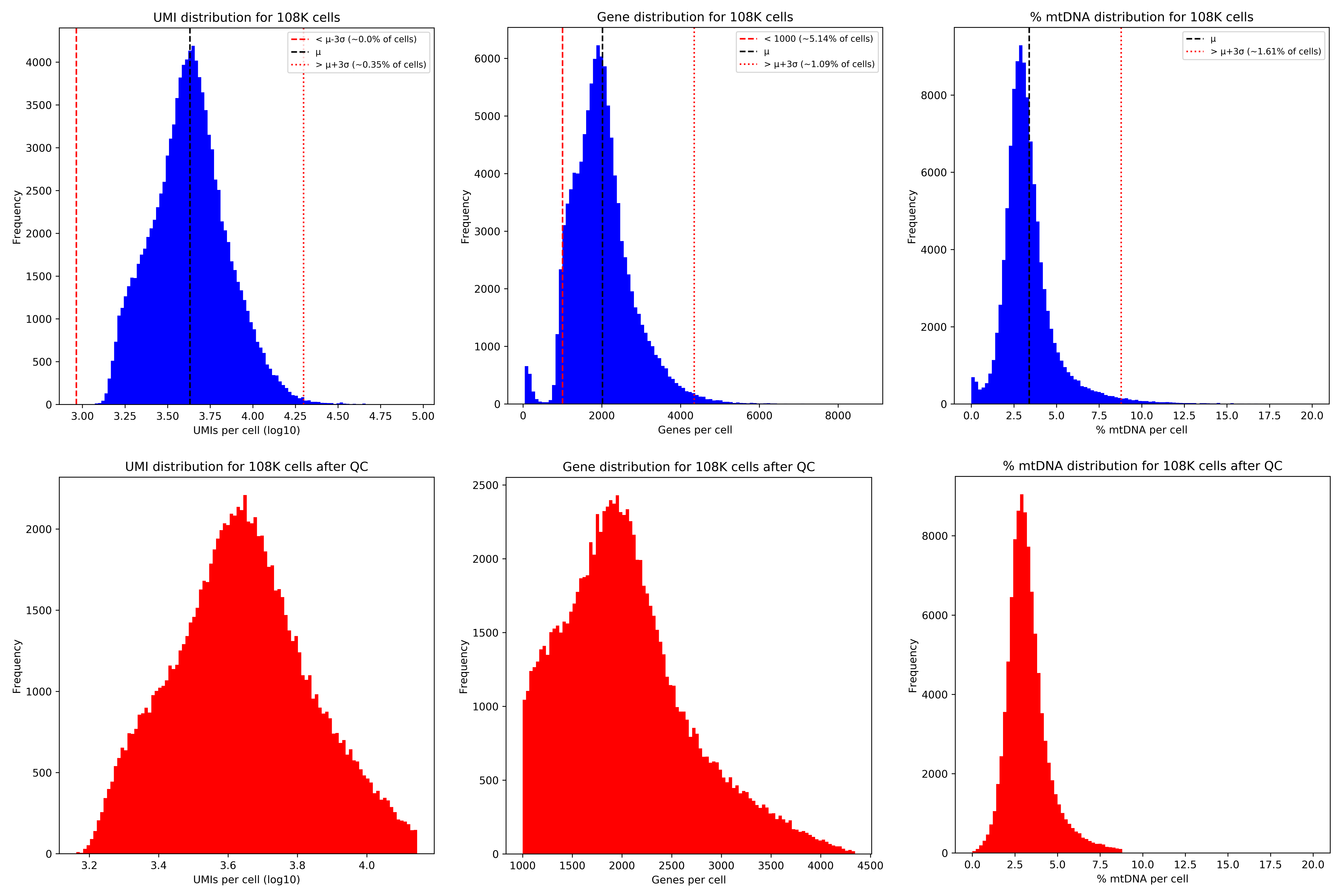

**Figure S2. K-means (k=20) clustering results for 1,306,127 mouse brain cells from the 10x Genomics scRNA-seq dataset.**

These clustering results were comparatively the most distinguishable and least overlapping compared to other k-means clustering results (ranging from k=2 to k=19, data not shown) and were therefore chosen. The separation of clusters is somewhat limited for e.g. clusters 1, 2, 3, 4, or 5, 7. Cells belonging to clusters 18, 19 and 20 do not stand out clearly due to their low numbers.

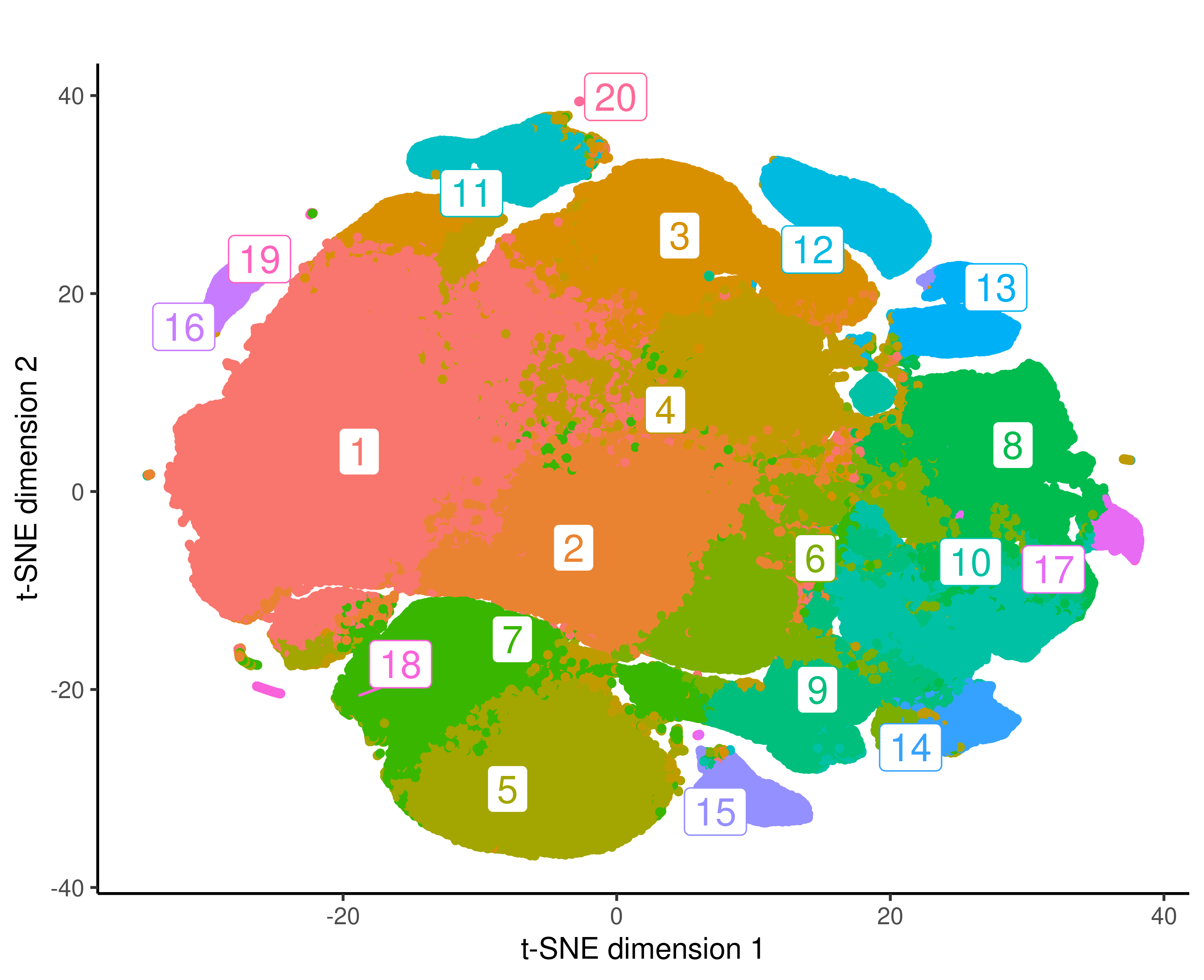

**Figure S3. Comparison of cluster compositions between the full 10x Genomics mouse brain scRNA-seq dataset and subsets.**

K-means clustering (k=20) was used to obtain the clusters, subsets were randomly subsampled from the full 1.3 million cell dataset (‘1.3 M cells’). Cluster compositions all strongly resembled each other, suggesting that no subset-specific biases towards any clusters were present.

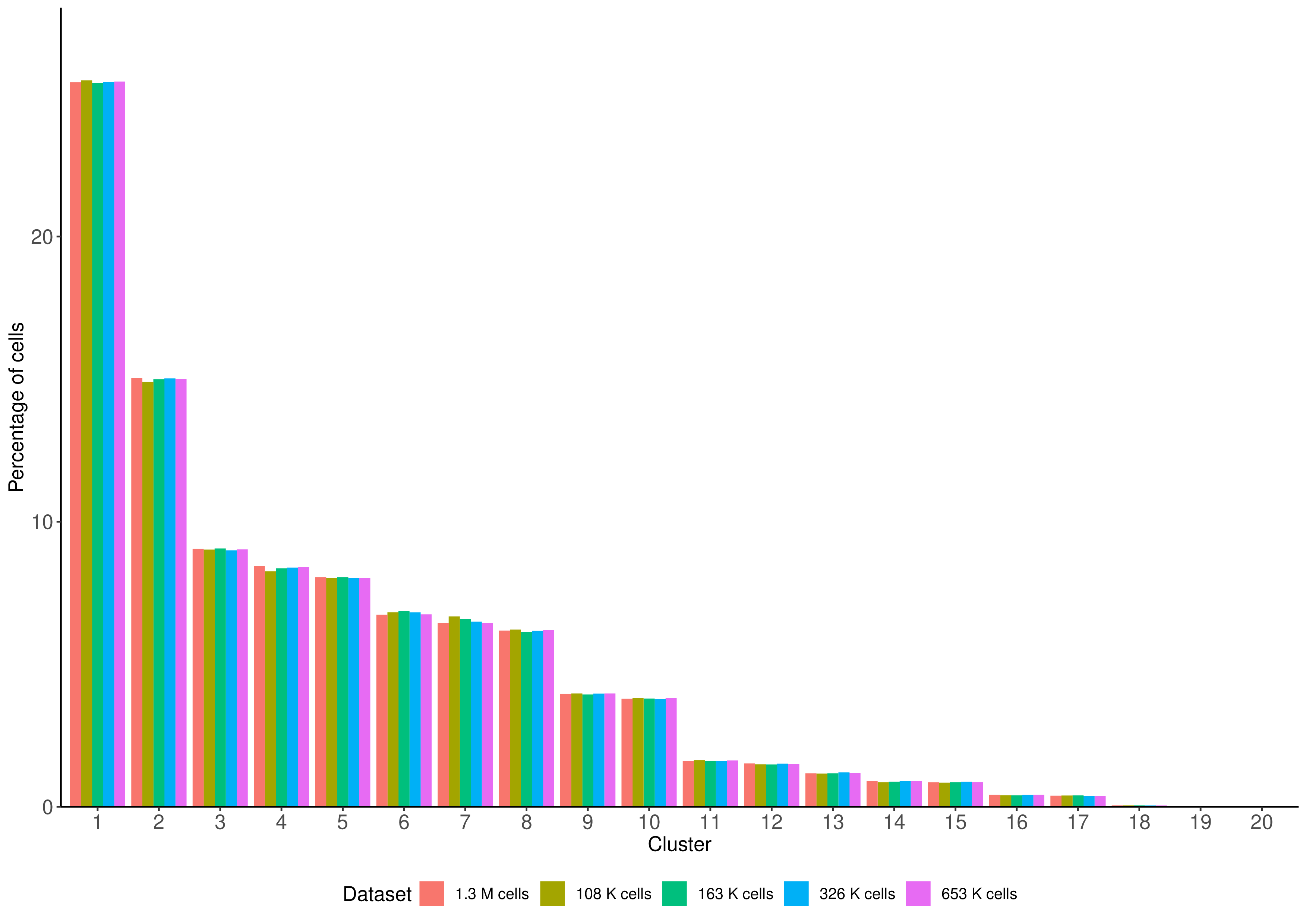

**Figure S4. 10x Genomics dataset with annotated cell types for 100,436 cells.**

Cell types were identified by matching the most-specifically expressed genes for k-means clustering (k=20) cell clusters to known brain cell type marker genes from literature. Four clusters are not shown since they did not pass quality control. The majority of cells are neuronal, yet a variety of glial, enteric and endothelial cells were also identified. Neuronal cell types and astrocytes are divided into multiple clusters. Few clusters are completely separated and most cluster share varying degrees of overlap with other clusters. In the bottom left a heterogeneous region of neurons, interneurons and neuroblasts can be seen.

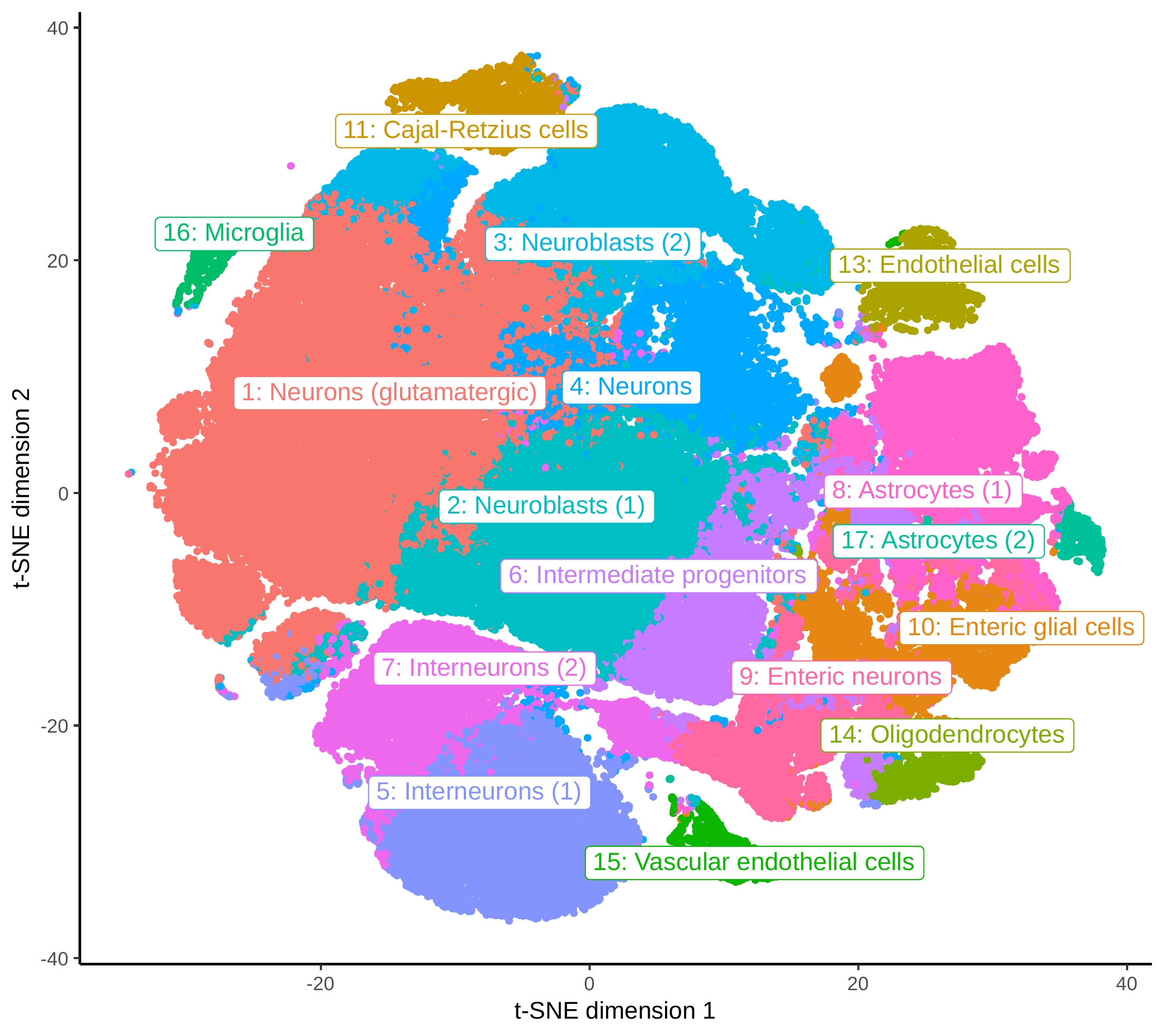

**Figure S5. LDSC sensitivity analysis in SCZ using the KI dataset.**

The LDSC parameters used by Skene et al.^4^ were considered the default parameters. Sensitivity analysis in SCZ shows that removing the Hapmap3 SNPs restriction leads to inflated results. Restricting only to significant SNPs (P<0.05) results in loss of statistical power. The red line indicates the Bonferroni threshold (P< 0.05/(24*42)). For further analyses, we used the most recent version of the GENCODE reference genome (v33) and the most recent LDSC version.

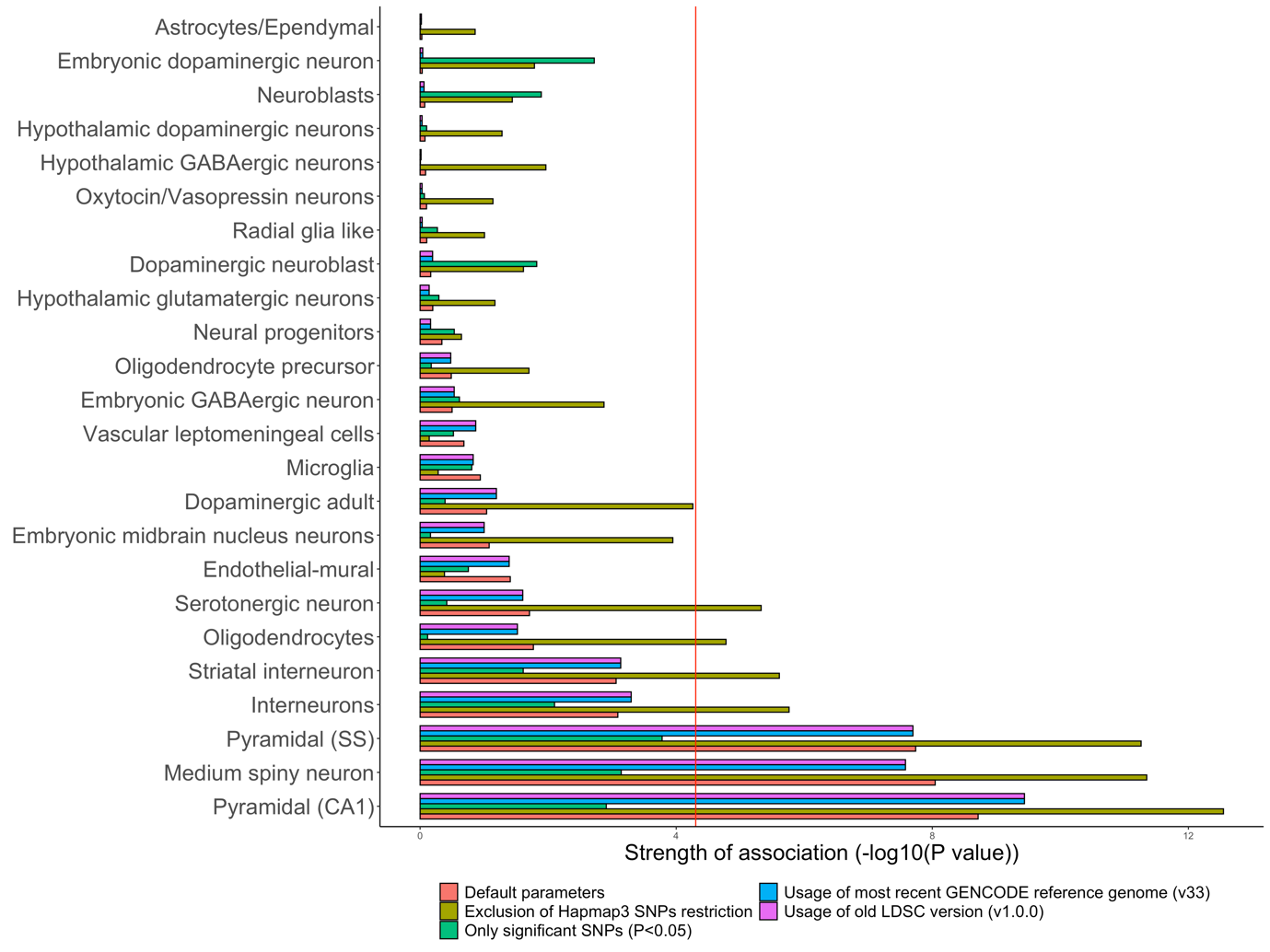

**Figure S6. MAGMA sensitivity analysis in SCZ using the KI and 10x Genomics dataset.**

*Sensitivity analysis in SCZ shows that restricting to Hapmap3 SNPs improves power to detect associated cell types, whilst not inflating enrichment scores of cell types that are not associated. Additionally, exclusion of the MHC region has little effect on cell type enrichment results. The red line indicates the Bonferroni threshold (KI; P< 0.05/(24*42), 10x Genomics; P< 0.05/(16*42)). Sensitivity analyses were performed in SCZ using the KI dataset (top row) with MAGMA (linear mode) and MAGMA (top 10% mode) and using the 10x Genomics dataset (bottom row).*

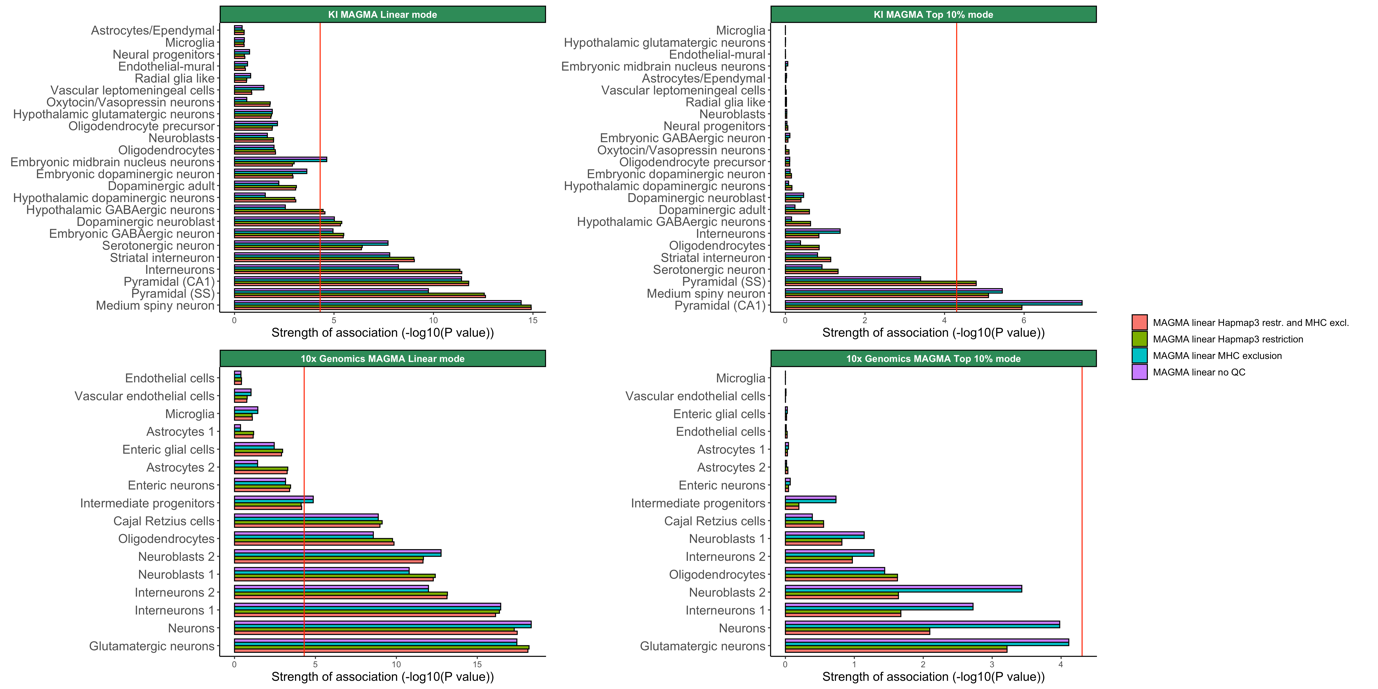

**Figure S7. Differences in cell type enrichment results using MAGMA version 1.07 and 1.08.**

All analyses were performed using 42 summary statistics that were restricted to HapMap3 SNPs and had the MHC region excluded. The enrichment results are parsed by scRNA dataset, i.e. KI and 10x Genomics, and MAGMA enrichment mode, i.e. linear and top 10% mode. **A.** Association of SCZ with cell types derived from the KI and 10x Genomics dataset. The red line indicates the Bonferroni threshold (KI; *P< 0.05/(24*42), 10x Genomics; P< 0.05/(16*42)).* ***B.*** *Boxplots of cell type association values (-log10(P value)).* ***C.*** *Scatterplots of association values (-log10(P value)) estimated using version 1.07 vs association values estimated using version 1.08.* ***D.*** *Histograms of association values (-log10(P value)).*

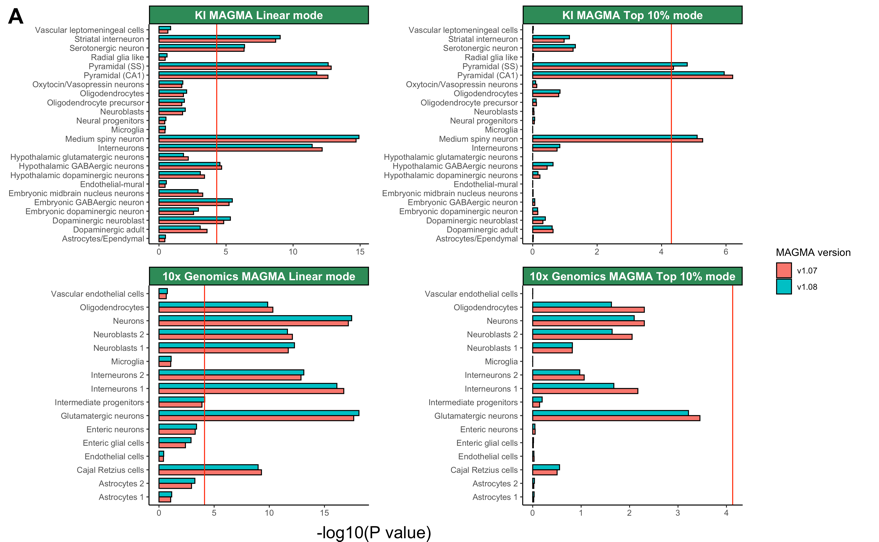

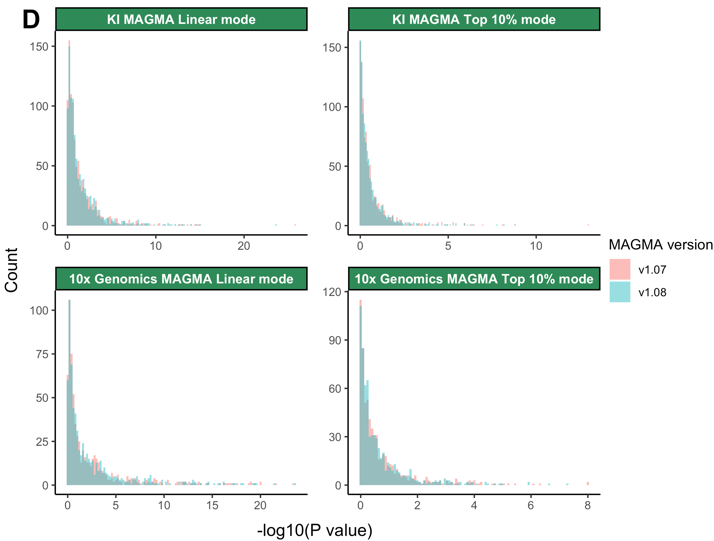

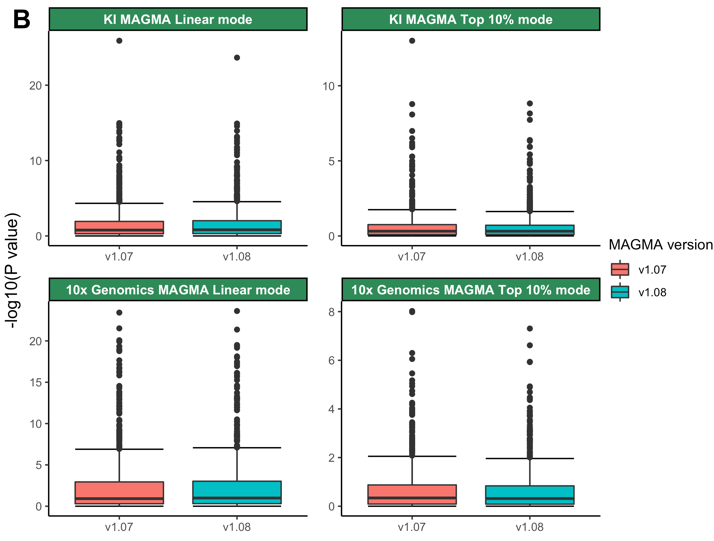

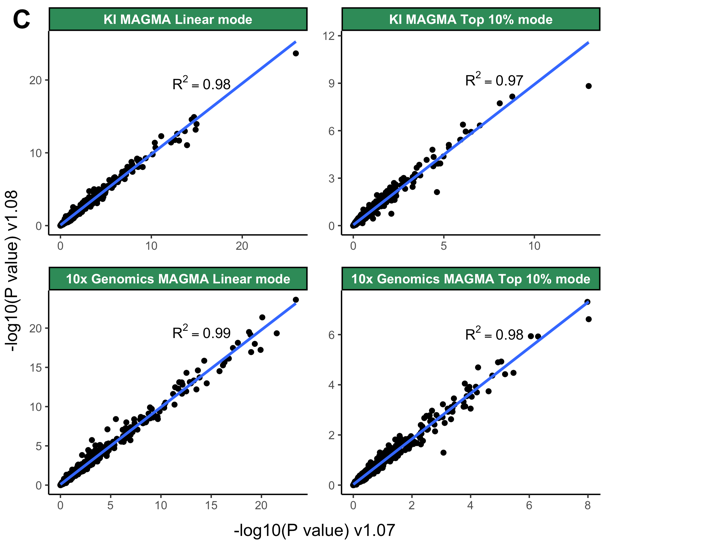

**Figure S8. DEPICT sensitivity analysis in SCZ using the KI dataset.**

*It is shown that restricting to Hapmap3 SNPs reduces power to identify associated cell types. Applying no quality control leads to improved statistical power, whilst not inflating enrichment results for cell types that are not associated. The black line indicates the Bonferroni threshold (P< 0.05/(24*42)).*

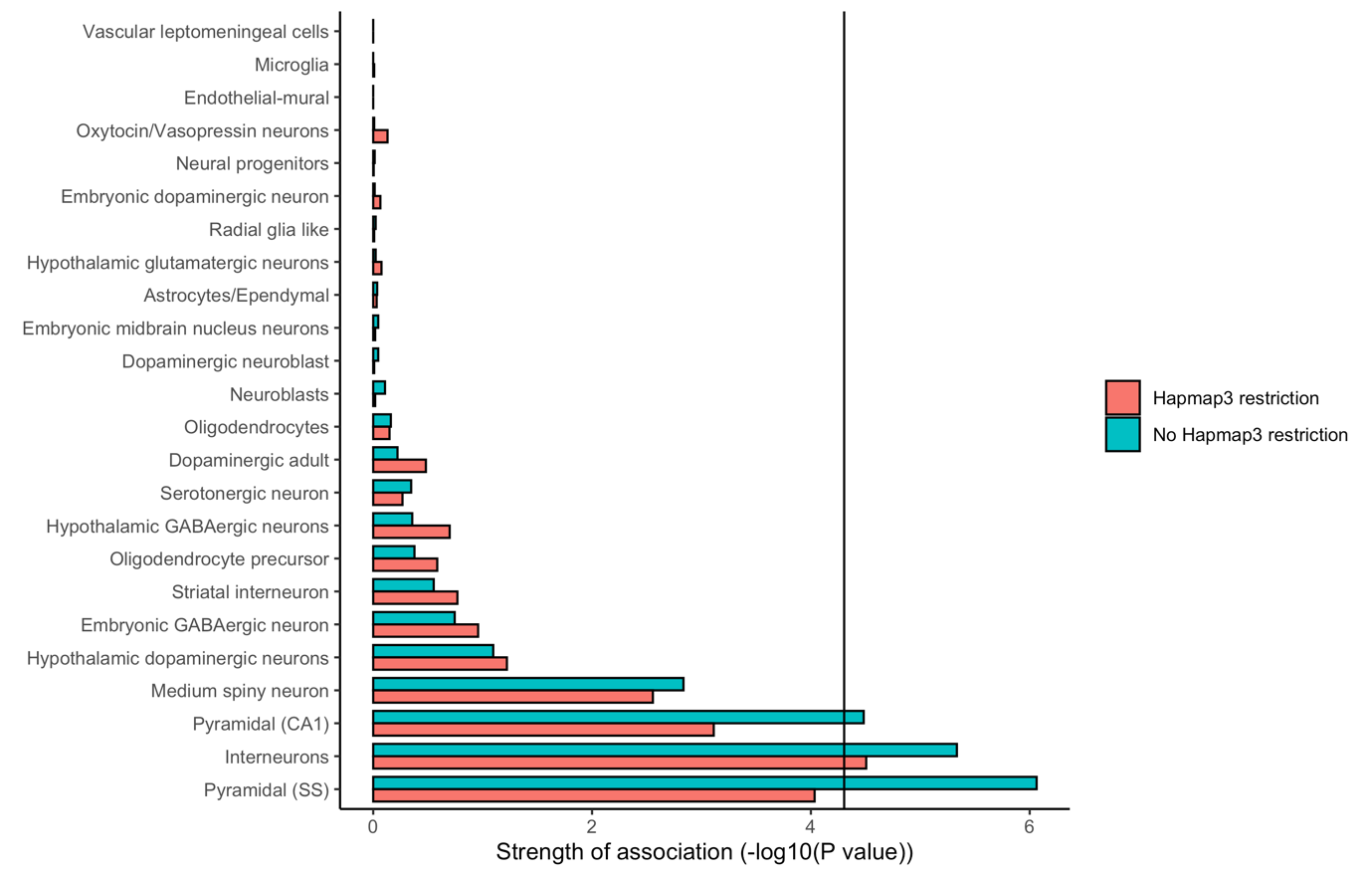

**Figure S9. SNP-h^2^ estimates of 41 brain-related phenotypes and 1 non-brain-related phenotype.**

*Abbreviations: ADHD; attention deficit hyperactivity disorder, ALS; Amyotrophic lateral sclerosis, BMI; body mass index, SUD: substance-use disorder.*

SNP-h^2^ *was estimated using LDSC using the 1000 Genomes Project phase 3 baseline model and regression weights for Hapmap3 SNPs. Error bars represent one standard error.*

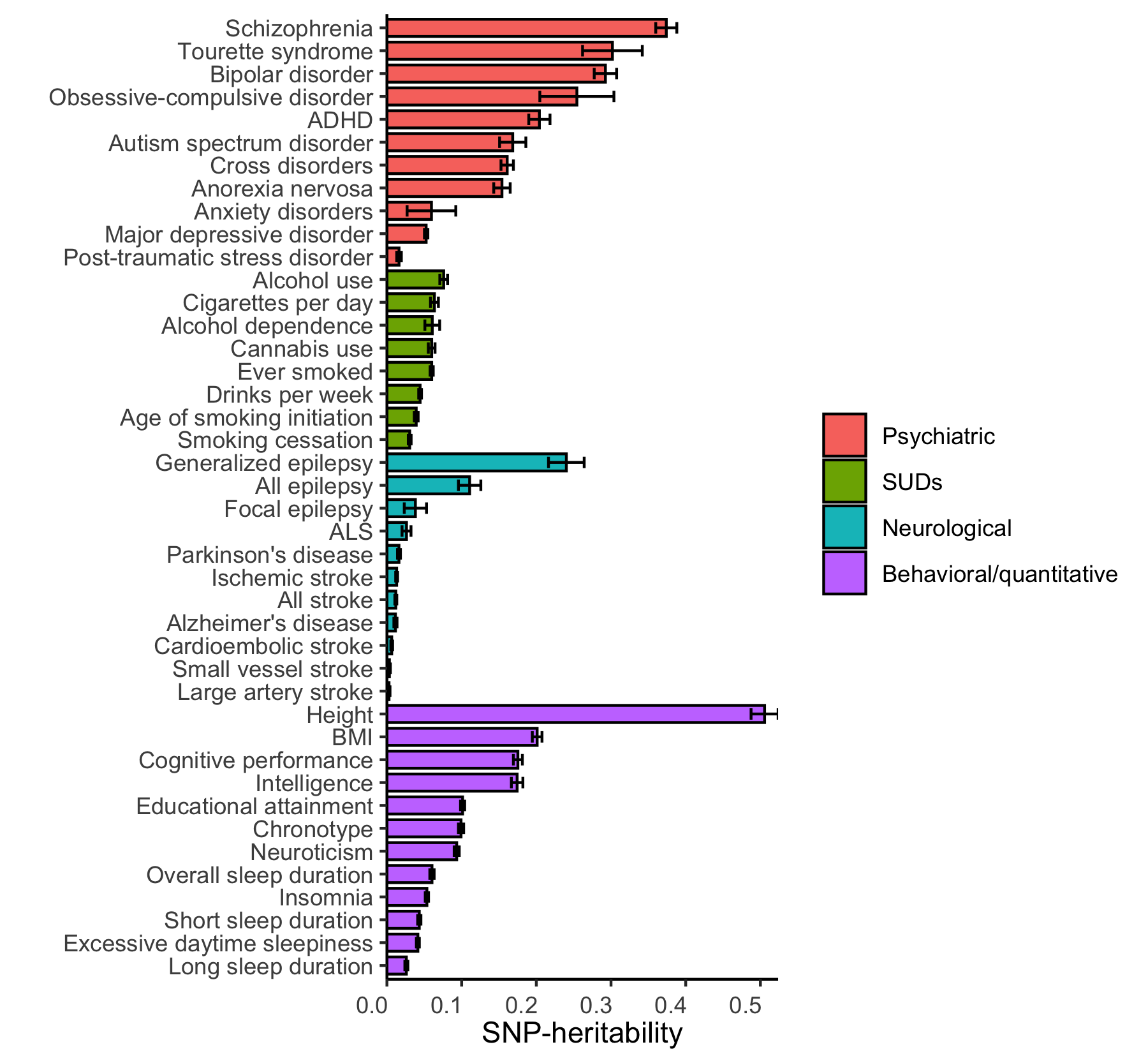

**Figure S10. Correlation between the sample size and SNP-h^2^ estimates.**

*Abbreviations: SUD; substance-use disorder.*

*No evidence was found for an inflation of SNP-h^2^ estimates in phenotypes with a large sample size (r=-0.23, P=0.14). The grey line represents a linear regression and the area coloured in a lighter shade of grey represents the standard error of the linear regression.*

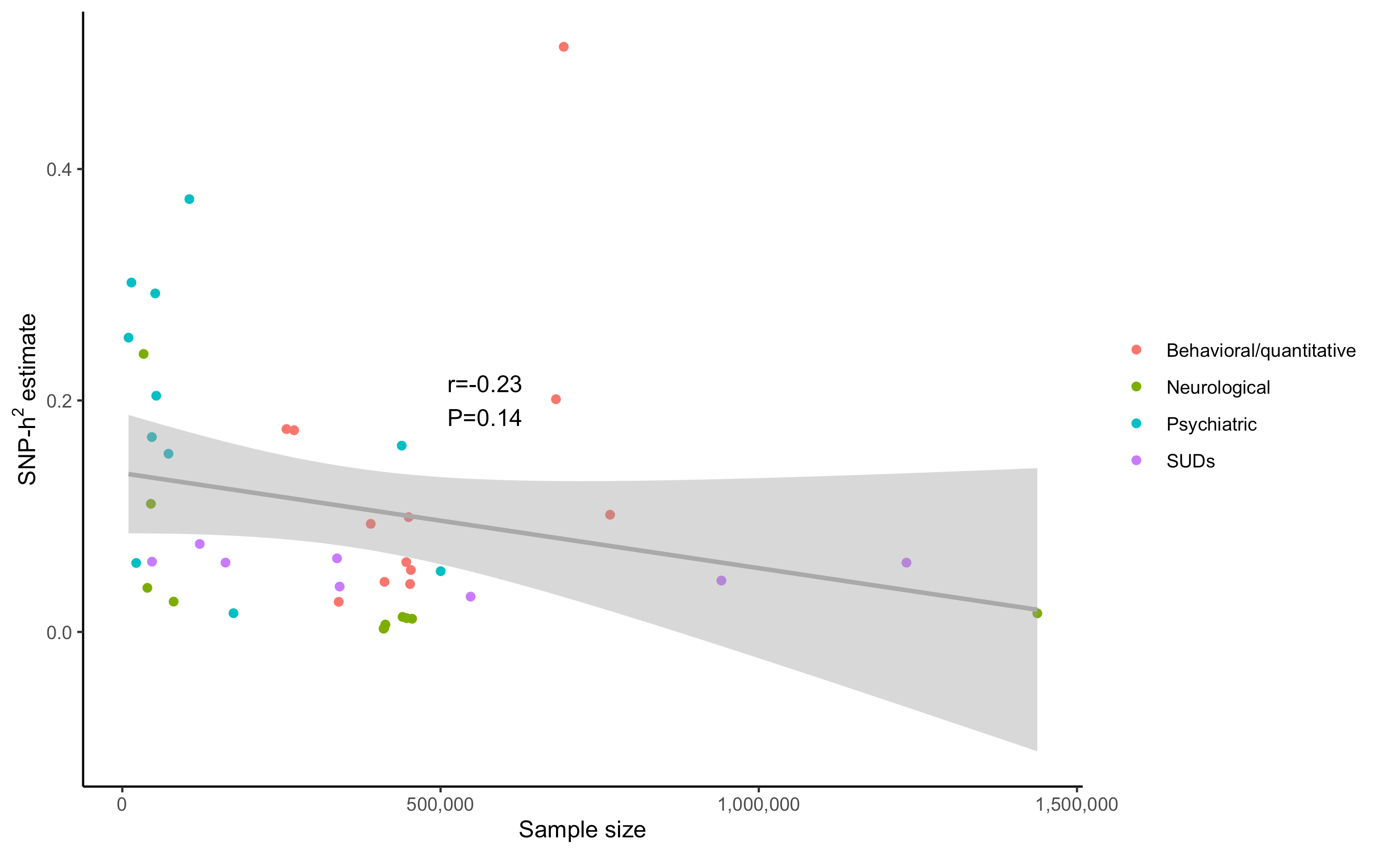

**Figure S11. Bivariate genetic correlations across 41 brain-related phenotypes and 1 non-brain-related phenotype.**

*Abbreviations: ADHD; attention deficit hyperactivity disorder, ALS; amyotrophic lateral sclerosis, BMI; body mass index.*

*Bivariate genetic correlations were estimated using LDSC using the 1000 Genomes Project phase 3 baseline model and regression weights for Hapmap3 SNPs. The lower triangle represents genetic correlation in circles. The size and colour intensity of the circles are proportional to the correlation estimate. Asterisks represent significance after Bonferroni correction (0.05/(42*(42-1)/2)). Shades of blue indicate a positive correlation and shades of red indicate a negative correlation.*

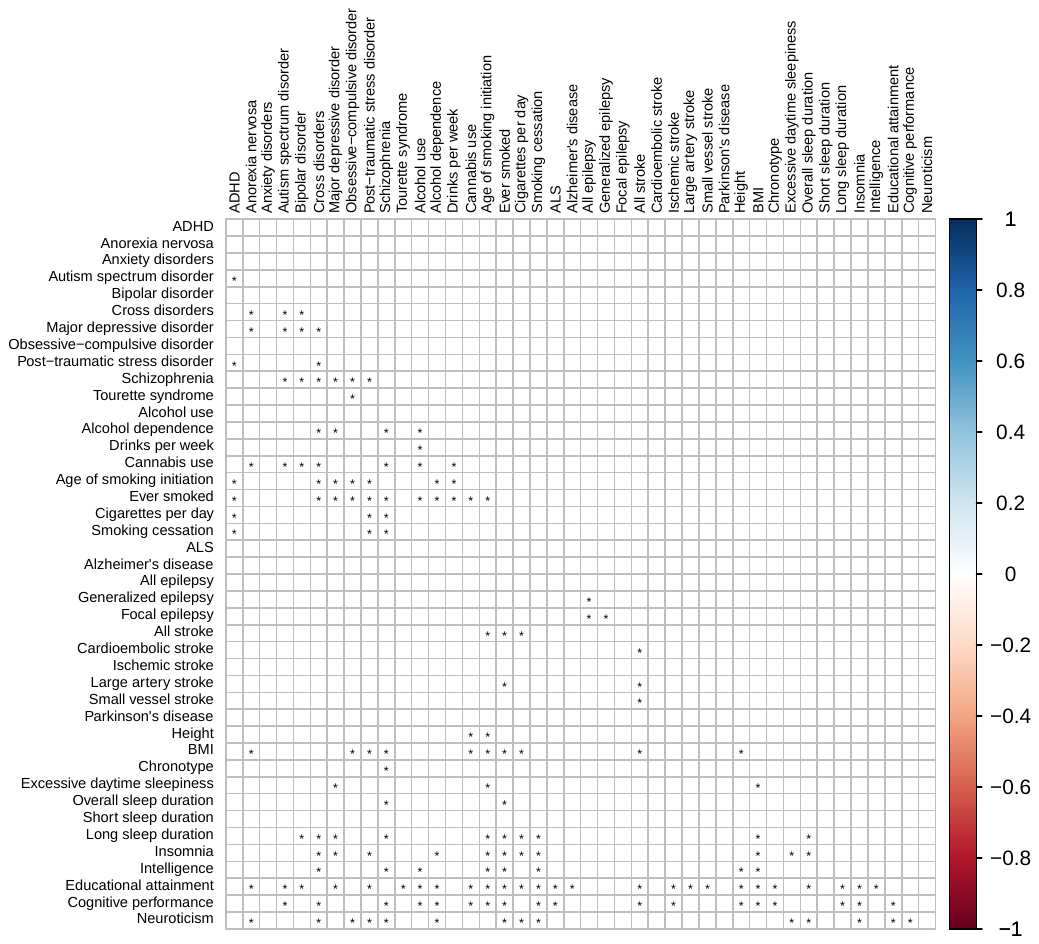

**Figure S12. Forest plots displaying the genetic correlation of each phenotype with all other phenotypes.**

*Abbreviations: ADHD; attention deficit hyperactivity disorder, ALS; amyotrophic lateral sclerosis, BMI; body mass index.*

*Bivariate genetic correlations were estimated using LDSC using the 1000 Genomes Project phase 3 baseline model and regression weights for Hapmap3 SNPs. Genetic correlations that pass Bonferroni correction (0.05/(42*(42-1)/2) are coloured in red. A grey x is shown when no genetic correlation could be estimated. The error bars represent the standard errors.*

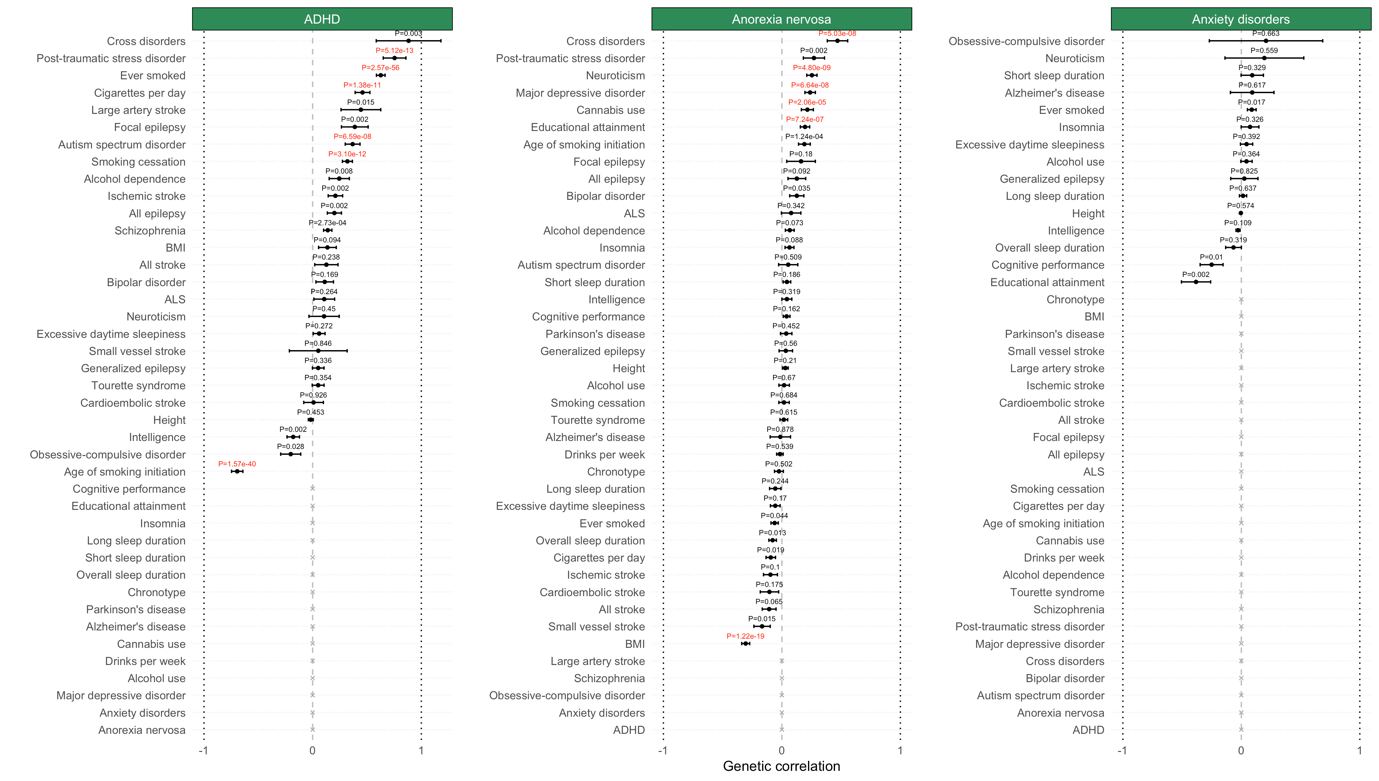

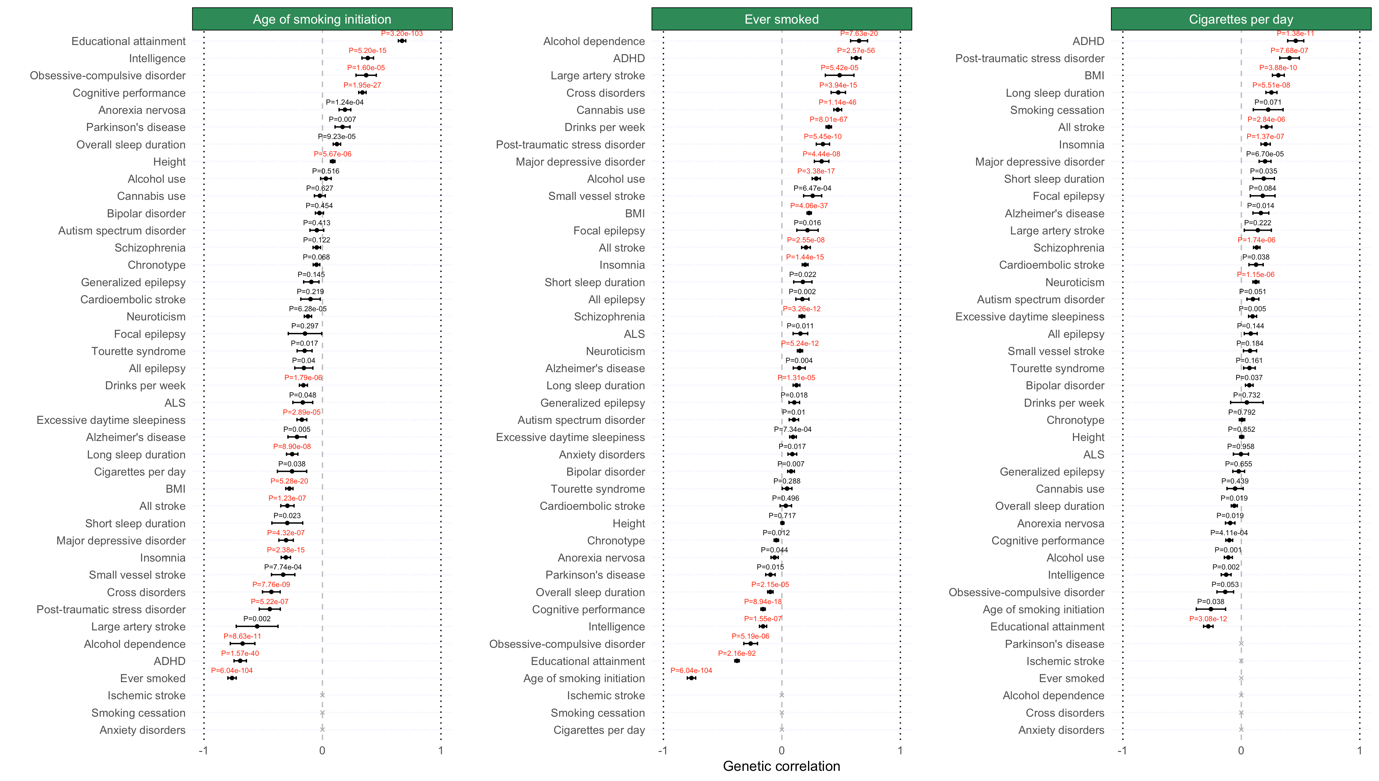

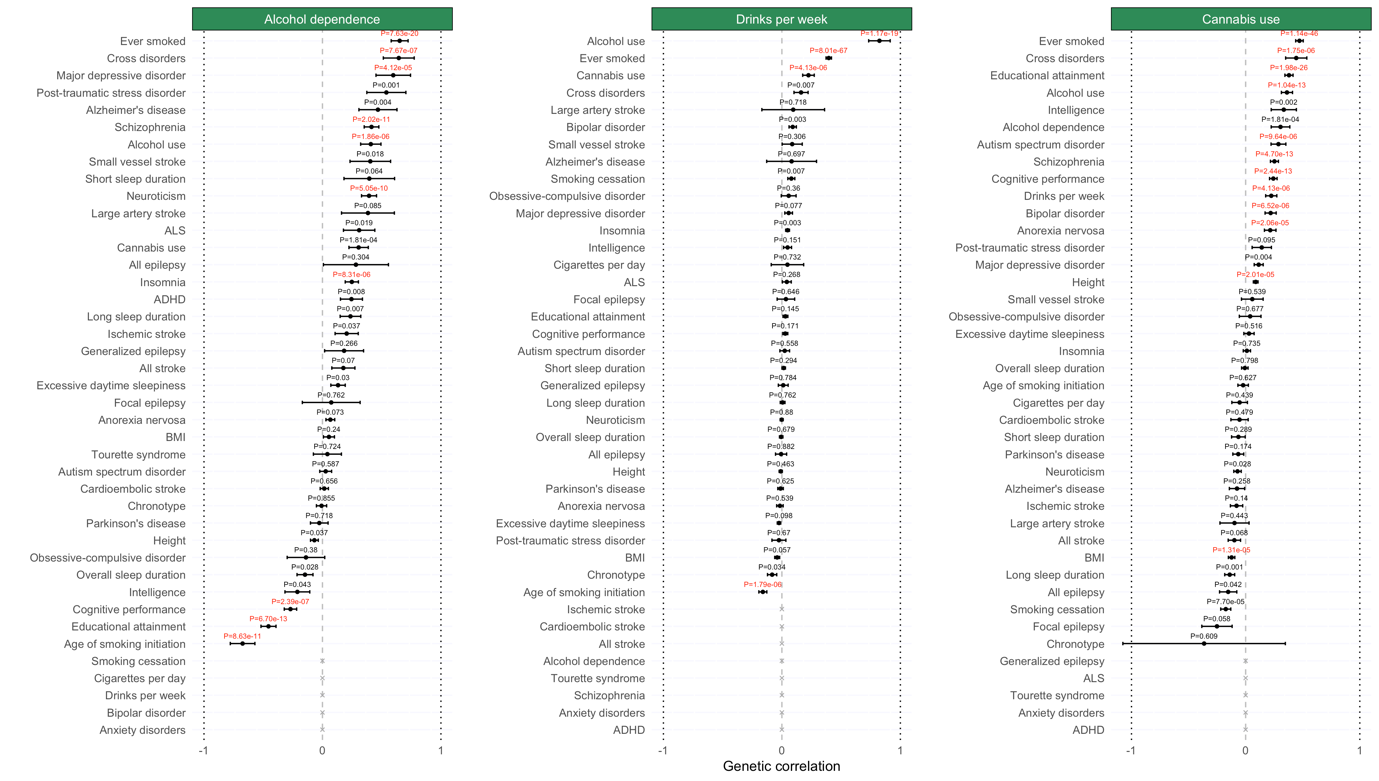

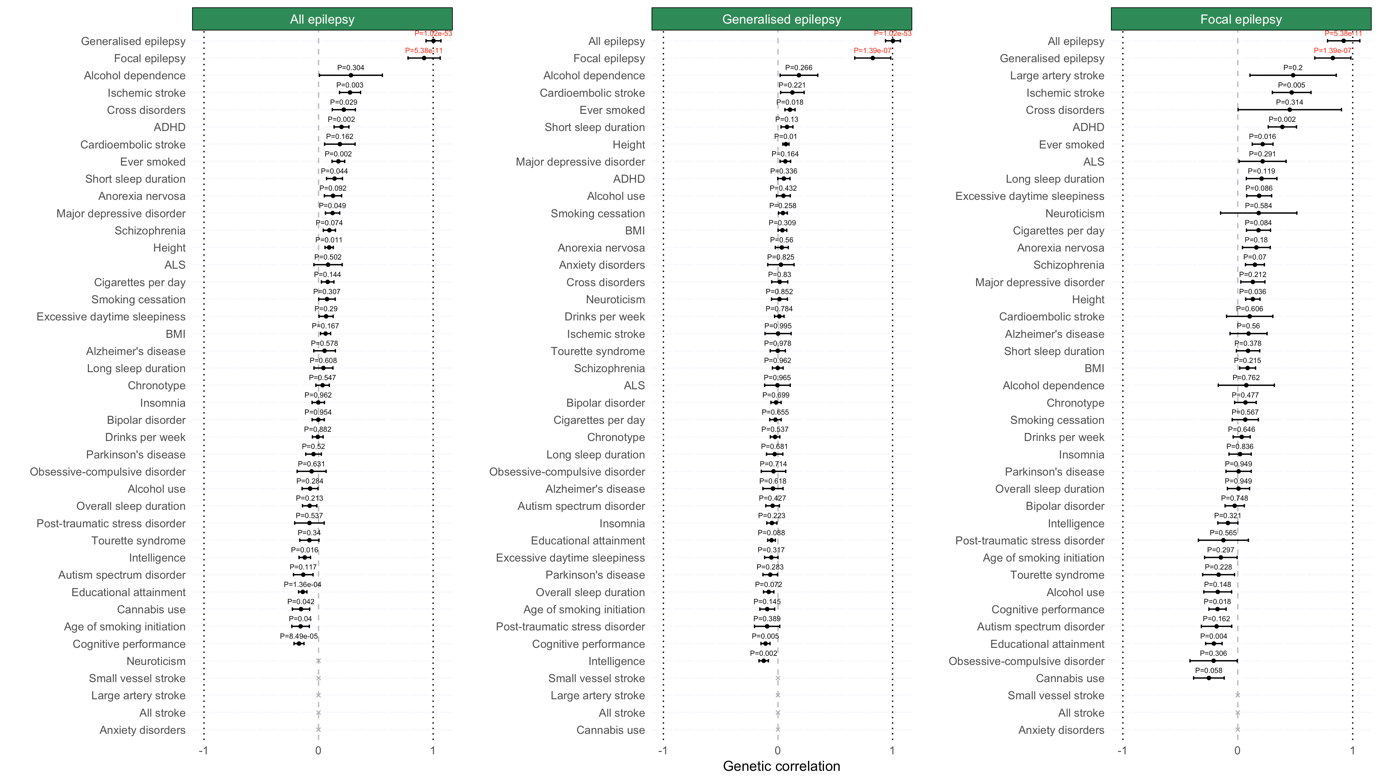

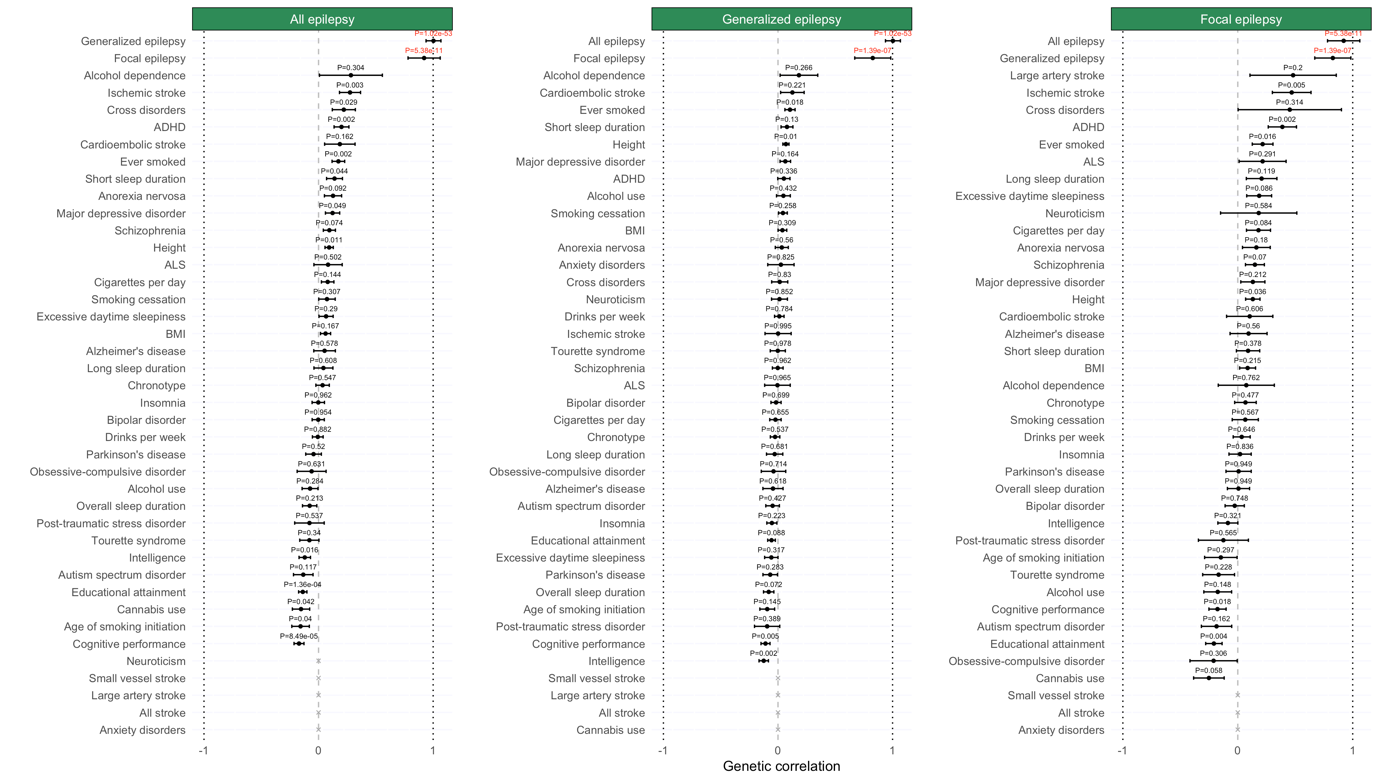

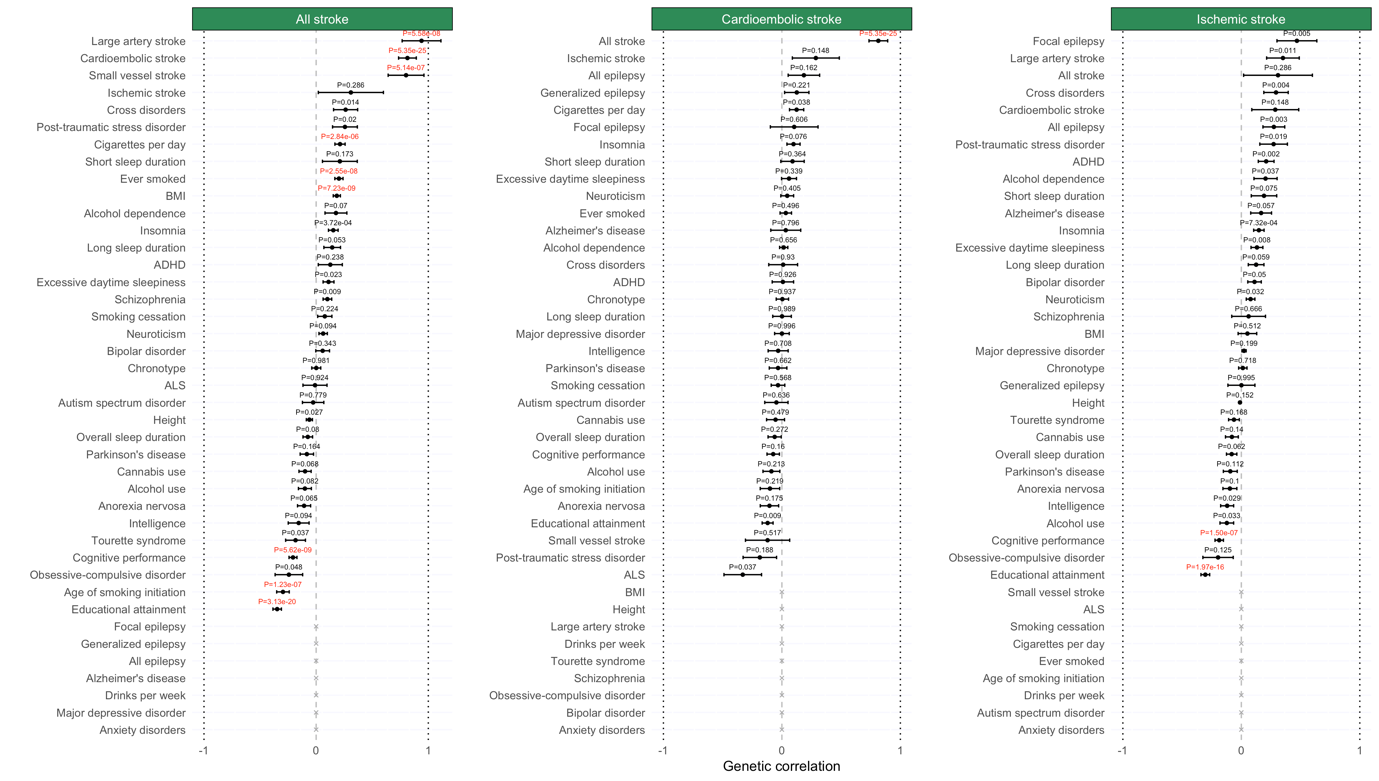

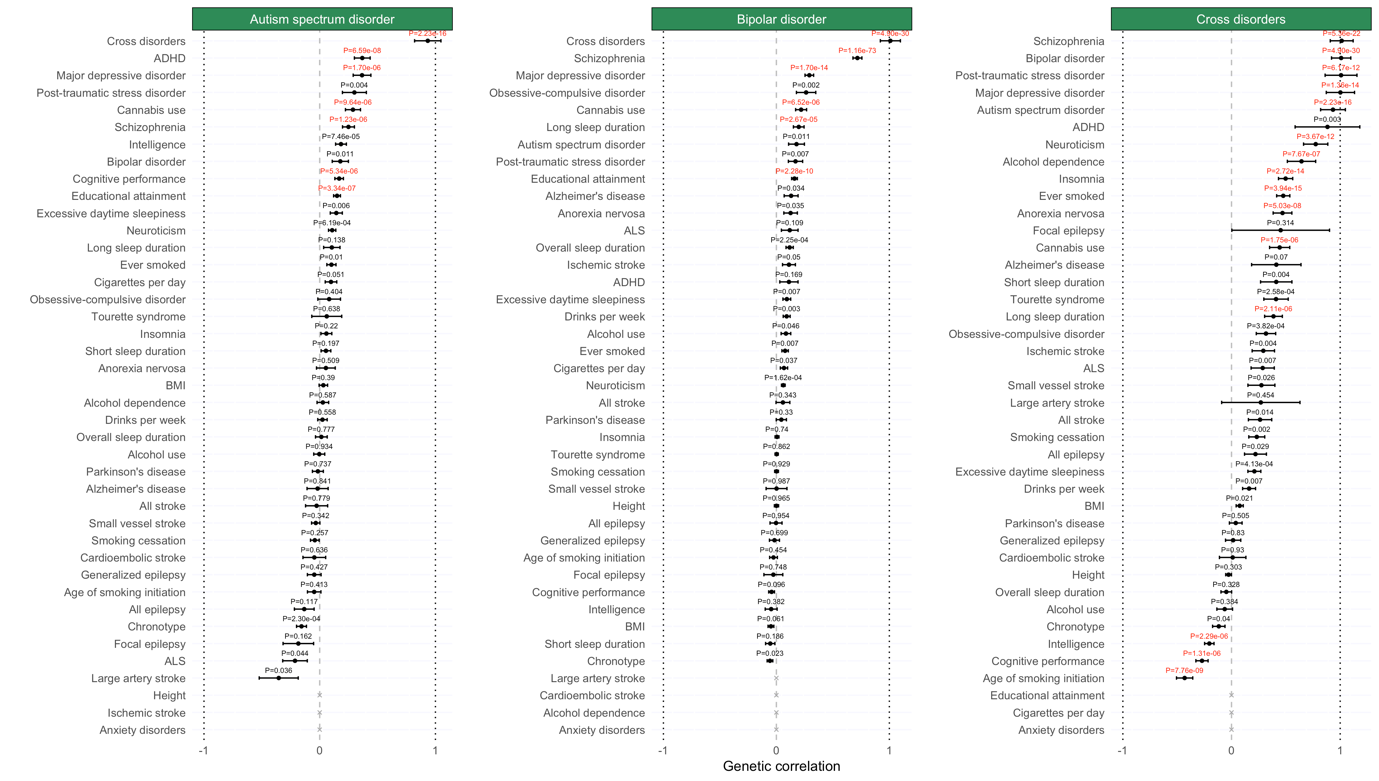

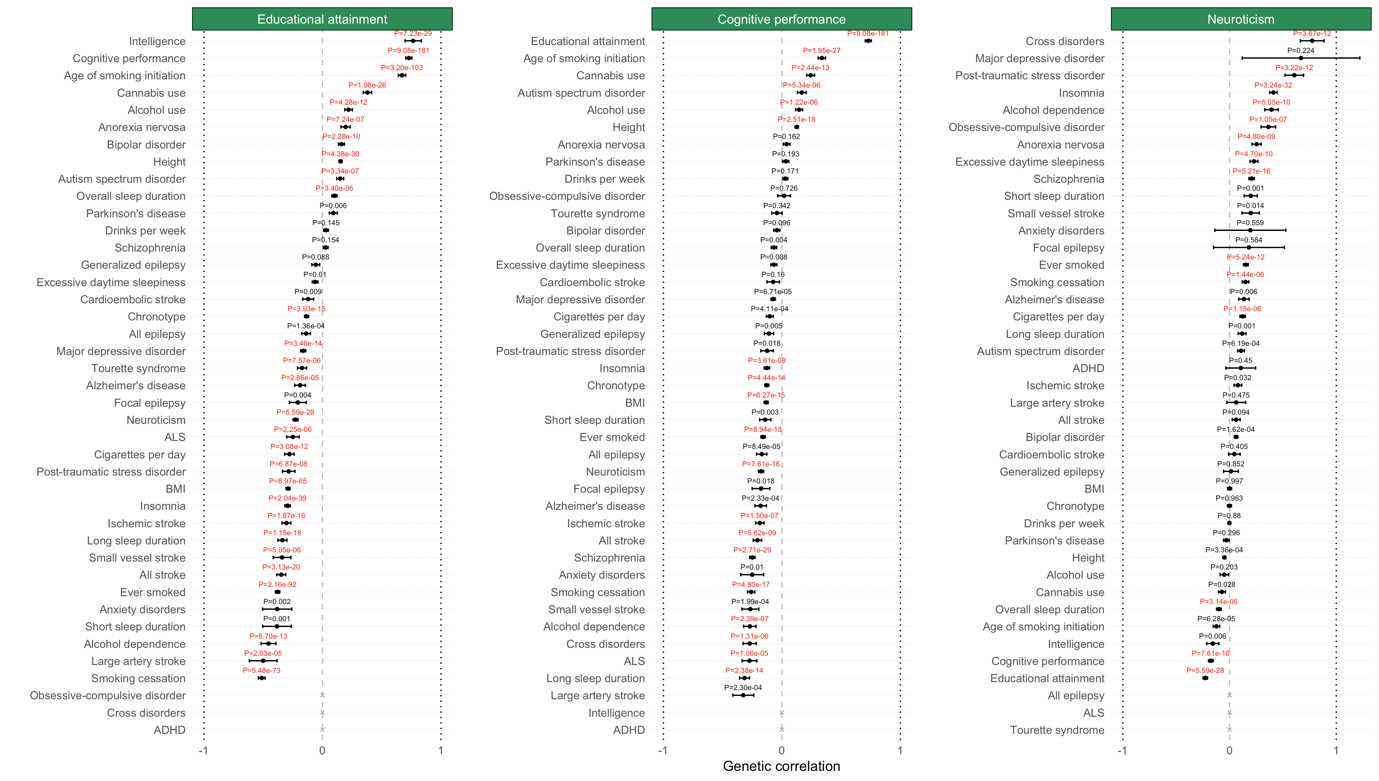

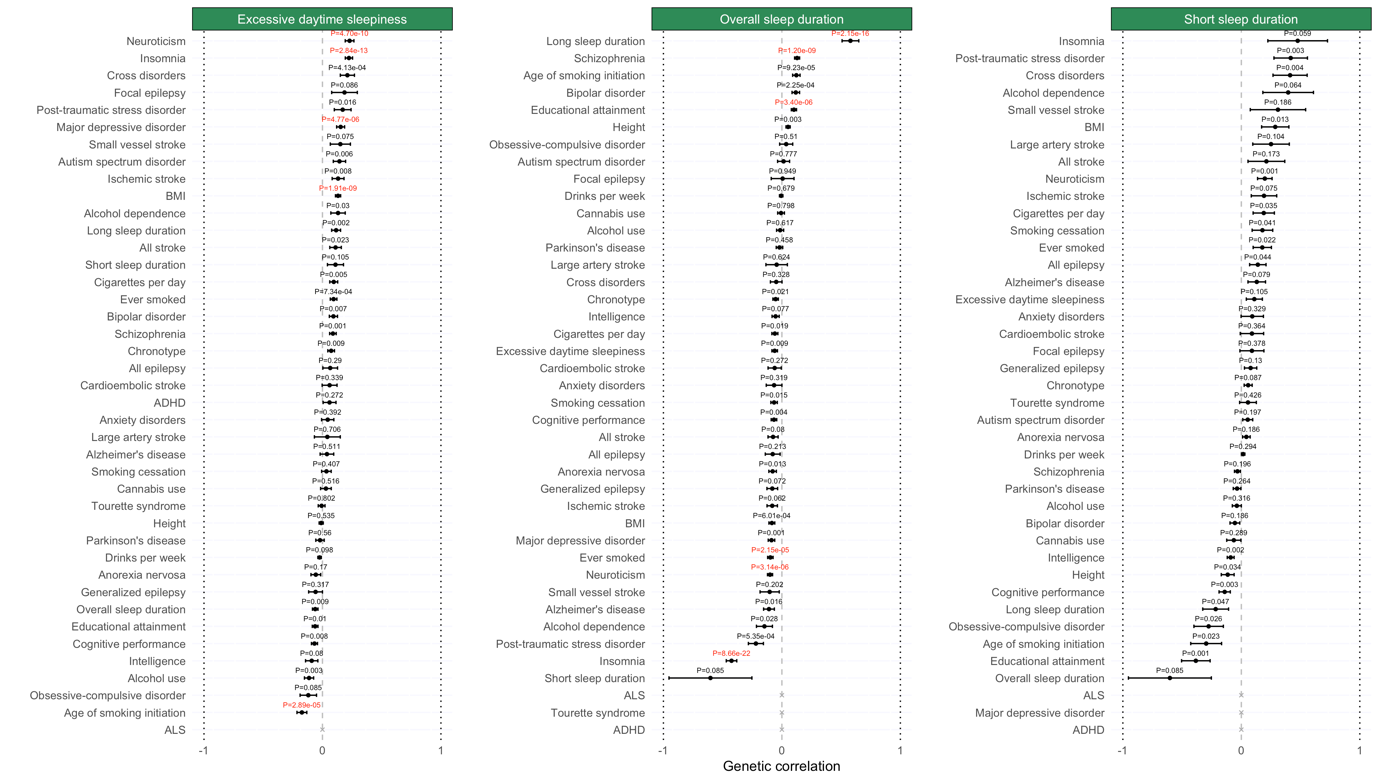

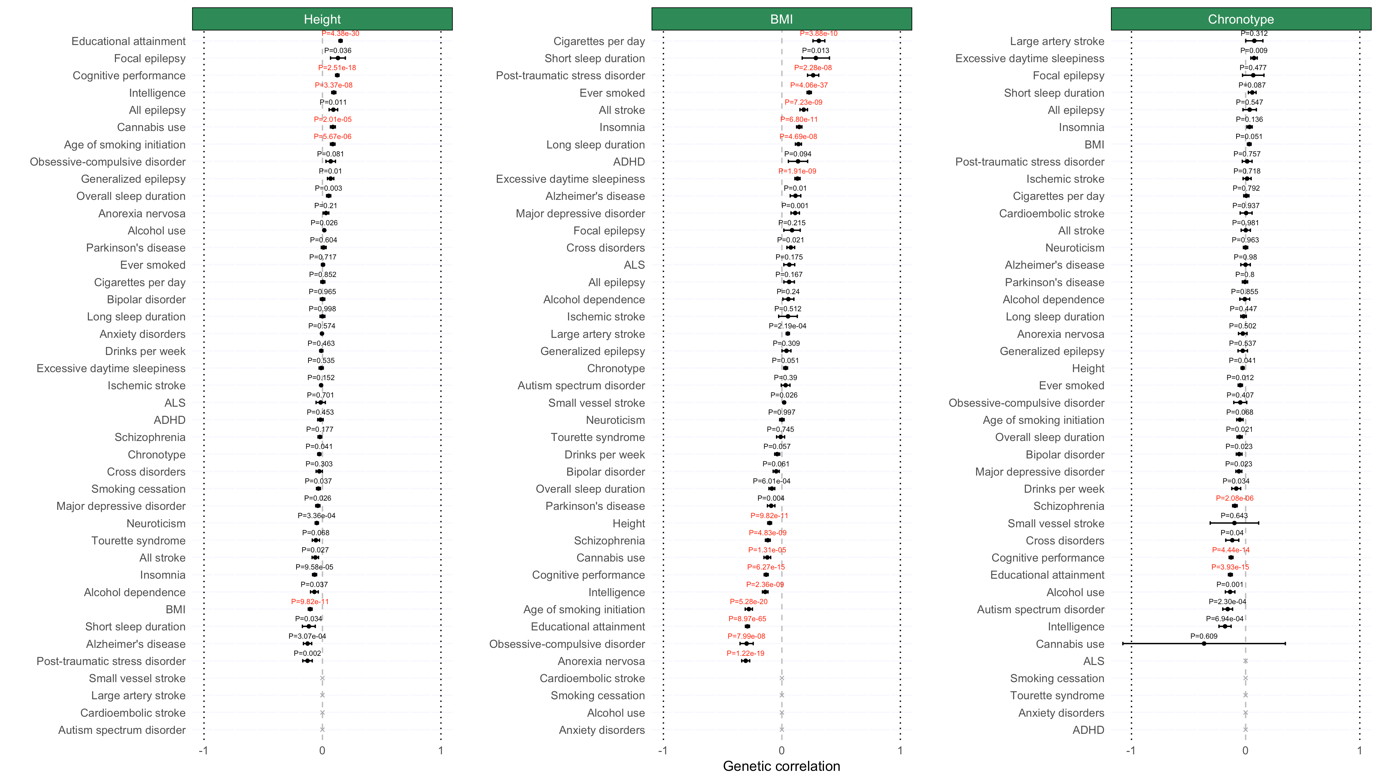

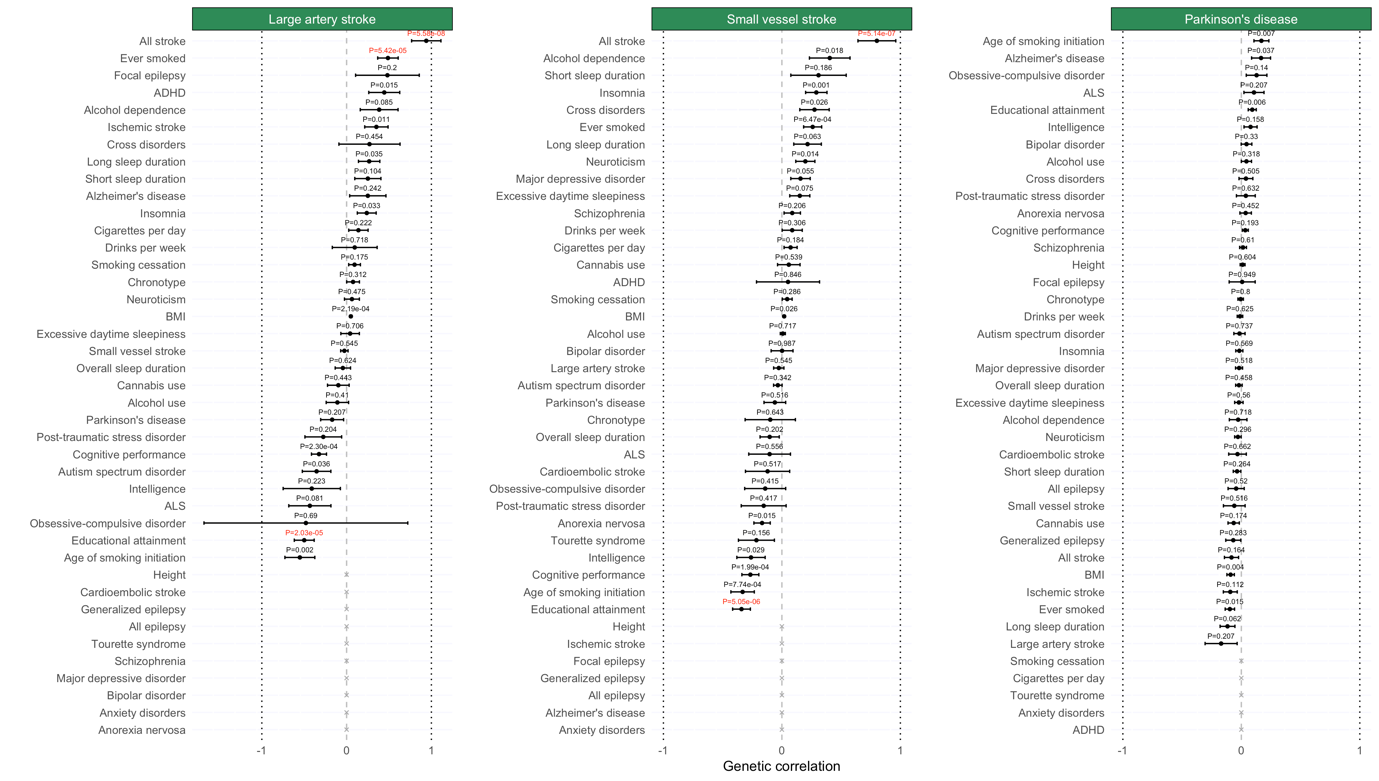

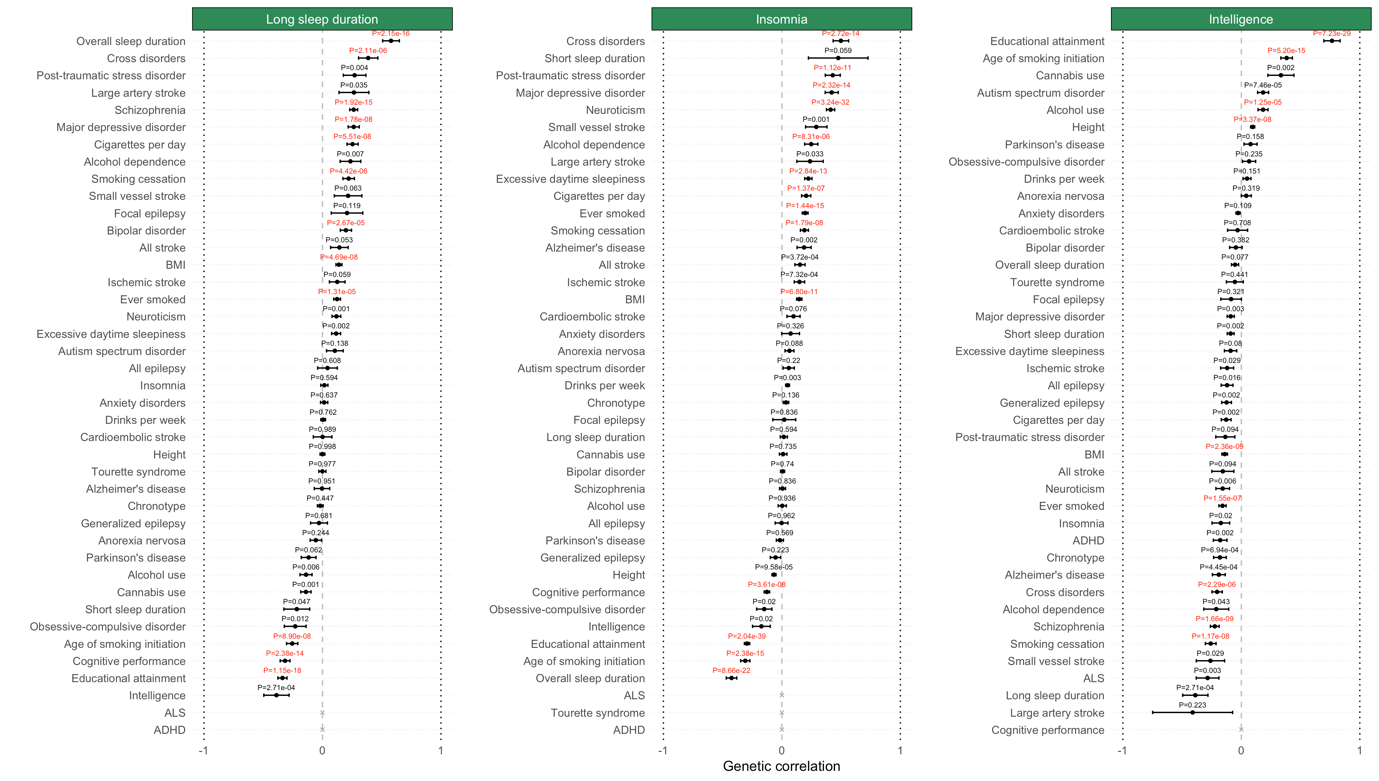

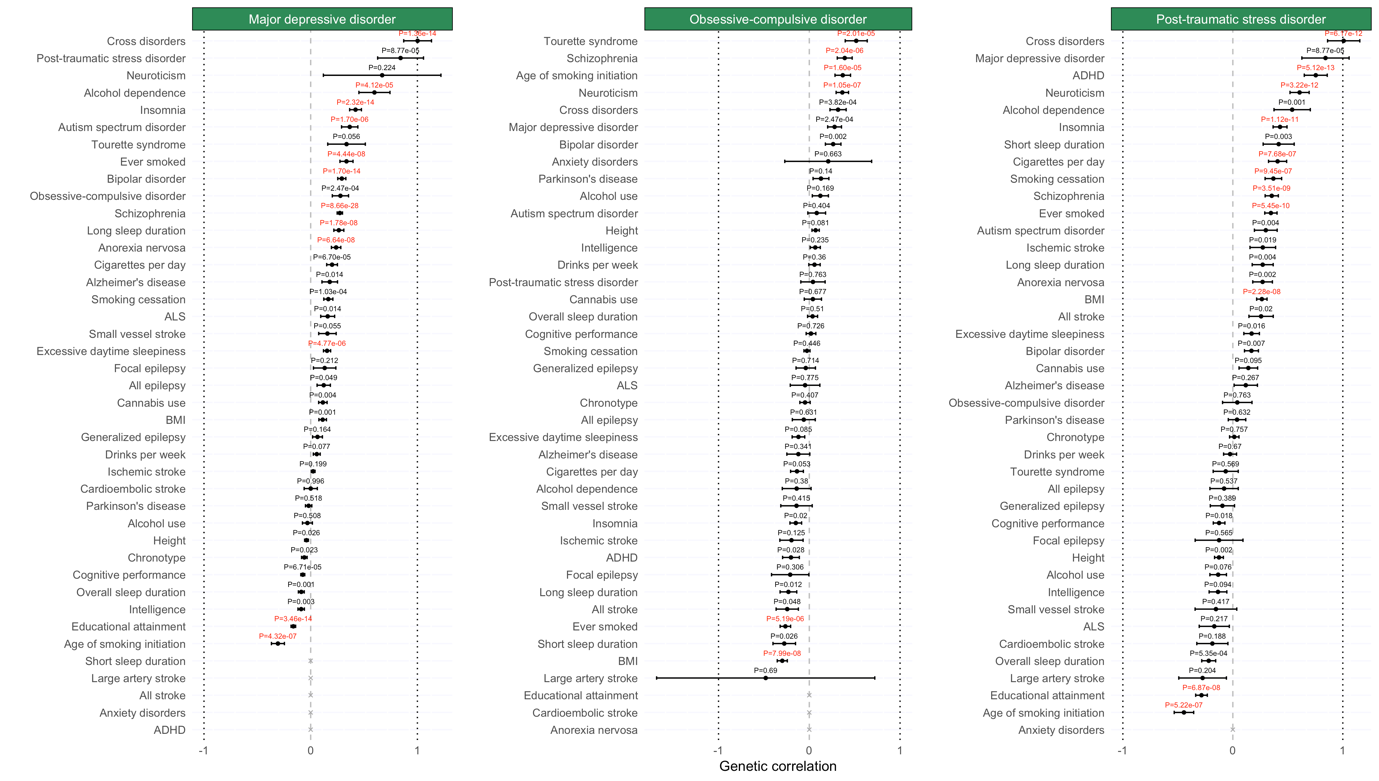

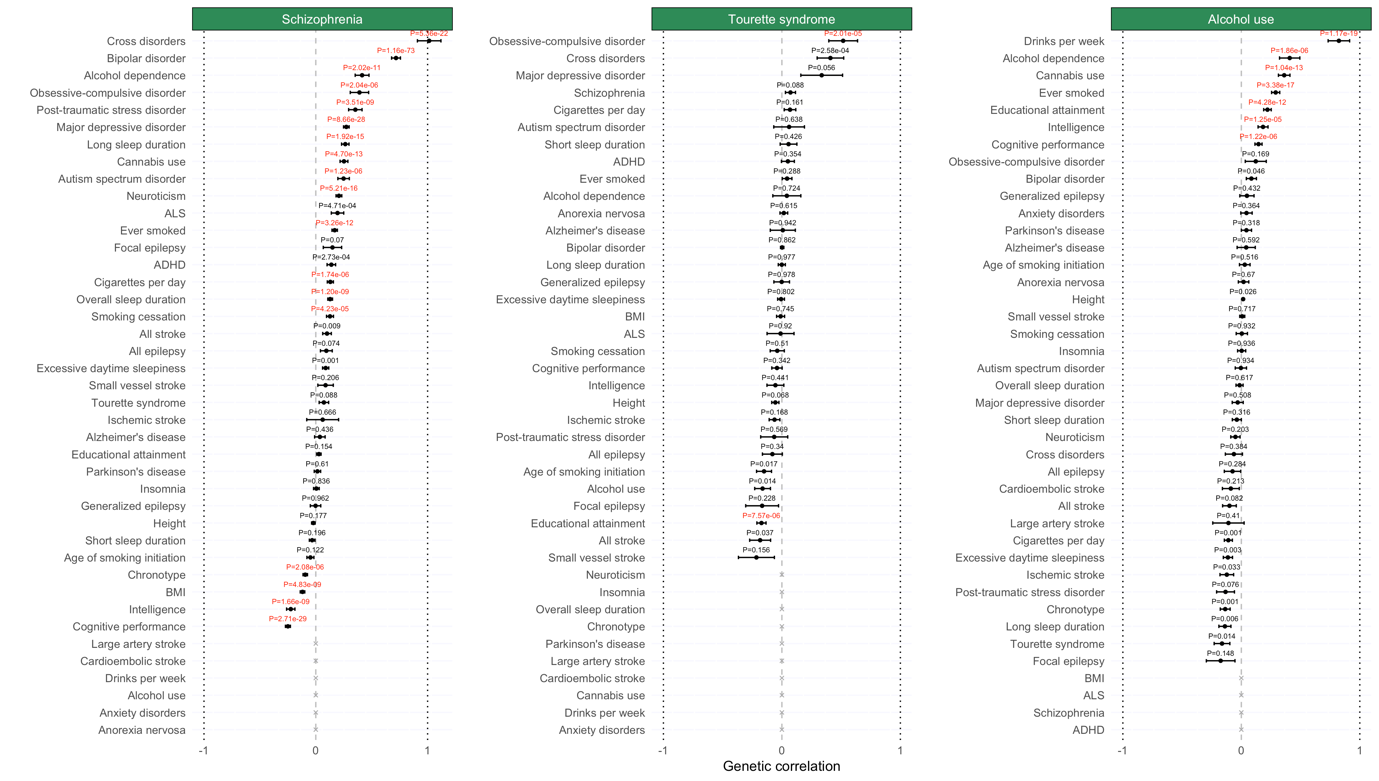

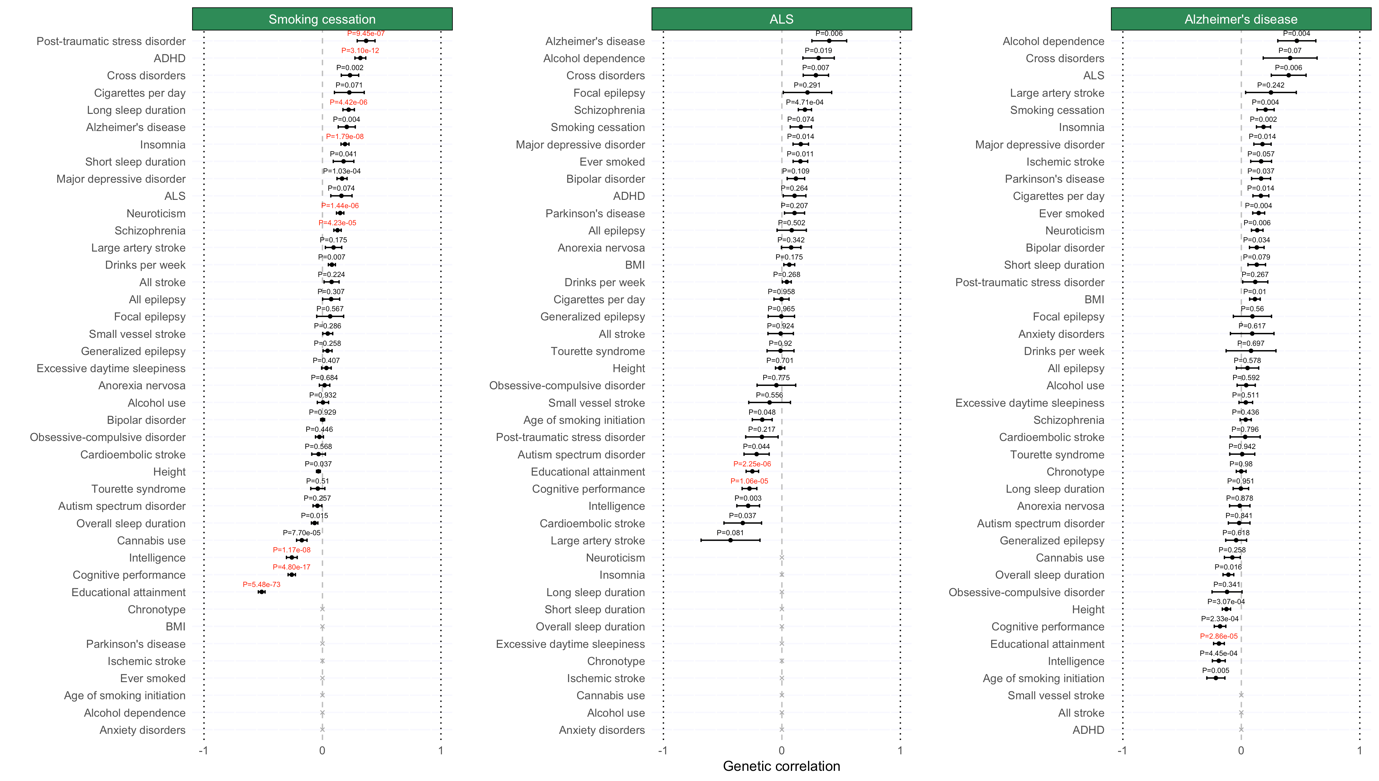

**Figure S13. Cell type enrichment estimated by DEPICT, LDSC, MAGMA (top 10% mode) and MAGMA (linear mode) of all traits using 10x Genomics data.**

*The red line indicates the Bonferroni threshold (P<0.05/(16*42)). The red line is solid if any of the methods identified any cell type as significantly associated, and if none of the methods identified any of the cell types as significantly associated, the red line is dashed.*

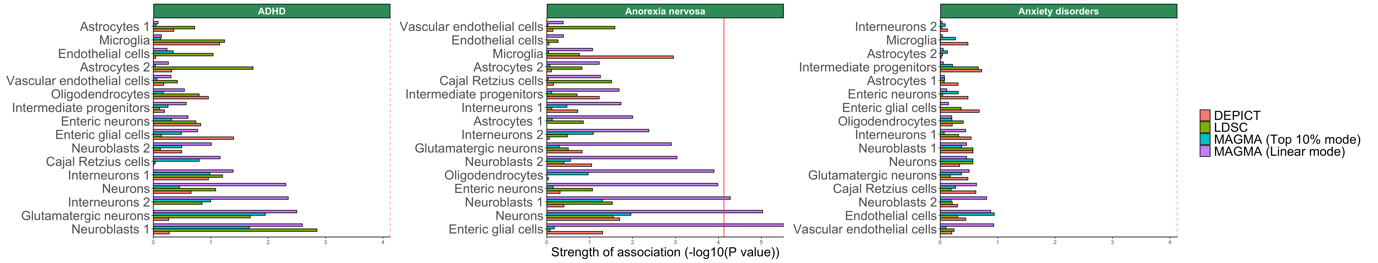

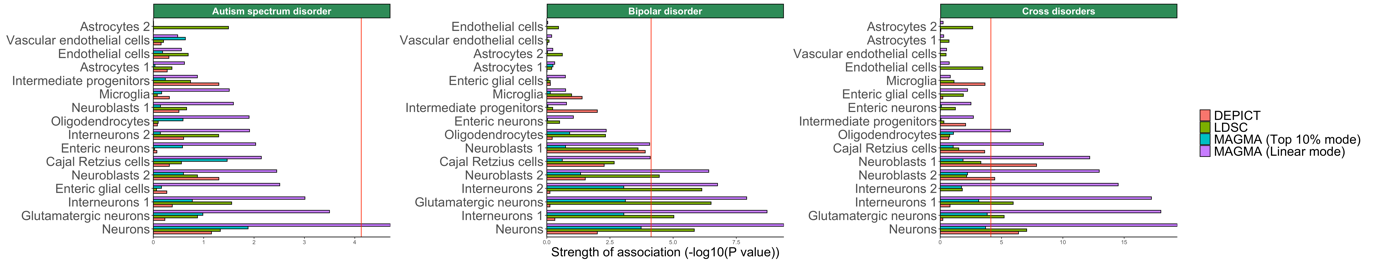

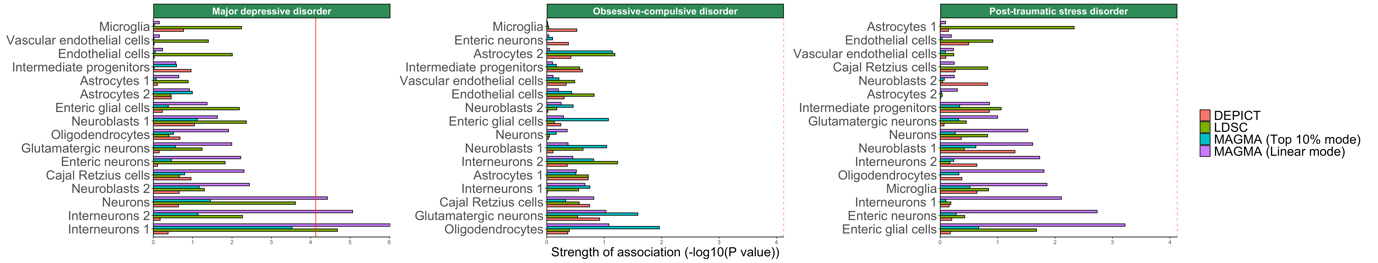

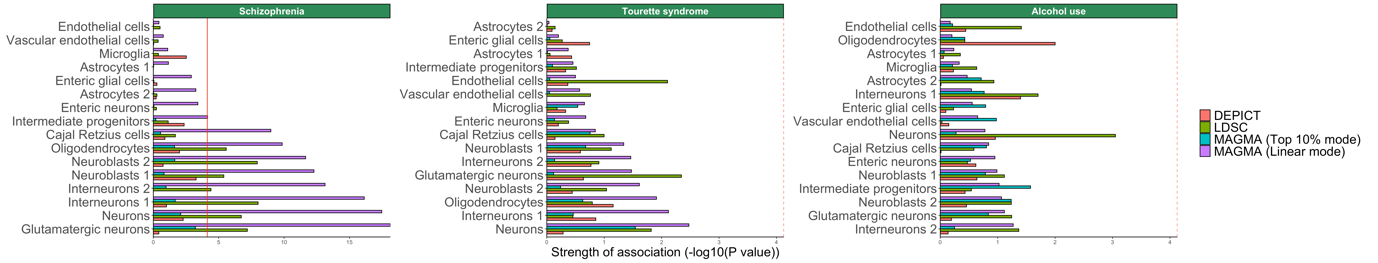

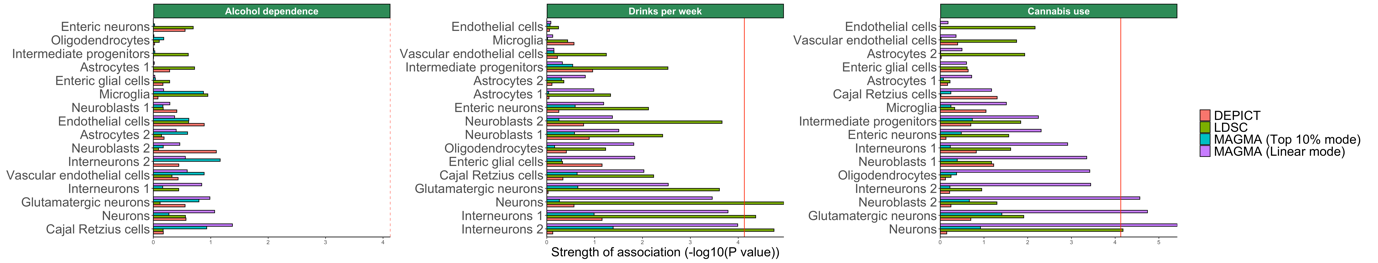

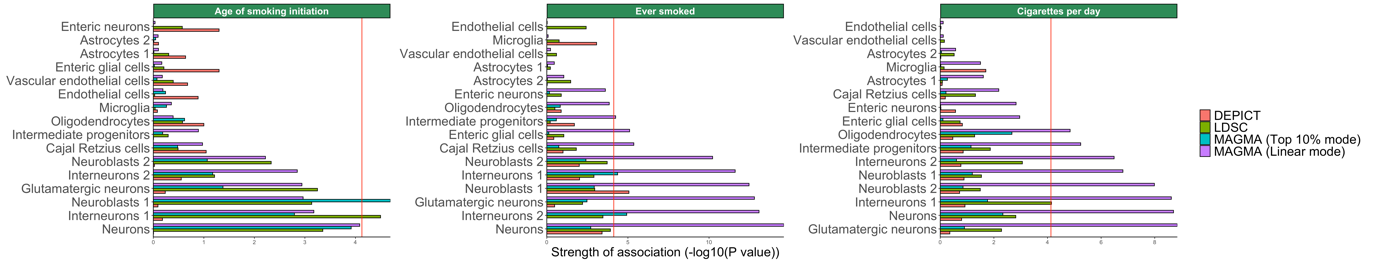

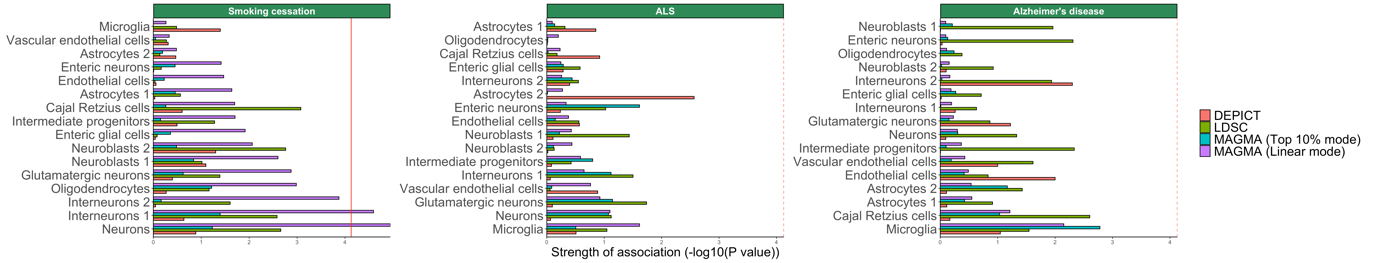

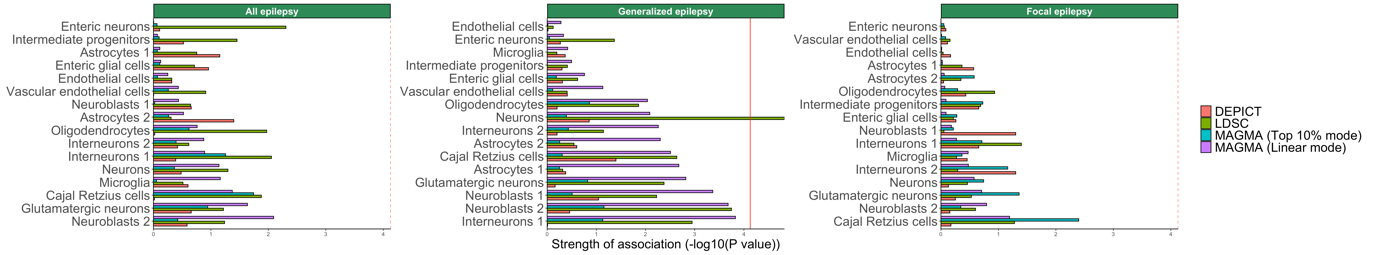

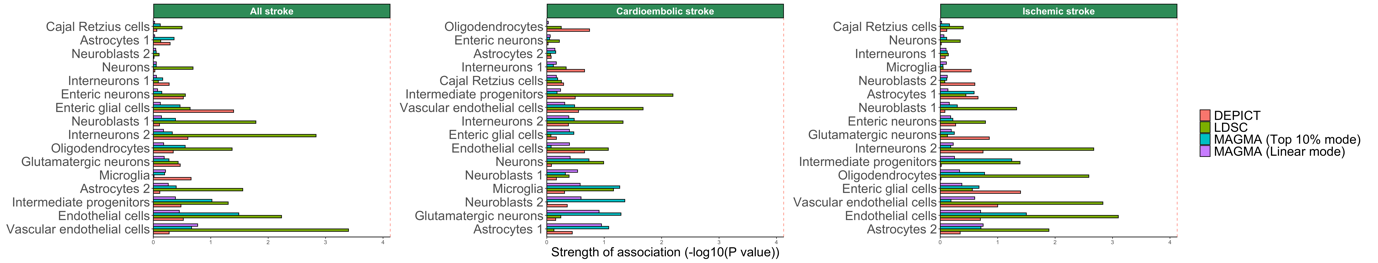

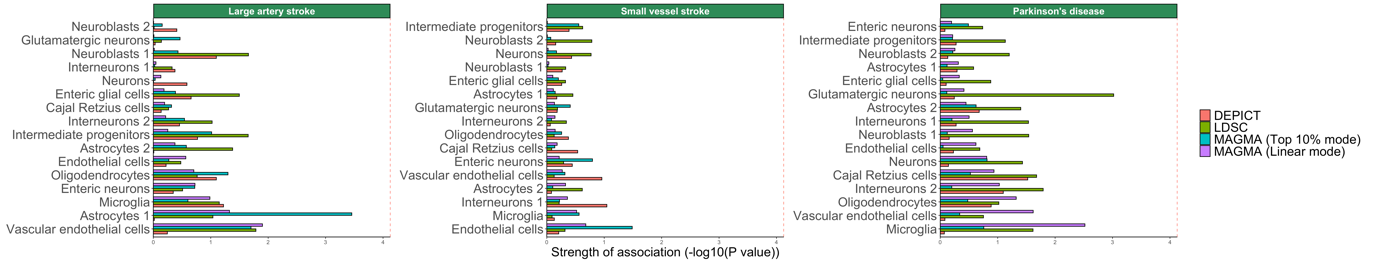

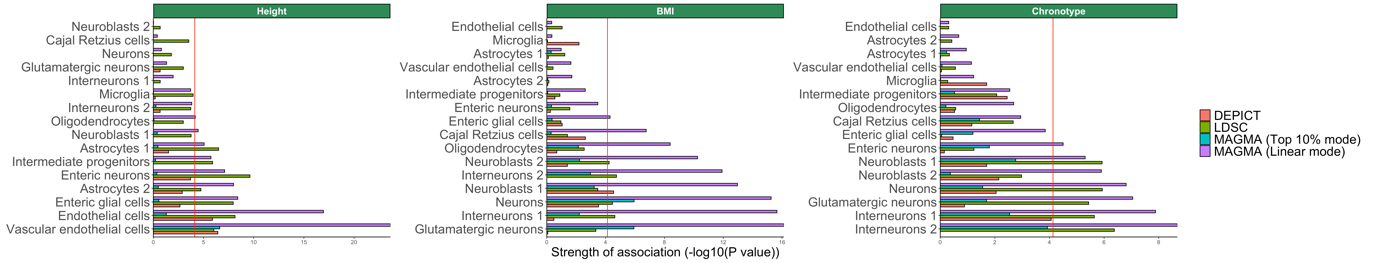

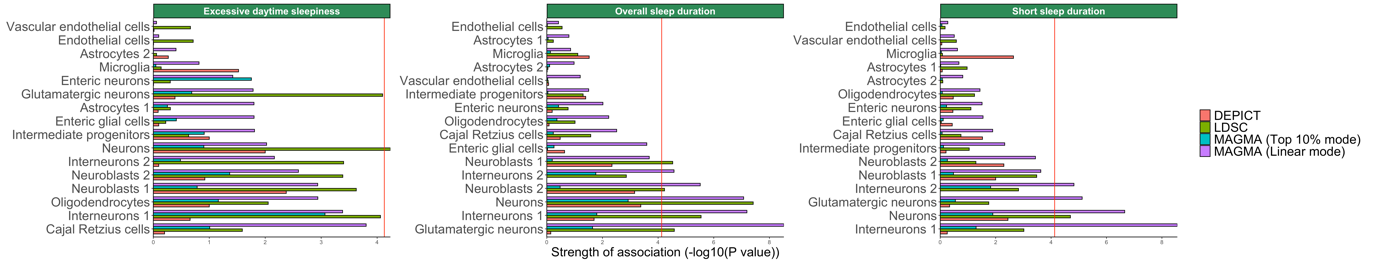

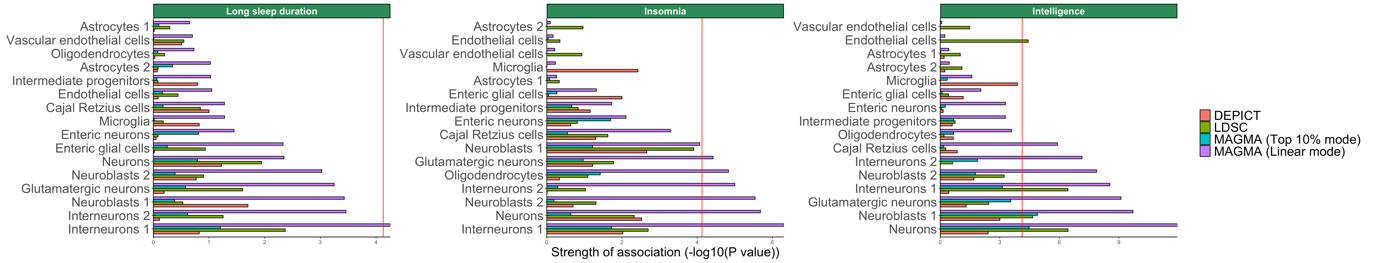

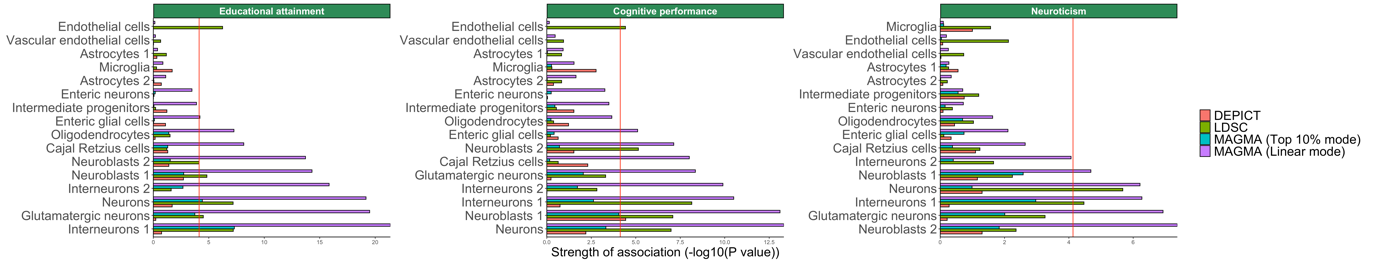

**Figure S14. Overview of enriched cell types of 42 common-variant psychiatric, neurologic and behavioral/quantitative GWAS results in the 10x Genomics dataset.**

*Abbreviations: ADHD; attention deficit hyperactivity disorder, ALS; amyotrophic lateral sclerosis, BMI; body mass index.*

*Analyses from LDSC, DEPICT and MAGMA (top 10% mode), referred to as ‘methods’ in the graph, show enrichment in certain interneurons, certain neuroblasts, neurons, excitatory glutamatergic neurons and endothelial cells. Phenotypes and cell types are grouped by hierarchal clustering. Shades of pink are proportional to the mean strength of association (-log10(P)) of all methods. The color of the frames refers to the number of methods that identified a given cell type as significant in a given phenotype, after Bonferroni correction (P<0.05/(16*42)). Frames: grey; one method (chronotype, BIP, overall sleep duration, SCZ, educational attainment, short sleep duration, age of smoking initiation, cigarettes per day, MDD, generalized epilepsy, ever smoked, neuroticism, excessive daytime sleepiness, drinks per week, cannabis use), black; two methods (cross-disorders, intelligence, cognitive performance and BMI), red; all three methods (human height).*

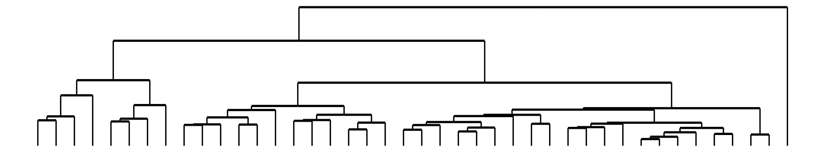

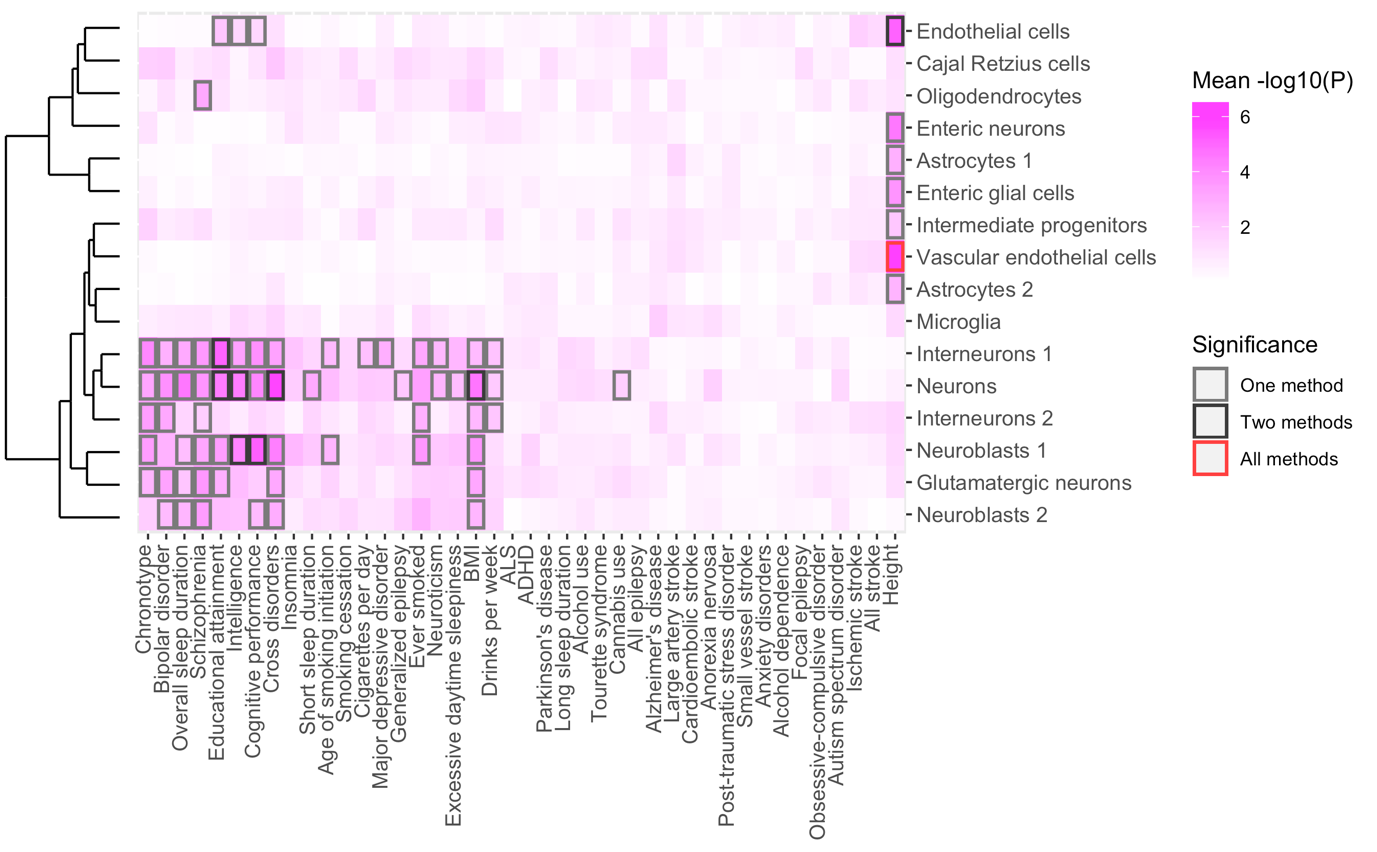

**Figure S*15*. Overview of enriched cell types of 42 common-variant psychiatric, neurologic and behavioral/quantitative GWAS results in the 10x Genomics dataset, including MAGMA in linear mode.**

*Abbreviations: ADHD; attention deficit hyperactivity disorder, ALS; amyotrophic lateral sclerosis, BMI; body mass index.*

*Analyses from LDSC, DEPICT, MAGMA (top 10% mode) and MAGMA (linear mode) show enrichment in certain interneurons, certain neuroblasts, neurons, excitatory glutamatergic neurons and endothelial cells. Phenotypes and cell types are grouped by hierarchal clustering. Shades of pink are proportional to the mean strength of association (-log10(P)) of all methods. The colour of the frames refers to the number of methods that identified a given cell type as significant in a given phenotype, after Bonferroni correction (P<0.05/(16*42)). Frames: light grey; one method (insomnia, age of smoking initiation, smoking cessation, generalized epilepsy, excessive daytime sleepiness, drinks per week, long sleep duration, anorexia nervosa, ASD), dark grey; two methods (chronotype, BIP, overall sleep duration, SCZ, short sleep duration, cigarettes per day, MDD, ever smoked, neuroticism, cannabis use), black; three methods (educational attainment, intelligence, cognitive performance, cross-disorders, BMI), red; all four methods (human height).*

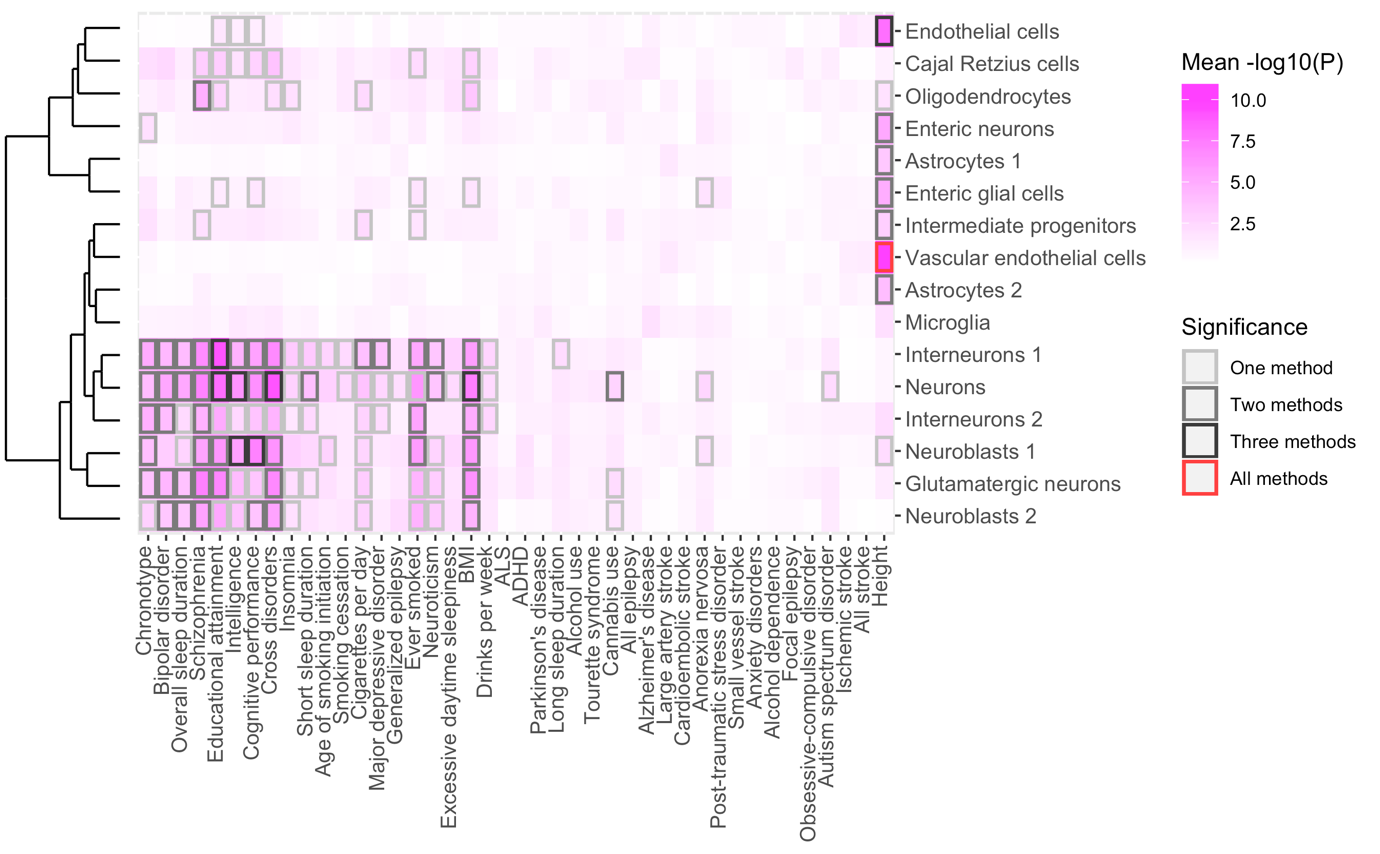

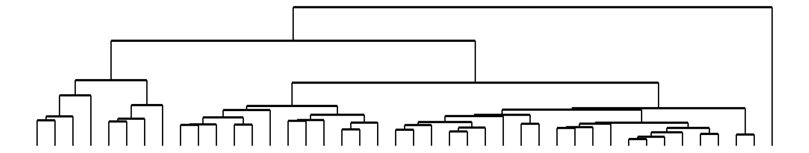

**Figure S16. Comparison of P-values computed by LDSC, MAGMA and DEPICT in the 10x Genomics dataset.**

*Axes are -log10(P) transformed P-values from LDSC, MAGMA (top 10%), MAGMA (linear) and DEPICT. Dots represent cell types within the 10x Genomics dataset. The blue line represents a linear regression and the area coloured in grey represents the standard error of the linear regression.*

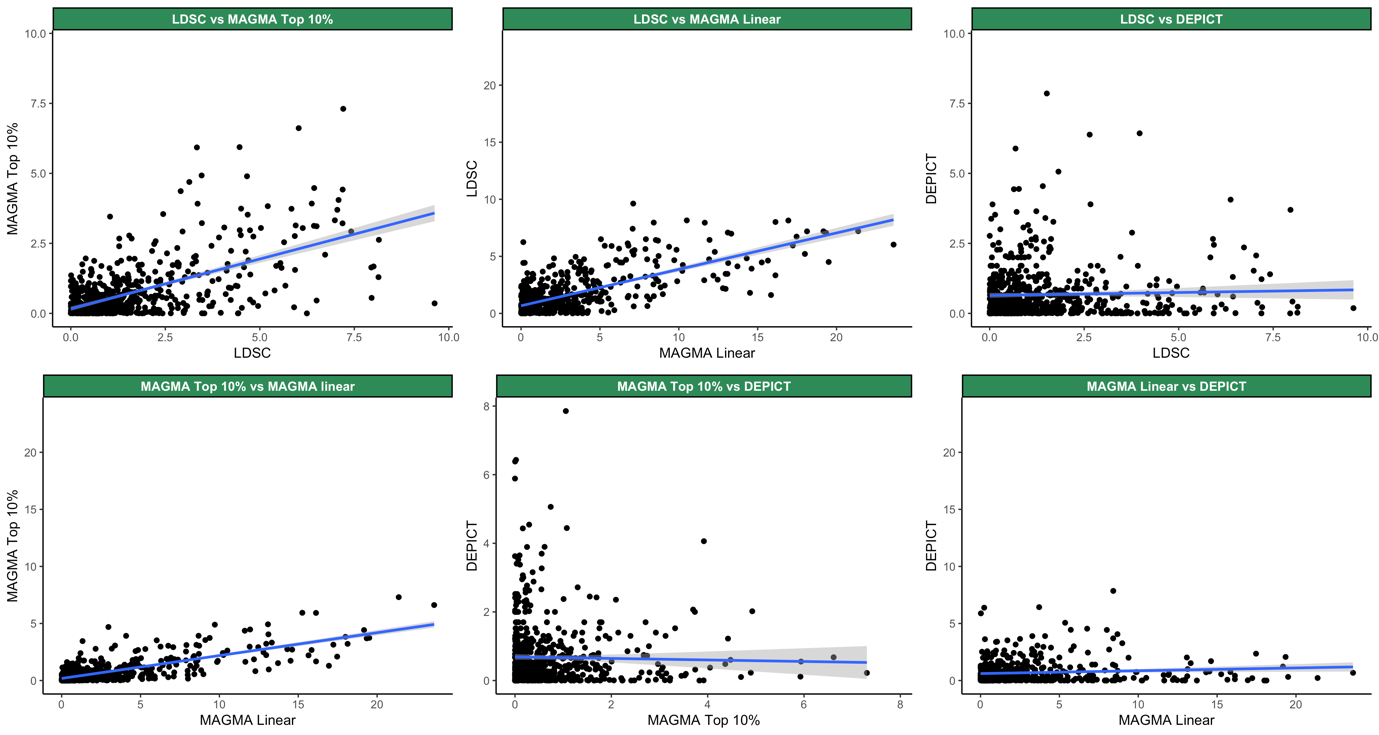

**Figure S17. Correlations between the top associated 10x Genomics cell types with GWAS sample size.**

*Abbreviations: SUD; substance-use disorder.*

*The strength of association reflects the association of the top associated cell type with a given phenotype. The strongest association, out of LDSC, DEPICT, MAGMA (top 10% mode) and MAGMA (linear mode), was selected to denote the strength of association. The grey line represents a linear regression and the area coloured in a lighter shade of grey represents the standard error of the linear regression.* ***A.*** *Correlation between the top associated cell type, out of LDSC, DEPICT, MAGMA (top 10% mode) and MAGMA (linear mode), and sample size.* ***B.*** *Correlation between the top associated cell type, out of all methods excluding MAGMA (linear mode), and sample size.*

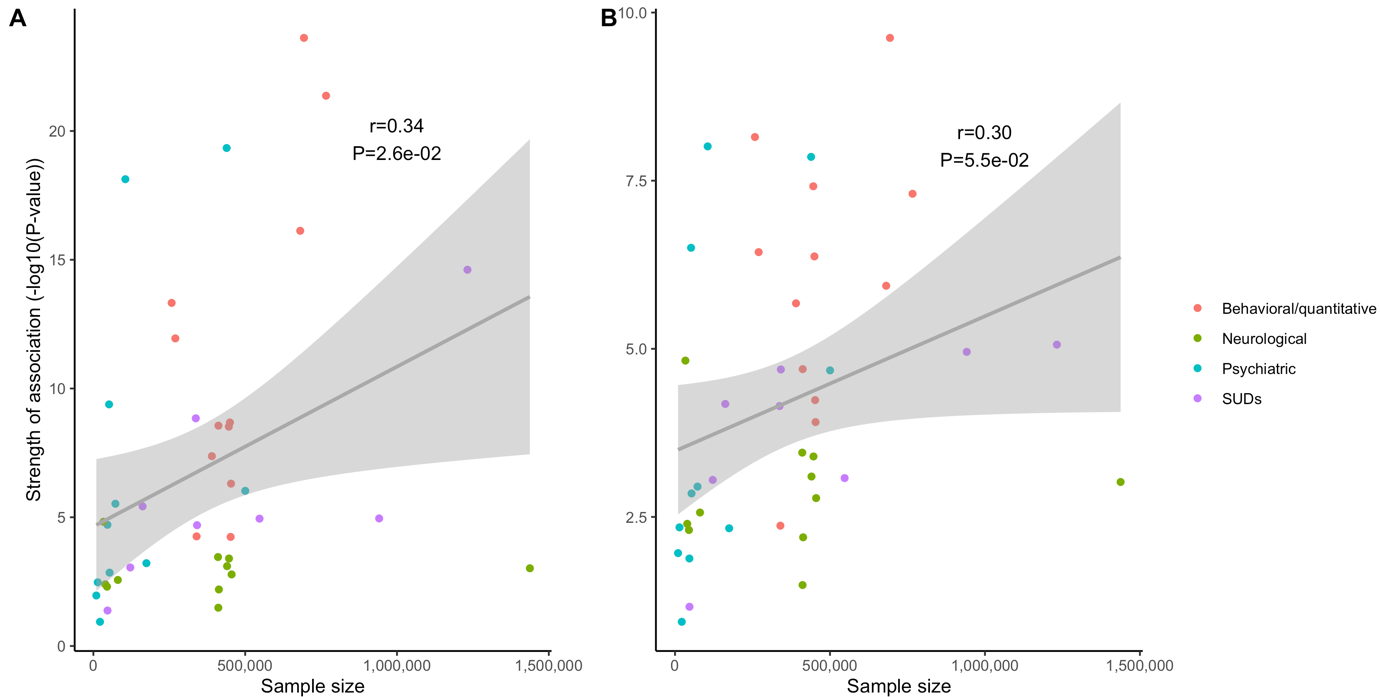

**Figure S18. Cell type enrichment estimated by DEPICT, LDSC, MAGMA (top 10% mode) and MAGMA (linear mode) of all traits using KI data.**

*The red line indicates the Bonferroni threshold (P<0.05/(24*42)). The red line is solid if any of the methods identified any cell type as significantly associated, and if none of the methods identified any of the cell types as significantly associated, the red line is dashed.*

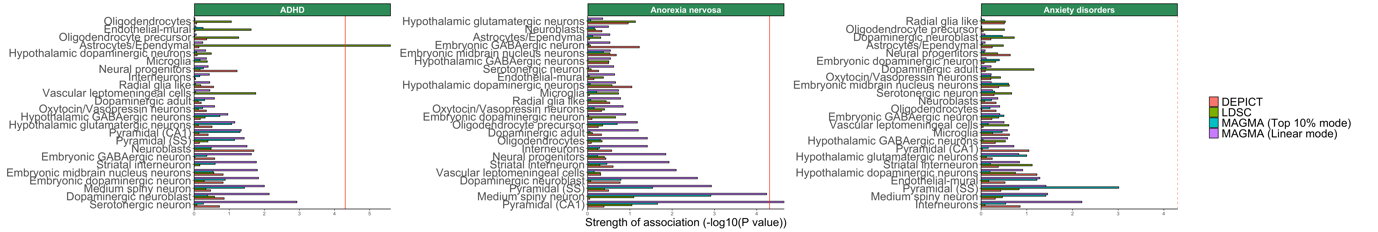

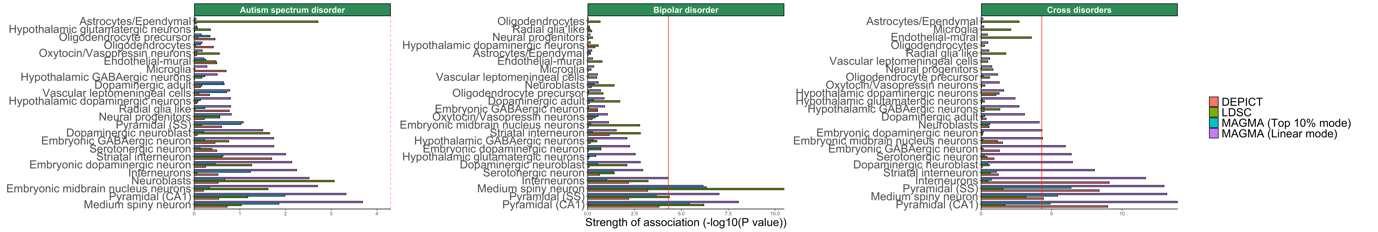

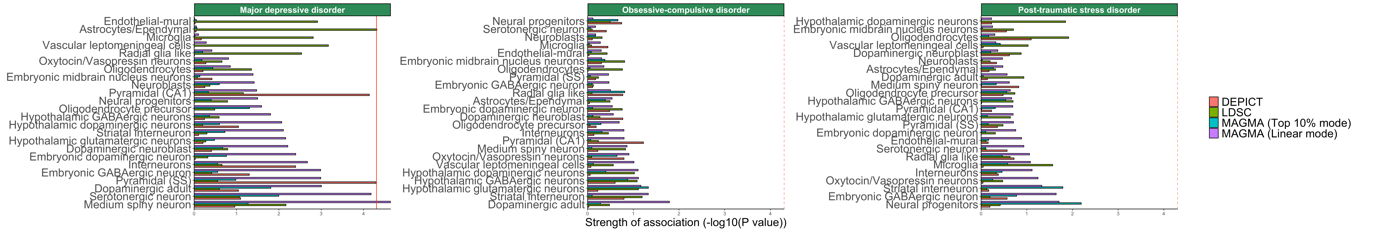

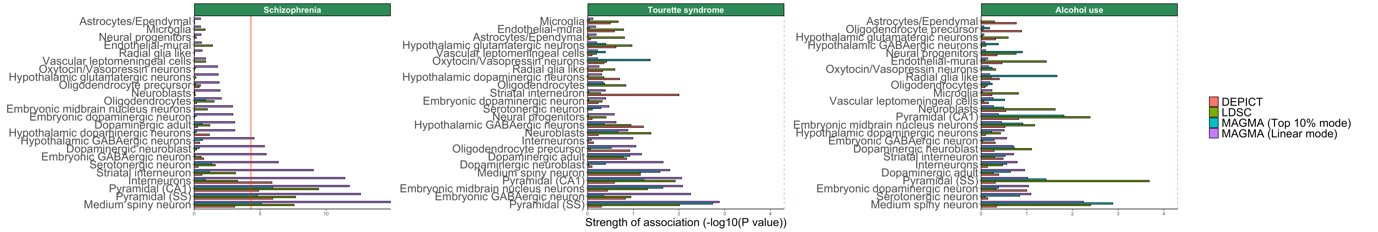

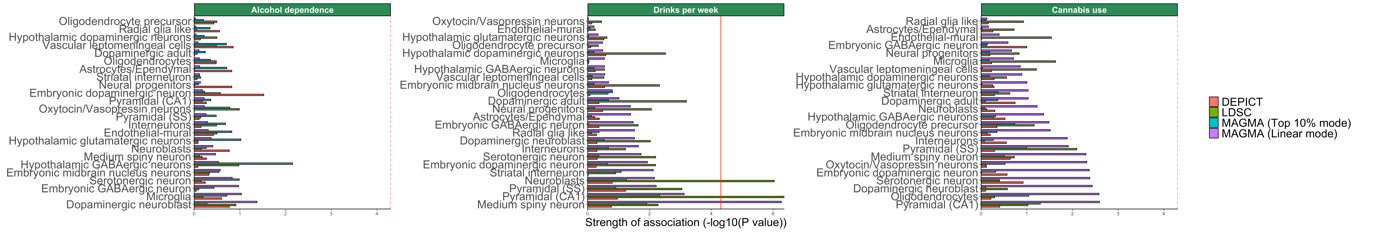

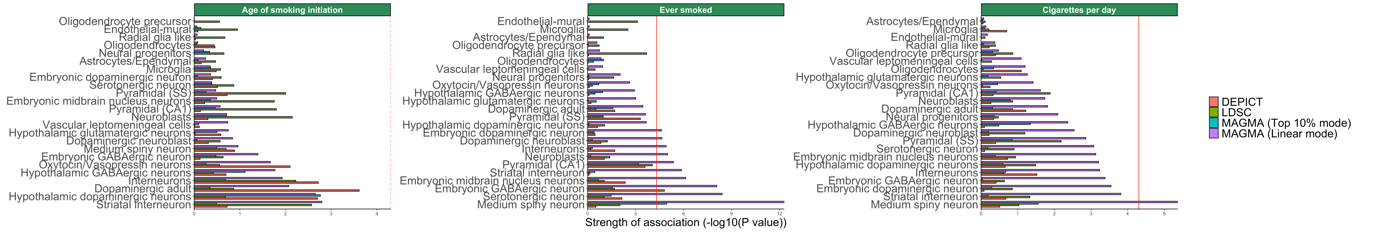

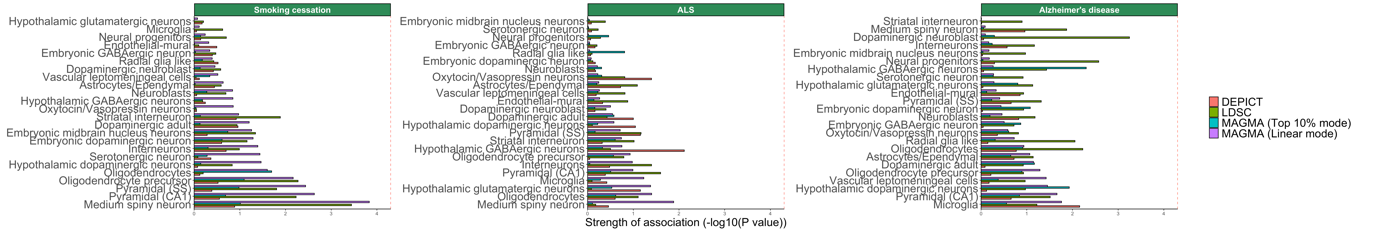

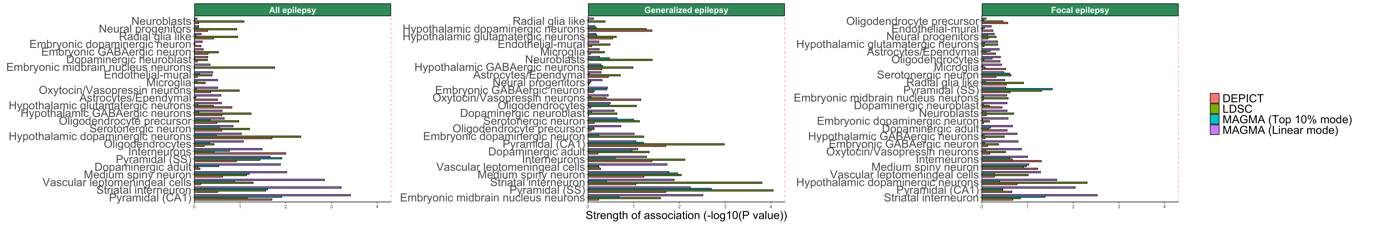

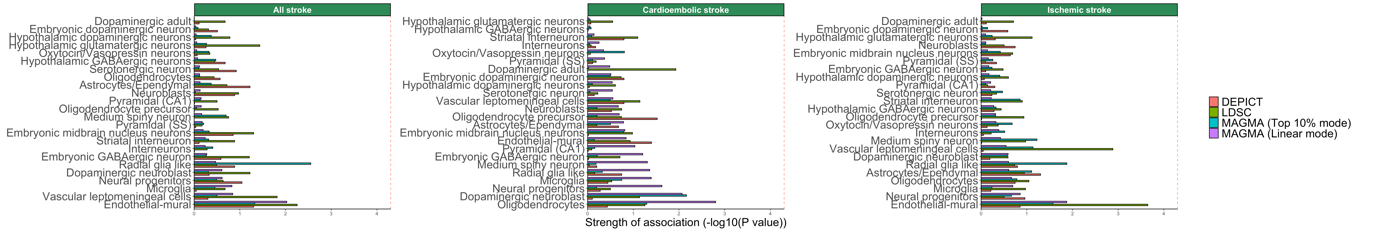

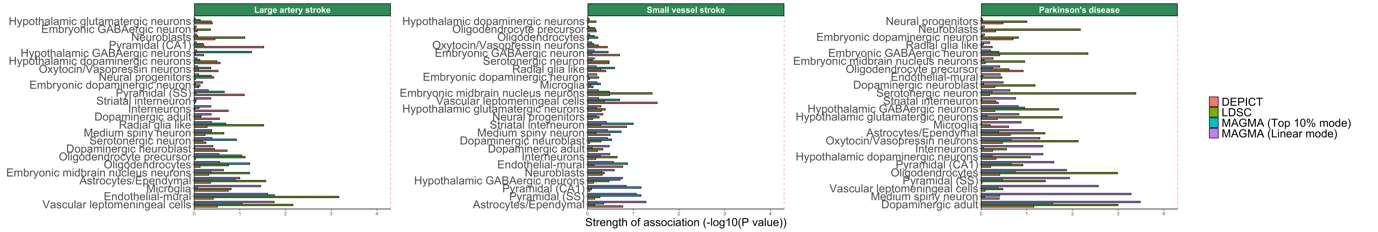

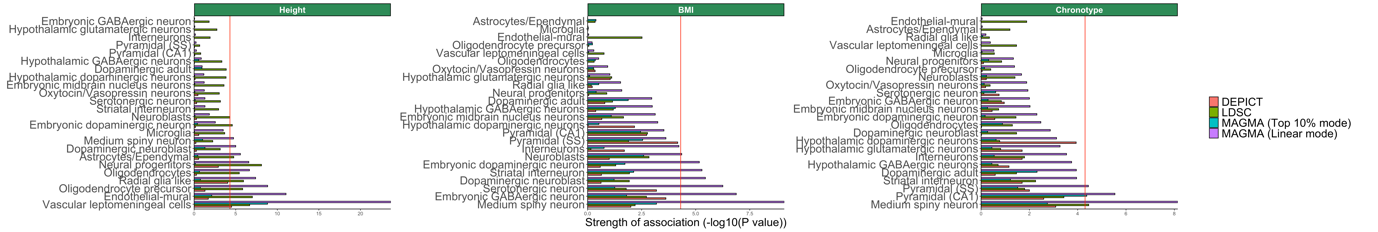

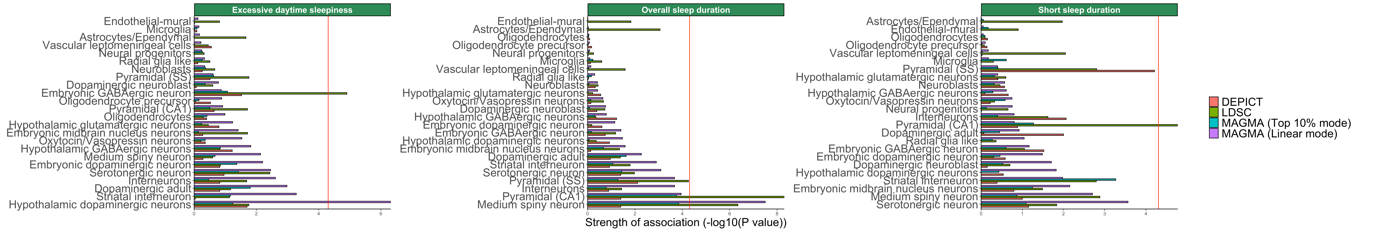

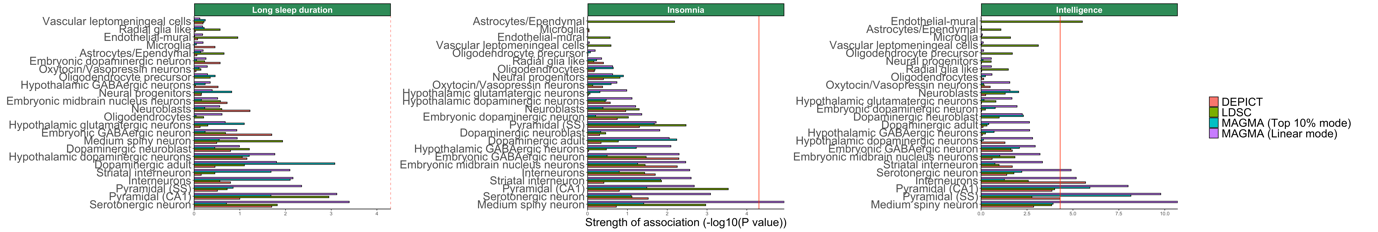

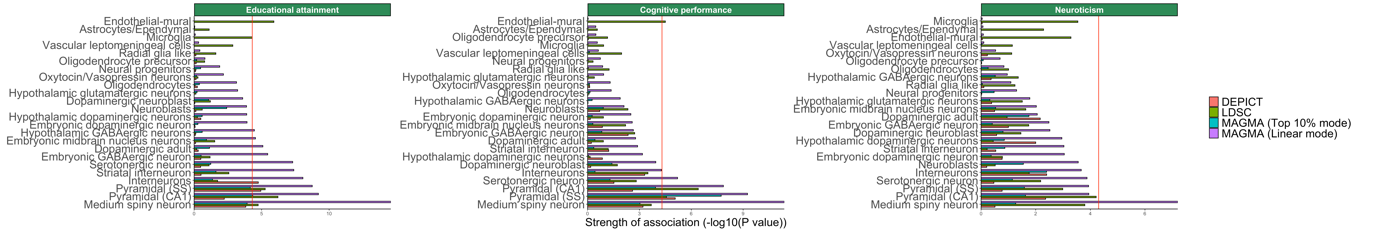

**Figure S19. Overview of enriched cell types of common-variant GWAS results in the KI dataset.**

*Abbreviations: ADHD; attention deficit hyperactivity disorder, ALS; amyotrophic lateral sclerosis, BMI; body mass index.*

*Analyses from LDSC, DEPICT, MAGMA (top 10% mode) and MAGMA (linear mode) consistently show enrichment in MSNs and pyramidal cells (CA1) and pyramidal cells (SS) across brain-related phenotypes. Phenotypes and cell types are grouped by hierarchal clustering. Shades of pink are proportional to the mean strength of association (-log10(P)) of all methods. The colour of the frames refers to the number of methods that identified a given cell type as significant in a given phenotype, after Bonferroni correction (P<0.05/(24*42)). Frames: light grey; one method (excessive daytime sleepiness, cigarettes per day, anorexia nervosa, ADHD, drinks per week, BMI, short sleep duration, insomnia, MDD, neuroticism), dark grey; two methods (intelligence, ever smoked, chronotype, overall sleep duration), black; three methods (cross-disorders, educational attainment, BIP), red; all four methods (human height, cognitive performance, SCZ.*

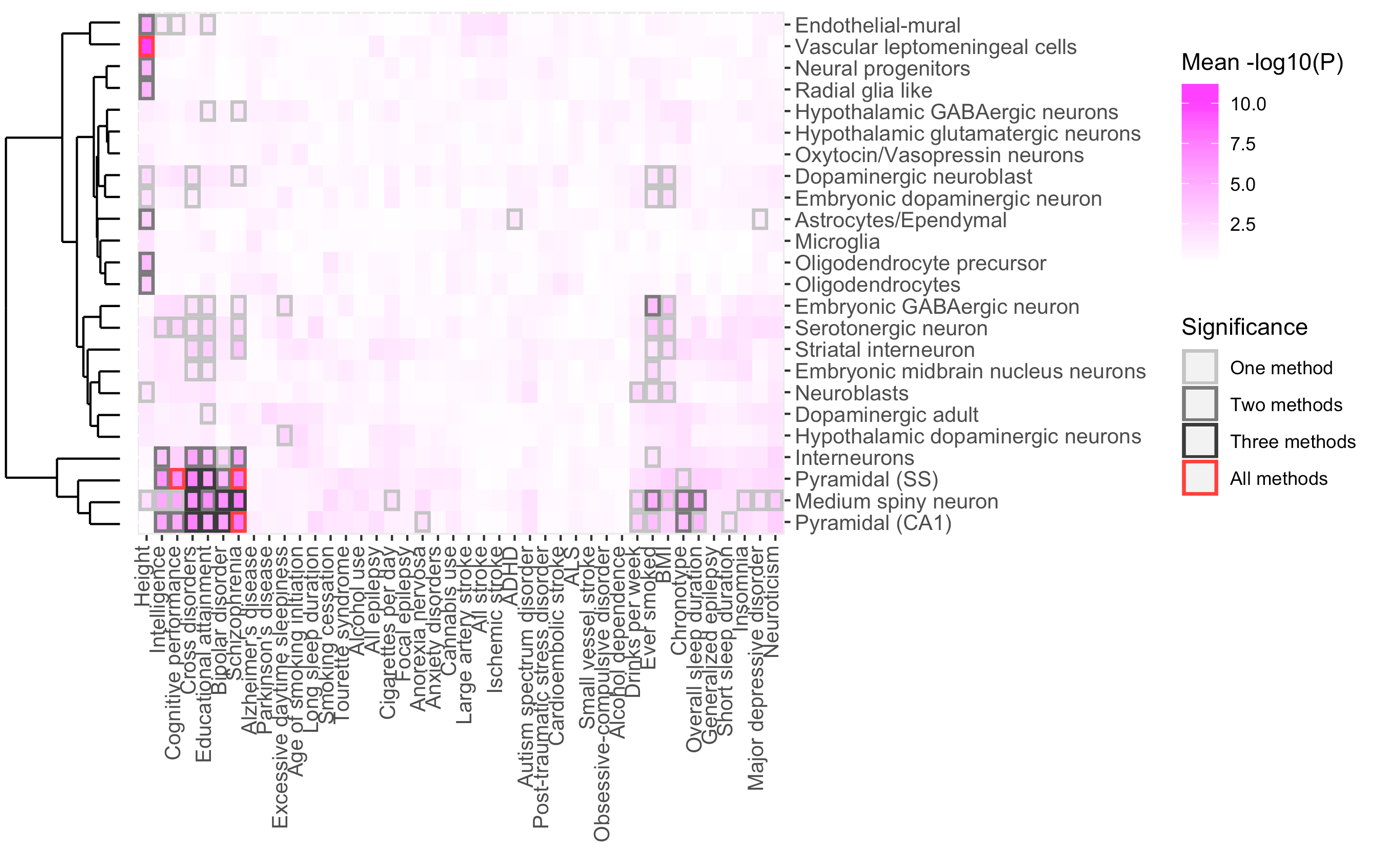

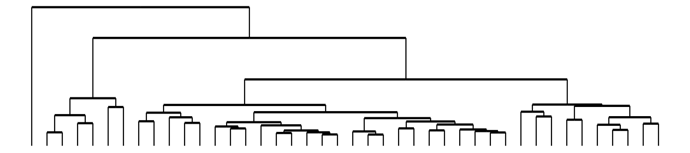

**Figure S20.Enrichment estimated by DEPICT, LDSC, MAGMA (top 10% mode) and MAGMA (linear mode) of all traits using KI level 2 data.**

*The top 50 most associated cell types, out of 149, per phenotype are displayed. The red line indicates the Bonferroni threshold (P<0.05/(149*42)). The red line is solid if any of the methods identified any cell type as significantly associated, and if none of the methods identified any of the cell types as significantly associated, the red line is dashed.*

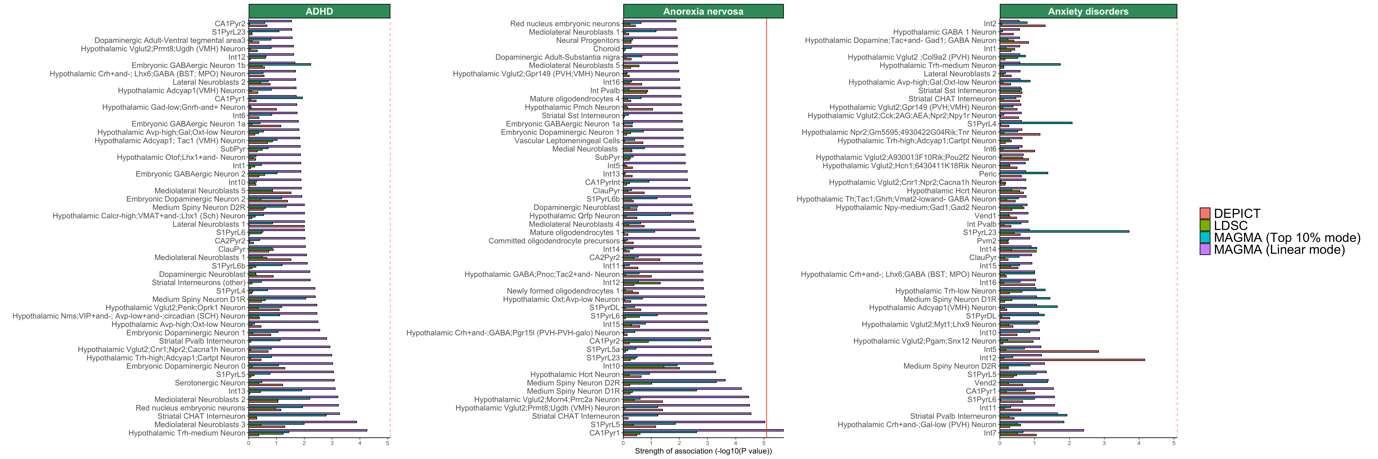

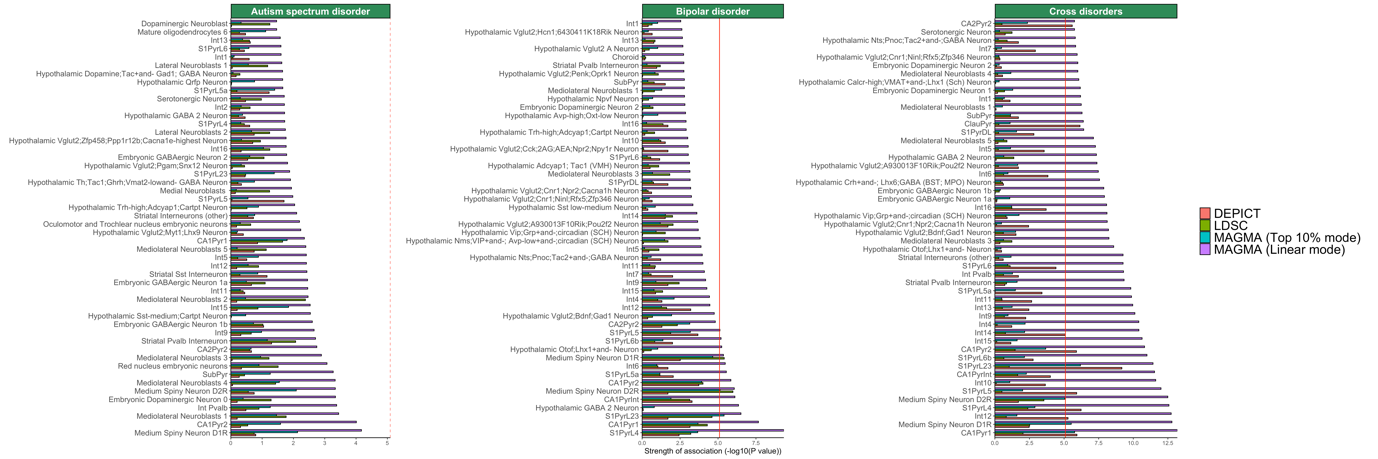

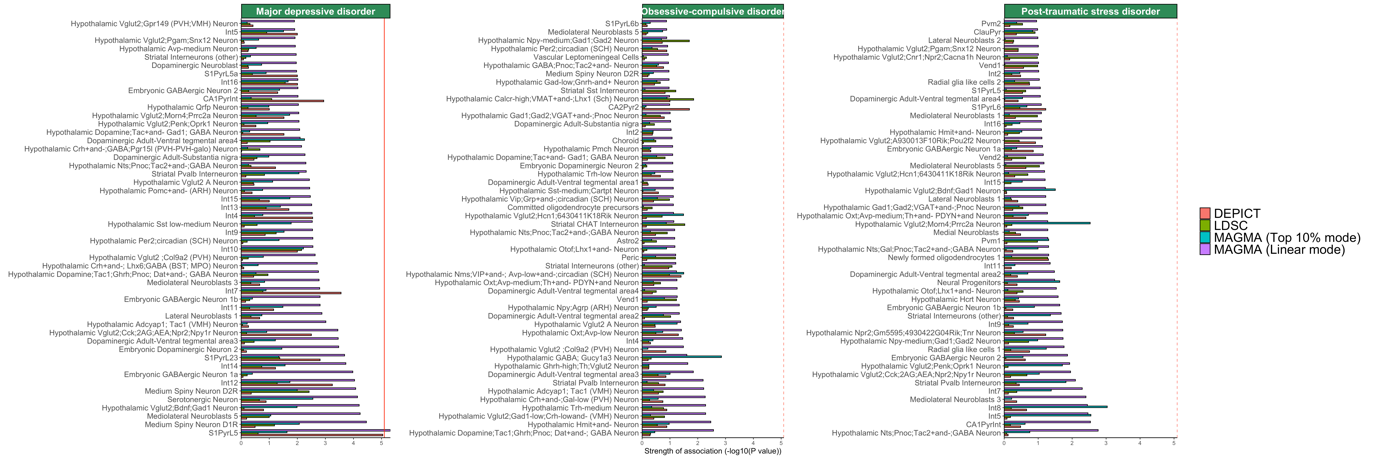

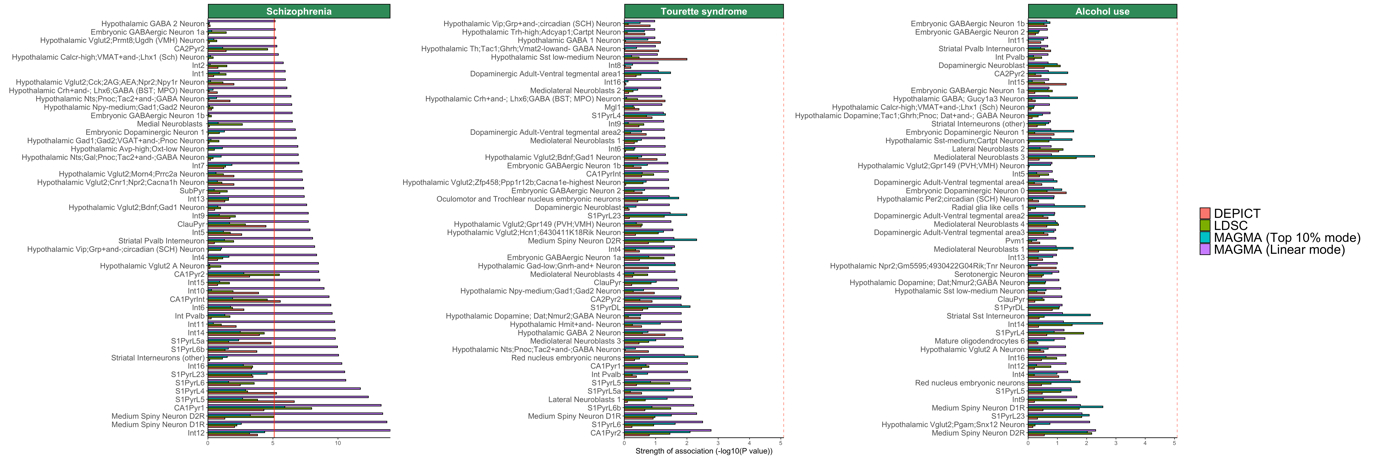

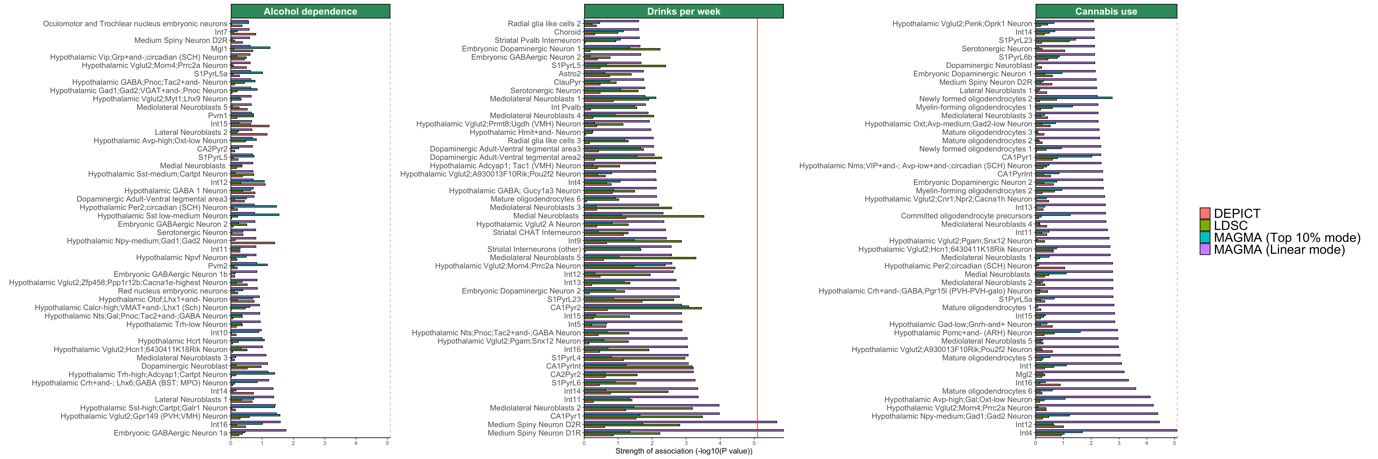

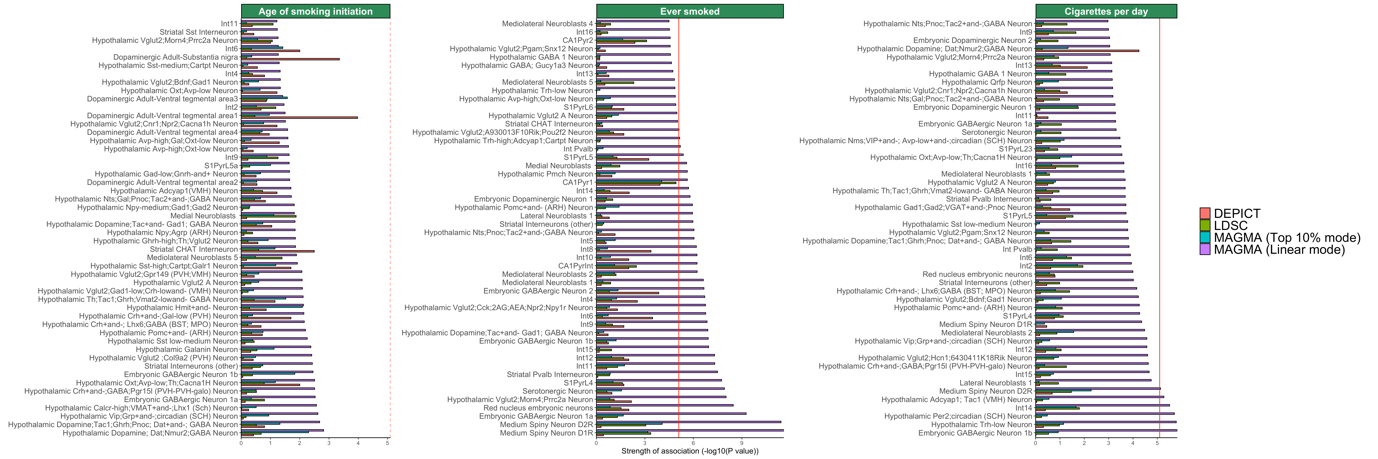

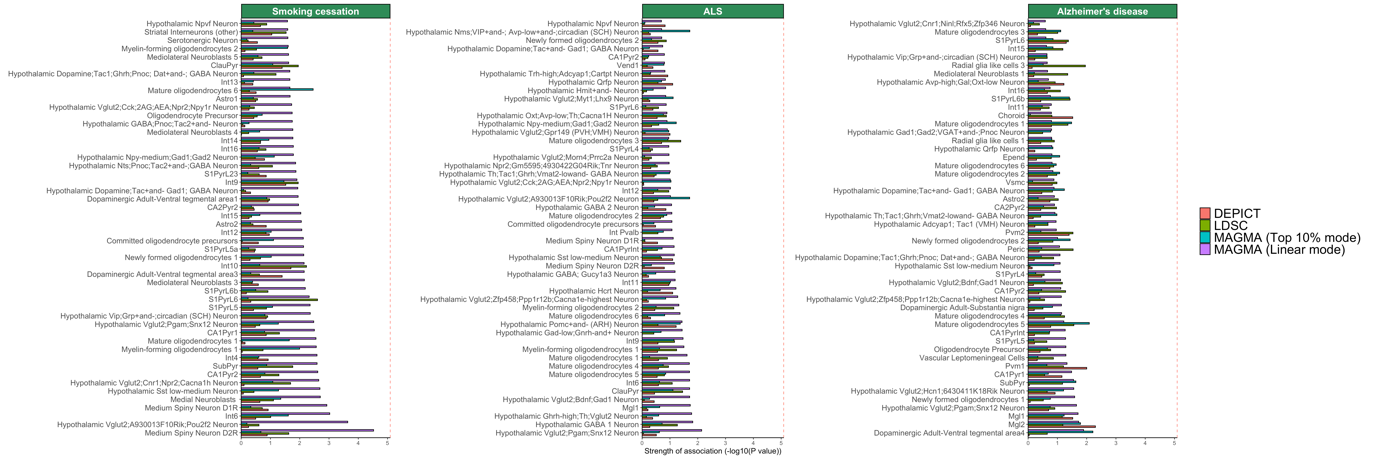

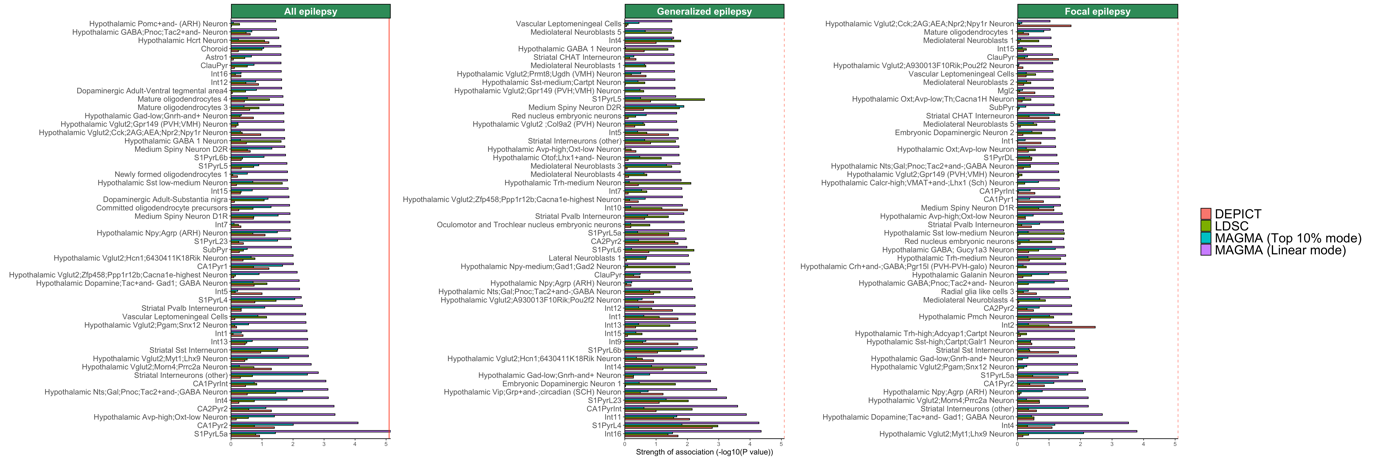

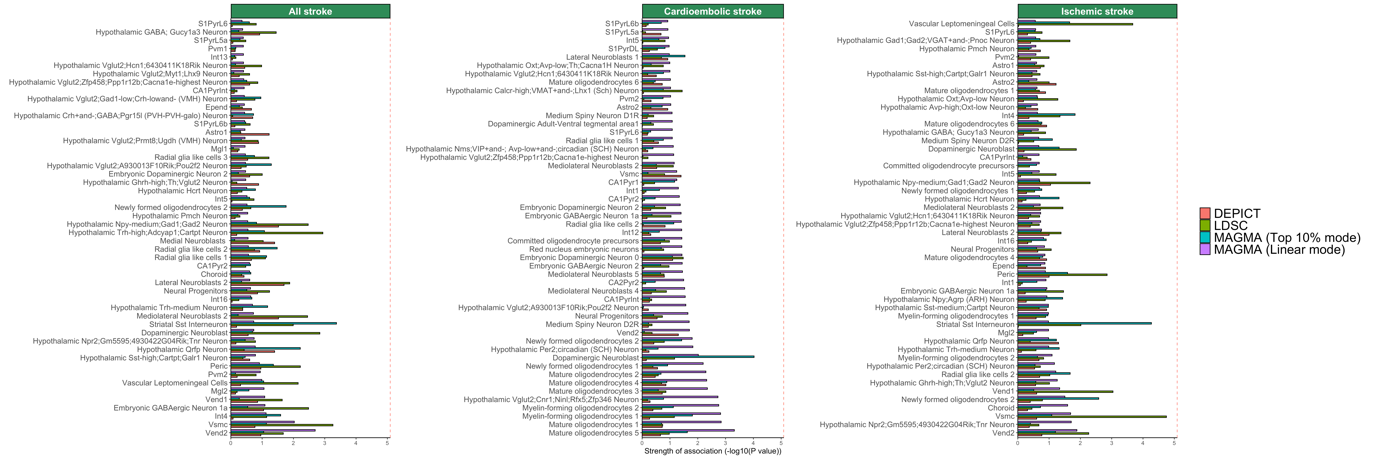

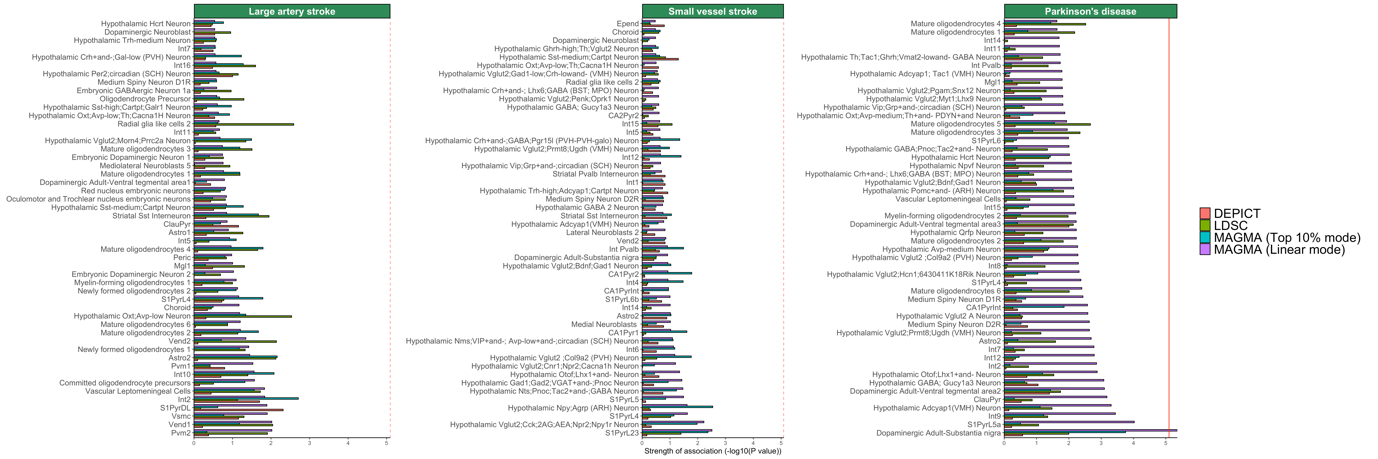

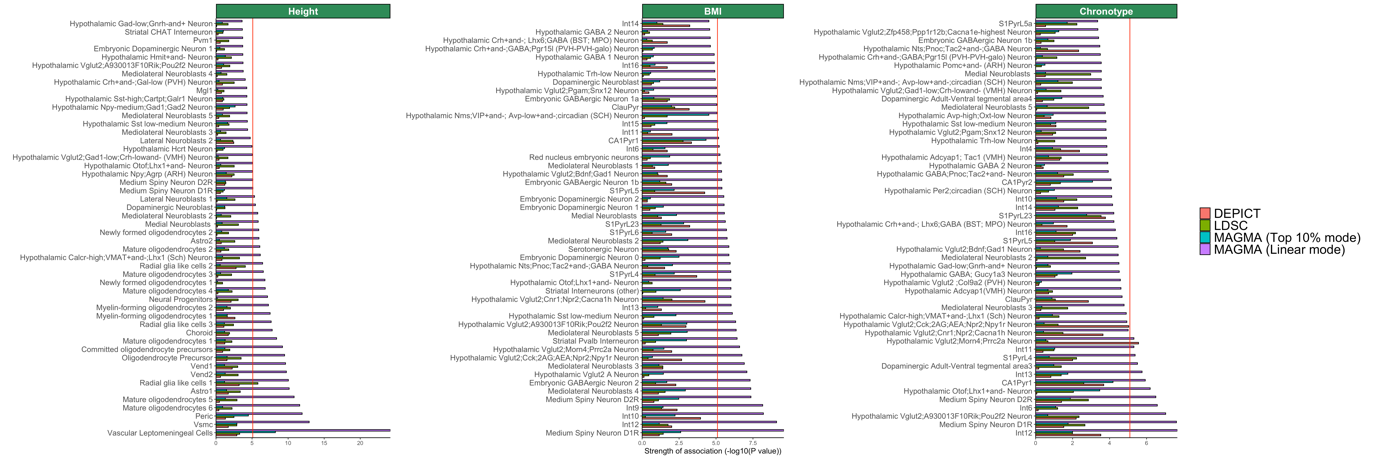

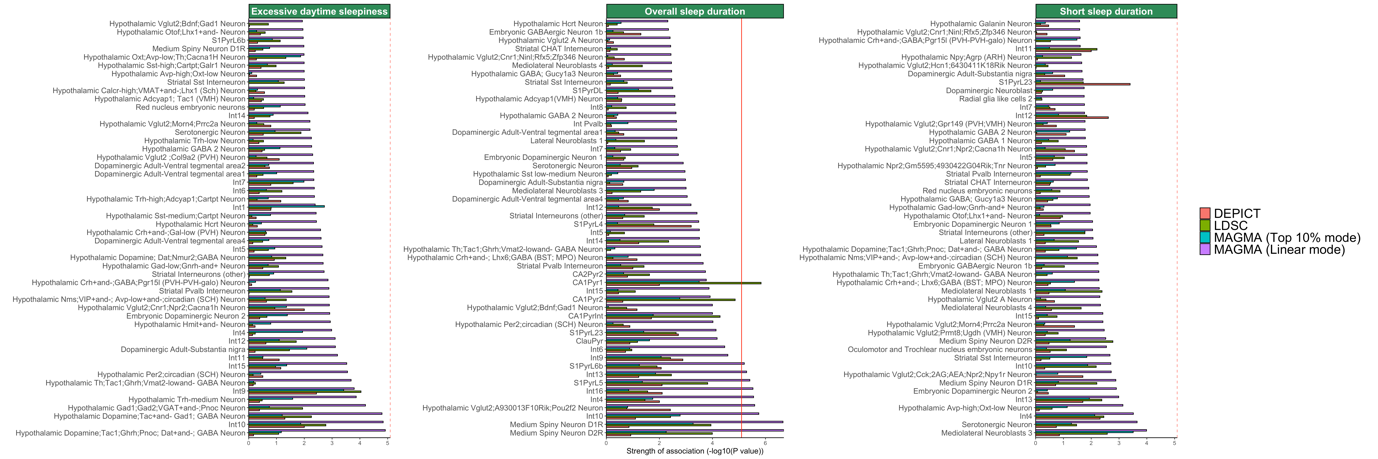

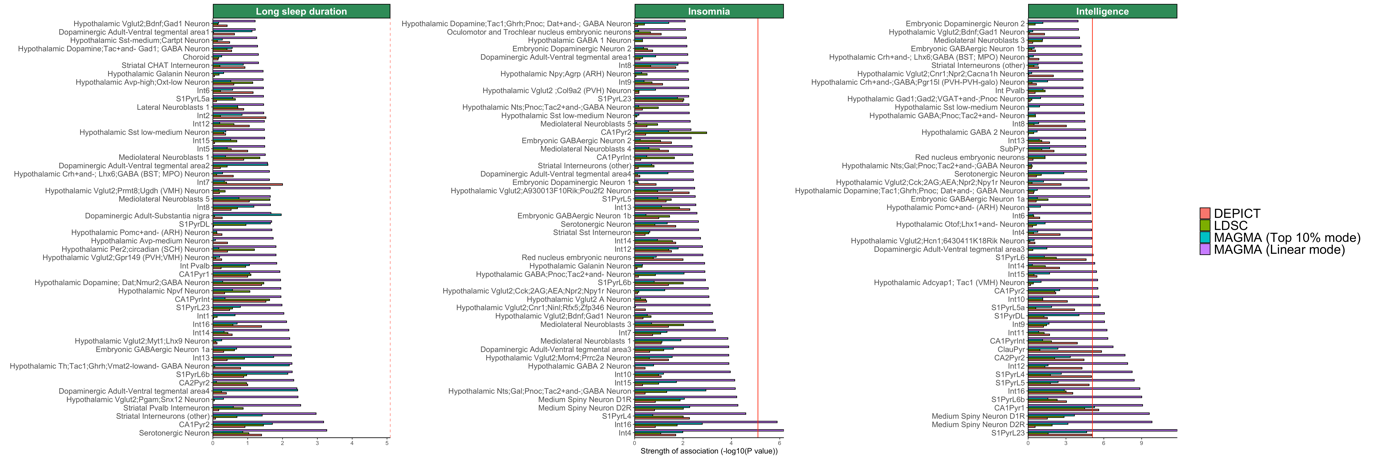

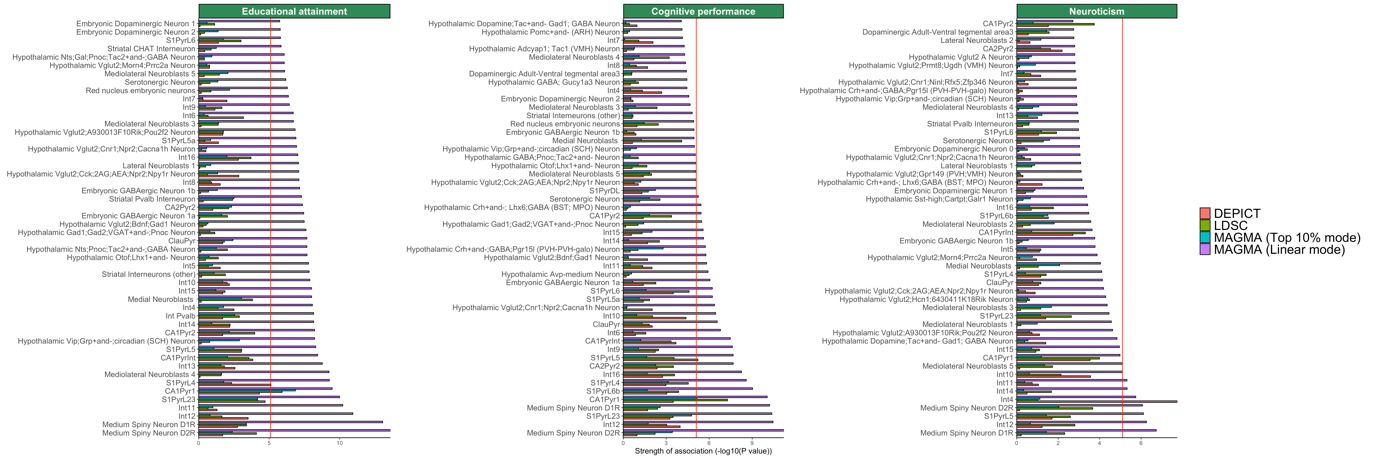

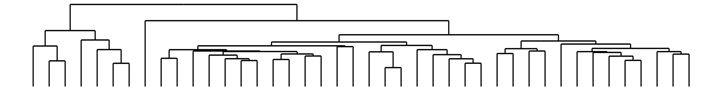

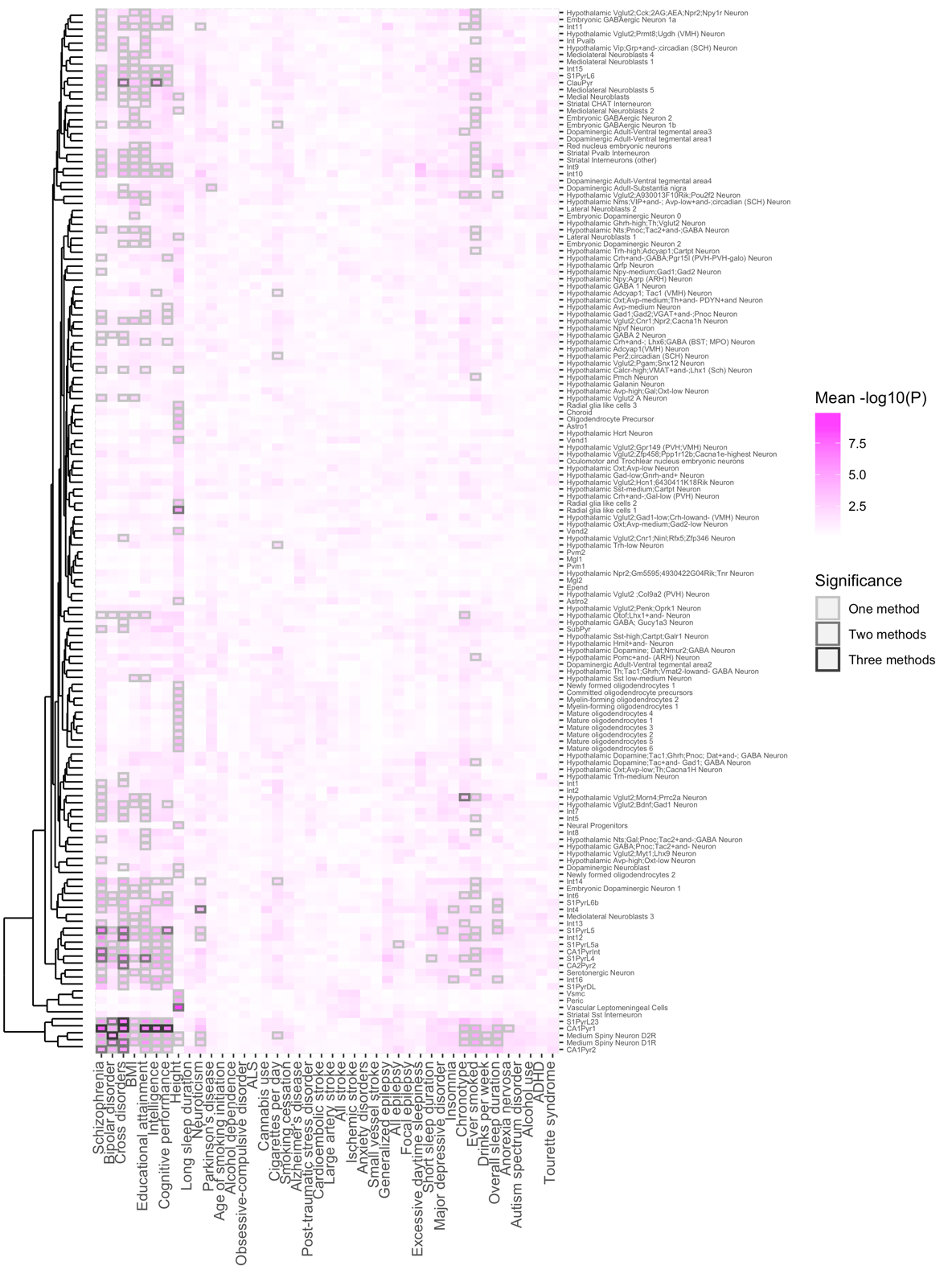

**Figure S21. Overview of enriched cell types of 42 common-variant psychiatric, neurologic and behavioral/quantitative GWAS results in the KI level 2 dataset, including MAGMA in linear mode.**

*Abbreviations: ADHD; attention deficit hyperactivity disorder, ALS; amyotrophic lateral sclerosis, BMI; body mass index.*

Analyses from LDSC, DEPICT, MAGMA (linear mode) and MAGMA (top 10% mode) show mostly enrichment in various excitatory pyramidal cells and certain MSNs. . Phenotypes and cell types are grouped by hierarchal clustering. Shades of pink are proportional to the mean strength of association (-log10(P)) of all methods. The colour of the frames relates to the number of methods that identified a given cell type as significant in a given phenotype, after Bonferroni correction (P<0.05/(149*42)). Frames: light grey; one method (BMI, parkinson’s disease, cigarettes per day, all epilepsy, short sleep duration, MDD, insomnia, ever smoked, drinks per week, overall sleep duration, anorexia nervosa, dark grey; two methods (human height, chronotype), black; three methods (SCZ, BIP, cross-disorders, educational attainment, intelligence, cognitive performance).

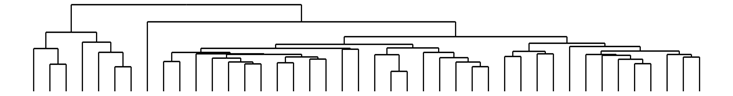

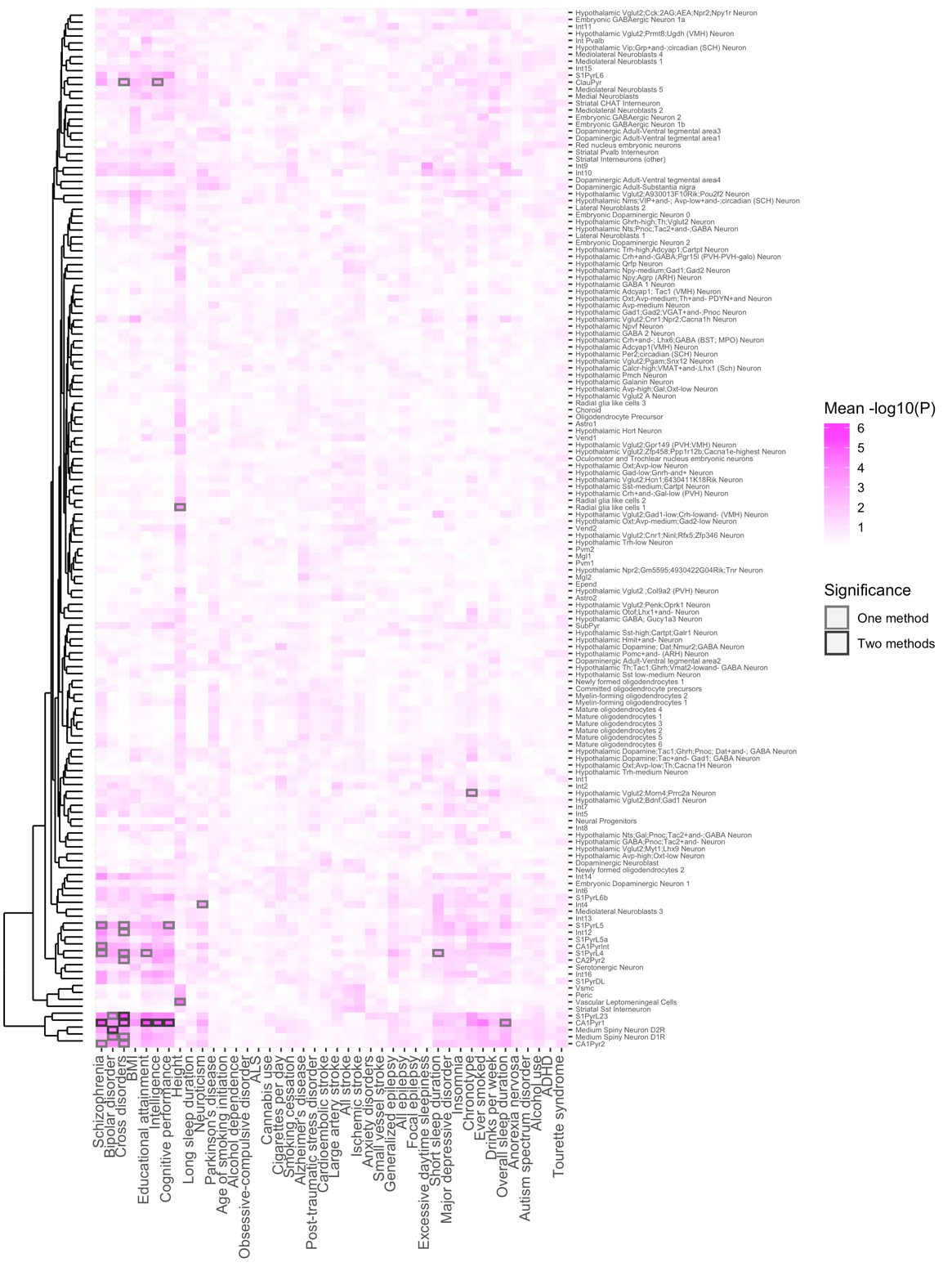

**Figure S22. Overview of enriched cell types of 42 common-variant psychiatric, neurologic and behavioral/quantitative GWAS results in the KI level 2 dataset, including MAGMA in linear mode.**

*Abbreviations: ADHD; attention deficit hyperactivity disorder, ALS; amyotrophic lateral sclerosis, BMI; body mass index.*

Analyses from LDSC, DEPICT, MAGMA (linear mode), MAGMA (top 10% mode) and MAGMA (top 10% mode) show mostly enrichment in various excitatory pyramidal cells and certain MSNs. Phenotypes and cell types are grouped by hierarchal clustering. Shades of pink are proportional to the mean strength of association (-log10(P)) of all methods. The colour of the frames relates to the number of methods that identified a given cell type as significant in a given phenotype, after Bonferroni correction (P<0.05/(149*42)). Frames: grey; one method (human height, neuroticism, short sleep duration, chronotype, overall sleep duration), black; two methods (SCZ, BIP, cross-disorders, educational attainment, intelligence, cognitive performance).

**Figure S23. Comparison of P-values computed by LDSC, MAGMA and DEPICT in the KI dataset.**

*Axes are -log10(P) transformed P-values from LDSC, MAGMA (top 10%), MAGMA (linear) and DEPICT. Dots represent cell types within the KI dataset. The blue line represents a linear regression and the area coloured in grey represents the standard error of the linear regression.*

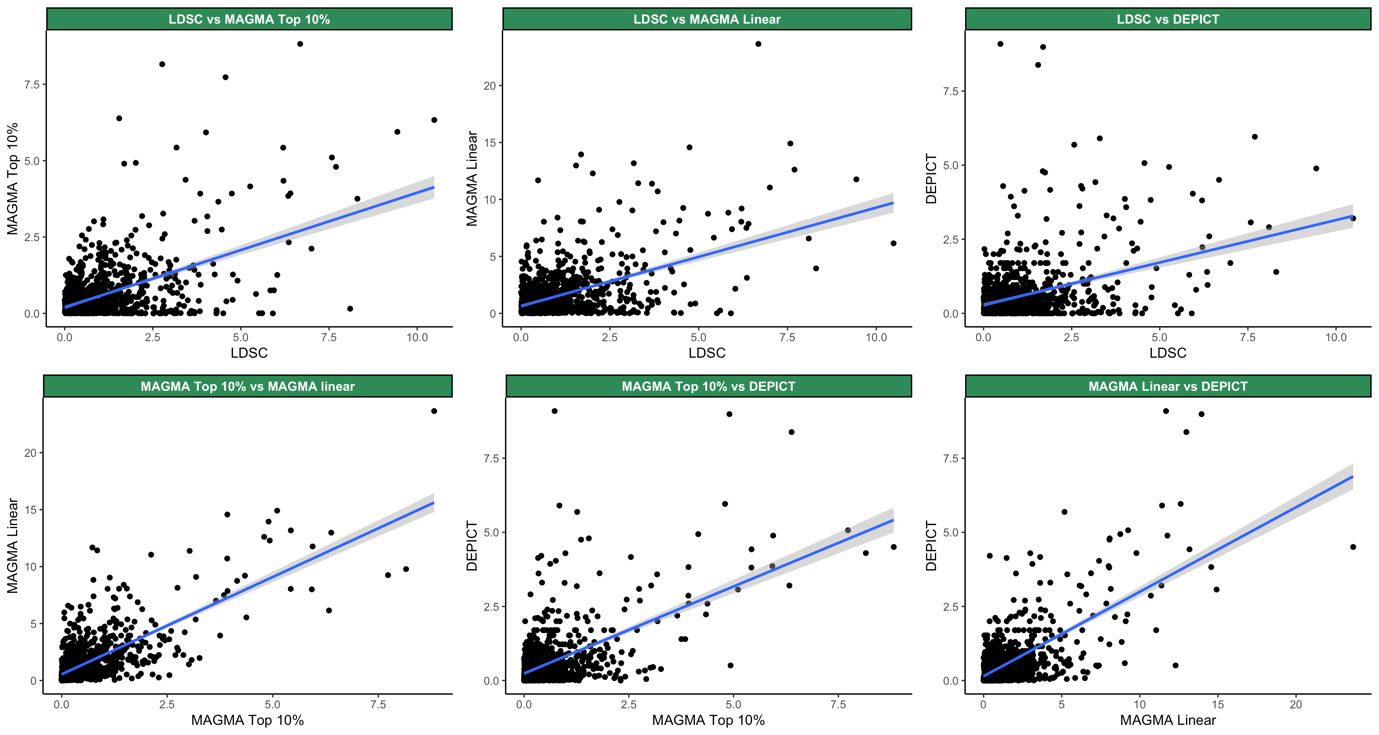

**Figure S24. Comparison of P-values computed by LDSC, MAGMA and DEPICT in the KI level 2 dataset.**

*Axes are -log10(P) transformed P-values from LDSC, MAGMA (top 10%), MAGMA (linear) and DEPICT. Dots represent cell types within the KI level 2 dataset. The blue line represents a linear regression and the area coloured in grey represents the standard error of the linear regression.*

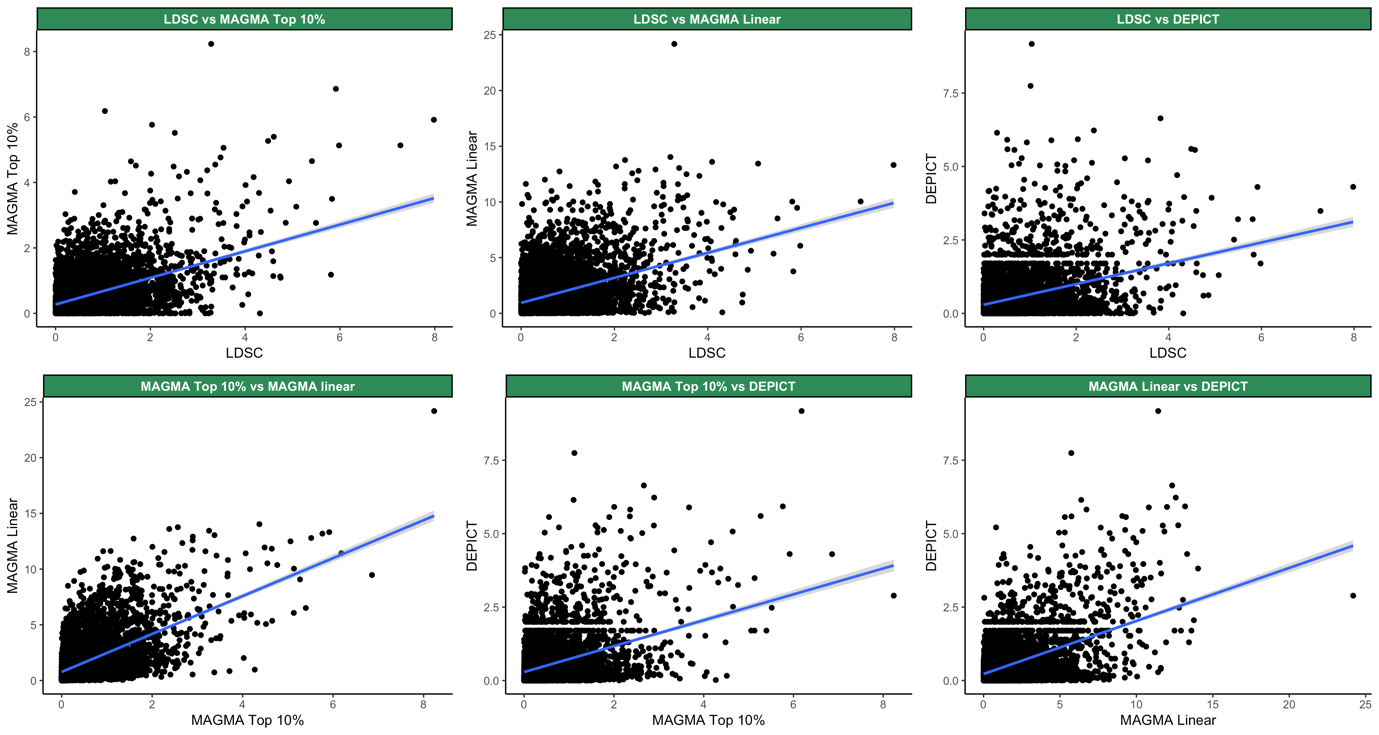

**Figure S25. Correlation between the top associated KI cell types with GWAS sample size.**

*Abbreviations: SUD; substance-use disorder.*

*The strength of association reflects the association of the top associated cell type with a given phenotype. The strongest association, out of LDSC, DEPICT, MAGMA (top 10% mode) and MAGMA (linear mode), was selected to denote the strength of association. The grey line represents a linear regression and the area coloured in a lighter shade of grey represents the standard error of the linear regression.* ***A.*** *Correlation between the top associated cell type, out of LDSC, DEPICT, MAGMA (top 10% mode) and MAGMA (linear mode), and sample size.* ***B.*** *Correlation between the top associated cell type, out of all methods excluding MAGMA (linear mode), and sample size.*

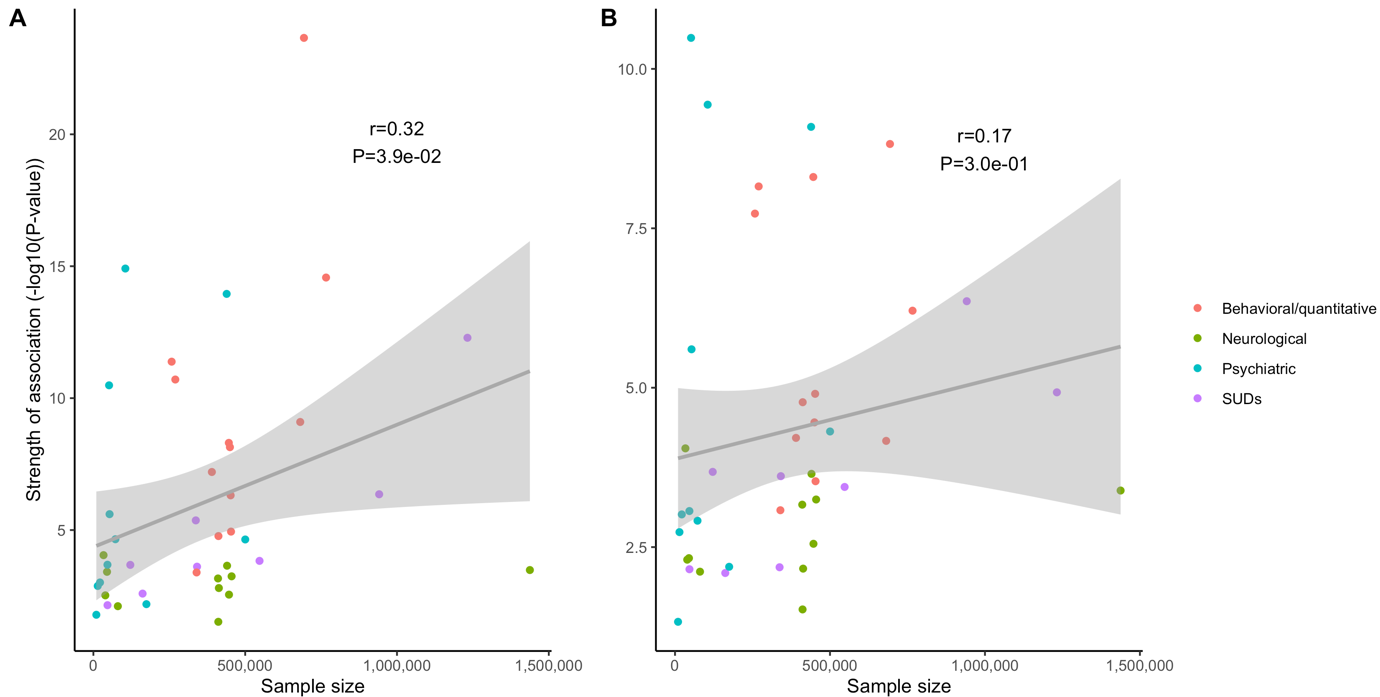

**Figure S26. Overview of confirmational cell type enrichment by FUMA.**

*Abbreviations: dLGN; dorsal lateral geniculate nucleus, CB; cerebellum, MB; midbrain, FC; frontal cortex, GP; globus pallidus, SN; substantia nigra, STR; striatum, PC; prefrontal cortex, SS; somatosensory cortex, DG; dentate gyrus, HC: hippocampus, iMSN/dMSN; (in)direct medium spiny neuron, D2; dopamine receptor2, MTG; middle temporal gyrus, SS; Somatosensory cortex,*

*Shades of pink are proportional to the strength of association (-log10(P). Areas are only shaded pink if a cell type is independently associated, after both within and across scRNA dataset conditional analysis, with a phenotype. Only phenotypes in which at least one cell type was enriched and only cell types that were implicated in at least one phenotype are shown.* ***A.*** *FUMA cell type enrichment results using human gene expression datasets.* ***B.*** *FUMA cell type enrichment results using mouse scRNA datasets.*

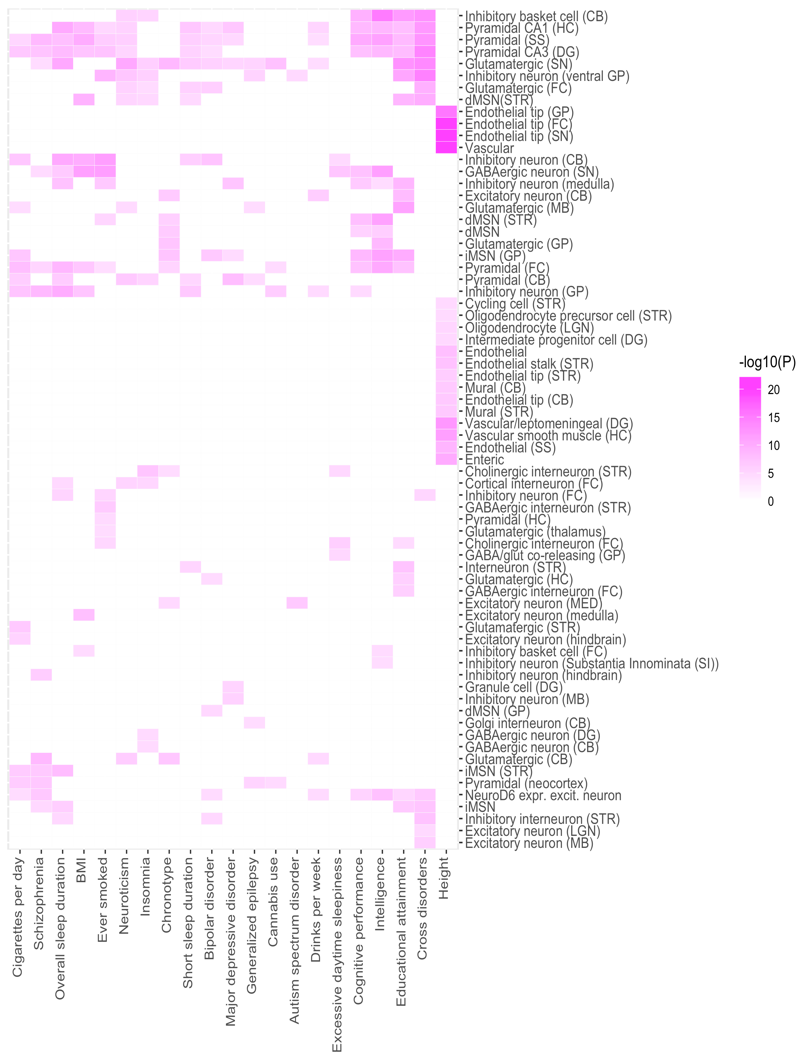

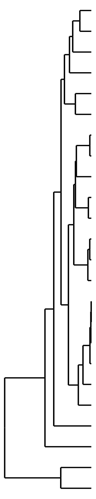

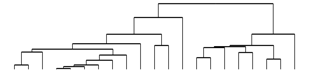

**B**

**A**

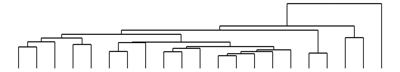

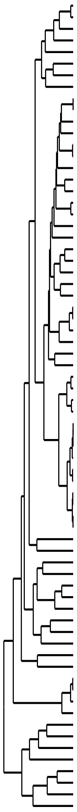

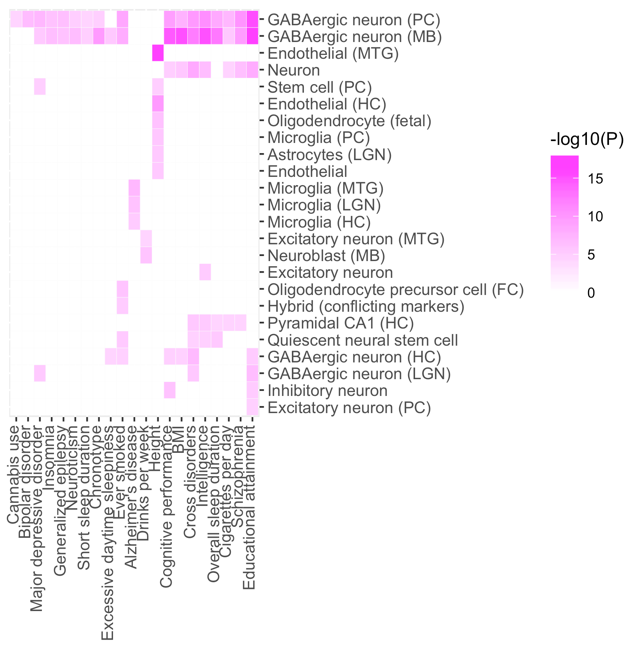

### **Supplementary Tables**

**Table S1. Dataset features.**

**Sample count of a phenotype that is part of larger group.*

|  | Trait | Reference | Cases | Controls | Samples | Ancestry |
| --- | --- | --- | --- | --- | --- | --- |
| Psychiatric disorders | Attention-deficit/hyperactivity disorder (ADHD) | Demontis, Walters et al., 2018^21^ | 19,099 | 34,194 |  | European |
|  | Anorexia nervosa (AN) | Watson, Yilmaz et al., 2019^22^ | 16,992 | 55,525 |  | European |
|  | Anxiety disorders | Otowa et al., 2016^23^ | 7,016 | 14,745 |  | European |
|  | Autism spectrum disorder (ASD) | Grove, Ripke et al., 2019^24^ | 18,382 | 27,969 |  | European |
|  | Bipolar disorder (BIP) | Stahl, Breen et al., 2019^25^ | 20,352 | 31,358 |  | European |
|  | Cross disorders | Cross-Disorder Group of the Psychiatric Genomics Consortium. Electronic address and Cross-Disorder Group of the Psychiatric Genomics 2019^26^ | 162,151 | 276,846 |  | European |
|  | Major depressive disorder (MDD) | Howard, Adams et al., 2019^27^ | 170,756 | 329,443 |  | European |
|  | Obsessive-compulsive disorder (OCD) | International Obsessive Compulsive Disorder Foundation Genetics Collaborative & O. C. D. Collaborative Genetics Association Studies, 2017^28^ | 2,688 | 7,037 |  | European |
|  | Post-traumatic stress disorder (PTSD) | Nievergelt, Mahofer et al., 2019^29^ | 23,212 | 151,447 |  | European |
|  | Schizophrenia (SCZ) | Pardiñas et al., 2018^30^ | 40,675 | 64,643 |  | European |
|  | Tourette syndrome (TS) | Yu et al., 2019^31^ | 4,819 | 9,488 |  | European |
| Substance use disorders | Alcohol use | Sanchez-Roige et al., 2019^32^ |  |  | 121,604 | European |
|  | Alcohol dependence | Walters et al., 2018^33^ | 11,569 | 34,999 |  | European |
|  | Drinks per week | Liu et al., 2019^34^ |  |  | 941,280 | European |
|  | Cannabis | Pasman et al., 2018^35^ |  |  | 162,082 | European |
|  | Age smoking initiation | Liu et al., 2019^34^ |  |  | 341,427 | European |
|  | Ever smoked regularly | Liu et al., 2019^34^ |  |  | 1,232,091 | European |
|  | Cigarettes per day | Liu et al., 2019^34^ |  |  | 337,334 | European |
|  | Smoking cessation | Liu et al., 2019^34^ |  |  | 547,219 | European |
| Neurological disorders | Amyotrophic lateral sclerosis (ALS) | Nicolas et al., 2018^36^ | 20,806 | 59,804 |  | European |
|  | Alzheimer's disease | Jansen et al., 2019^37^ | 71,880 | 383,378 |  | European |
|  | All epilepsy | The International League Against Epilepsy Consortium on Complex Epilepsies^38^ | 15,212 | 29,677 |  | Multi-ancestry |
|  | Generalized epilepsy | The International League Against Epilepsy Consortium on Complex Epilepsies^38^ | 3,769* | 29,677* |  | Multi-ancestry |
|  | Focal epilepsy | The International League Against Epilepsy Consortium on Complex Epilepsies^38^ | 9,671* | 29,677* |  | Multi-ancestry |
|  | Any stroke | Malik et al., 2018^39^ | 40,585 | 406,111* |  | European |
|  | Ischemic stroke | Malik et al., 2018^39^ | 34,217* | 406,111* |  | European |
|  | Large artery stroke | Malik et al., 2018^39^ | 4,373* | 406,111* |  | European |
|  | Cardioembolic stroke | Malik et al., 2018^39^ | 7,193* | 406,111* |  | European |
|  | Small vessel stroke | Malik et al., 2018^39^ | 5,386* | 406,111* |  | European |
|  | Parkinson’s disease | Nalls et al., 2020^40^ | 37,688 | 1,400,000 |  | European |
| Behavioral/  quantitative | Body mass index (BMI) | Yengo et al., 2018^41^ |  |  | 681,275 | European |
|  | Height | Yengo et al., 2018^41^ |  |  | 693,529 | European |
|  | Chronotype | Jones et al., 2019^42^ |  |  | 449,734 | European |
|  | Excessive daytime sleepiness | Wang et al., 2019^43^ |  |  | 452,071 | European |
|  | Sleep duration | Dashti et al., 2019^44^ |  |  | 446,118 | European |
|  | Short sleep duration | Dashti et al., 2019^44^ | 106,192* | 305,742* |  | European |
|  | Long sleep duration | Dashti et al., 2019^44^ | 34,184* | 305,742* |  | European |
|  | Insomnia | Lane et al., 2019^45^ |  |  | 453,379 | European |
|  | Intelligence | Savage et al., 2018^46^ |  |  | 269,867 | European |
|  | Educational attainment | Lee et al., 2018^47^ |  |  | 766,345 | European |
|  | Cognitive performance | Lee et al., 2018^47^ |  |  | 257,828 | European |
|  | Neuroticism | Nagel et al., 2018^48^ |  |  | 390,278 | European |

**Table S2. FUMA names of mouse and human brain scRNA-seq datasets employed for FUMA analyses.**

*Abbreviations: LGd/LGN; lateral geniculate nucleus, CB; cerebellum, FC; frontal cortex, GP; globus pallidus, SN; substantia nigra, STR; striatum, MTG; middle temporal gyrus.*

| Mouse | Human |
| --- | --- |
| Allen_Mouse_LGd2_level3^49^ | PsychENCODE_Developmental^50^ |
| DropViz_CB_level2^51^ | Allen_Human_LGN_level2**^49^** |
| DropViz_FC_level2^51^ | DroNc_Human_Hippocampus^52^ |
| DropViz_GP_level2^51^ | GSE104276_Human_Prefrontal_cortex_per_ages^53^ |
| DropViz_SN_level2^51^ | GSE67835_Human_Cortex^54^ |
| DropViz_STR_level2^51^ | Linnarsson_GSE76381_Human_Midbrain^10^ |
| Linnarsson_GSE101601_Mouse_Somatosensory_cortex^55^ | Allen_Human_MTG_level2**^49^** |
| Linnarsson_GSE104323_Mouse_Dentate_gyrus^56^ |  |
| Linnarsson_GSE60361_Mouse_Cortex_Hippocampus_level2^8^ |  |
| Linnarsson_MouseBrainAtlas_level5^57^ |  |
| GSE97478_Mouse_Striatum_Cortex^58^ |  |

**Table S3. Marker genes for brain cell types in mice described in literature.**

| Gene | Cell type |
| --- | --- |
| From Zeisel et al.^8^ | |
| Tbr1 | Excitatory neurons |
| Olig1 | Oligodendrocytes |
| Aldoc | Astrocytes |
| Gfap | Astrocytes |
| Ly6c1 | Endothelial cells |
| Pnoc | Interneurons |
| Gad1 | Inhibitory neurons / Interneurons |
| From Tasic et al.^15^ | |
| Snap25 | Pan-neuronal |
| Gad1 | Inhibitory / GABAergic neurons |
| Gad2 | Inhibitory / GABAergic neurons |
| Vip | Inhibitory / GABAergic neurons |
| Sst | Inhibitory / GABAergic neurons |
| Pvalb | Inhibitory / GABAergic neurons |
| Slc17a7 | Excitatory / glutamatergic neurons |
| Aqp4 | Astrocytes |
| Mog | Oligodendrocytes |
| Itgam | Microglia |
| Flt1 | Endothelial cells |
| From Artegiani et al.^14^ | |
| Aldoc | Neural stem cells / Astrocytes |
| Eomes | Neural progenitors |
| Cspg4 | Oligodendrocyte precursors |
| Pdgfra | Oligodendrocyte precursors |
| Neurod1 | Neural progenitors |
| Sox9 | Neural stem cells / Astrocytes |
| Id4 | Neural stem cells / Astrocytes |
| Csf1r | Microglia |
| Cx3cr1 | Microglia |
| Olig1 | Oligodendrocyte precursors |
| Olig2 | Oligodendrocyte precursors |
| Sox10 | Oligodendrocyte precursors |
| Gfap | Neural stem cells / Astrocytes |
| Ndnf | Interneurons |
| Vwf | Endothelial cells |
| From 10x Genomics^11^ | |
| Stmn2 | Pan-neuronal |
| Tbr1 | Excitatory neurons |
| Gad1 | Inhibitory neurons |
| Hes1 | Glial cells |
| Aldoc | Astrocytes |
| Olig1 | Oligodendrocytes |
| From Bhaduri et al.^12^ | |
| Igfbp7 | Endothelial cells |
| Neurod6 | Neurons |
| Dlx2 | Interneurons |
| Dlx6os1 | Interneurons |
| Eomes | Intermediate progenitors |
| Vim | Radial glial cells |
| Reln | Cajal-Retzius cells |
| Olig1 | Oligodendrocytes |
| Aqp4 | Astroependymal cells |
| Igfbpl1 | Neuroblasts |
| Eomes | Neuroblasts |
| Riiad1 | Radial glial cells |
| Ascl1 | Progenitors |
| Slc17a7 | Glutamatergic neurons |
| Slc32a1 | GABEergic neurons |

**Table S4. Cell type identities assigned to clusters (k-means clustering, k=20) for the 10x Genomics dataset and their lines of evidence.**

Marker gene detection consisted of a Wilcoxon rank-sum test with Benjamini-Hochberg correction to select top-ranked genes for specificity to cell clusters. Clusters 12, 18, 19 and 20 were not included as they were removed during quality control earlier on. The column ‘Lines of evidence’ sums up the specific genes that were used to assign cell type identities.

Abbreviations: Lit.; matches with marker gene from literature, Wiki; matches with marker gene-/primarily expressed in cell type from companion wiki (<http://mousebrain.org/>), Expr.; shows exclusive expression of marker gene from literature (heat map not shown here).

|  |  | Top 10 marker gene detection results | | | | | | | | | |  |
| --- | --- | --- | --- | --- | --- | --- | --- | --- | --- | --- | --- | --- |
| Cluster | **Assigned identity** | **Gene #1** | **Gene #2** | **Gene #3** | **Gene #4** | **Gene #5** | **Gene #6** | **Gene #7** | **Gene #8** | **Gene #9** | **Gene #10** | **Lines of evidence** |
| 1 | Neurons (glutama-tergic) | Gria2 | Neurod2 | Gpm6a | Tsc22d1 | Cd24a | Tuba1a | Neurod6 | Nrp1 | Stmn4 | Ttc28 | - Lit. (Neurod6)  - Wiki (Gria2) |
| 2 | Neuro-blasts (1) | Igfbpl1 | Pcp4 | Sox11 | Ppp2r2b | Rnd2 | Tagln3 | 6330403K07Rik | Nfib | Unc5d | Sstr2 | - Lit. (Igfbpl1)  - Wiki (Igfbpl1, Sox11) |
| 3 | Neuro-blasts (2) | Islr2 | Mapt | Thra | Snca | Stmn2 | Rtn1 | Map1b | Zbtb20 | Tmsb10 | Fabp5 | - Lit. (Stmn2)  - Wiki (Islr2) |
| 4 | Neurons | Gria2 | Arpp21 | Neurod6 | Ptprd | Opcml | Satb2 | Mef2c | Robo2 | Ntm | Tmem108 | - Lit. (Neurod6)  - Wiki (Gria2) |
| 5 | Inter-neurons (1) | Nrxn3 | Maf | Meg3 | Mef2c | Nxph1 | Arx | Lhx6 | Dlx2 | Gad2 | Dlx1 | - Lit. (Dlx2) |
| 6 | Inter-mediate progenitors | Mdk | Neurog2 | Eomes | Rps18 | Dbi | Rps9 | Rpl13a | Rpl32 | Rpl18a | Rps6 | - Lit. (Eomes) |
| 7 | Inter-neurons (2) | Dlx6os1 | Nnat | Sp9 | Gad2 | Meg3 | Dlx2 | Nrxn3 | Dlx5 | Dlx1 | Bcl11b | - Lit. (Dlx6os1, Dlx2) |
| 8 | Astrocytes (1) | Dbi | Mt3 | Fabp7 | Aldoc | Slc1a3 | Phgdh | Vim | Ptn | Cst3 | Ddah1 | - Lit. (Aldoc), Wiki (Dbi) |
| 9 | Enteric neurons | 2810417H13Rik | H2afz | Hmgb2 | Rrm2 | Dut | Anp32b | Ranbp1 | Hmgn2 | Tuba1b | Dek | - Wiki (2810417H13Rik, Rrm2) |
| 10 | Enteric glial cells | Cks2 | Hmgb2 | Cenpa | Ube2c | Cenpf | H2afz | Birc5 | Cdca8 | Cdca3 | Cdc20 | - Wiki (Cks2, Cenpf, Ube2c, Cdc20) |
| 11 | Cajal-Retzius cells | Reln | Nhlh2 | Pcp4 | Gap43 | Snhg11 | Meg3 | Ndnf | Cacna2d2 | Ly6h | Cd200 | - Lit. (Reln) |
| 13 | Endothelial cells | Cldn5 | Itm2a | Igfbp7 | Sparc | Ramp2 | Bsg | Slc7a5 | Egfl7 | Vwa1 | Slc2a1 | - Lit. (Igfbp7)  - Wiki (Cldn5) |
| 14 | Oligoden-drocytes | Olig1 | Olig2 | Serpine2 | Fabp7 | Pdgfra | Cspg5 | Ptprz1 | Bcan | Ckb | Ncald | - Lit. (Olig1, Olig2) |
| 15 | Vascular endothelial cells | Igf2 | Vtn | Eva1b | Cald1 | Ndufa4l2 | Col4a1 | Gng11 | Atp1a2 | Higd1b | Igfbp7 | - Wiki (Igf2) |
| 16 | Microglia | Tyrobp | Ctsd | C1qc | C1qb | C1qa | B2m | Fcer1g | Lgmn | Apoe | Sepp1 | - Wiki (Tyrobp, C1qc, C1qb, C1ga)  - Expr. (Csf1r, Cx3cr1) |
| 17 | Astrocytes (2) | Clu | Dbi | Apoe | Sparc | Mt3 | Ttyh1 | Tmem47 | Id4 | Fabp7 | Vim | - Lit. (Apoe, Id4)  - Wiki (Clu, Apoe, Id4) |
